# How to map a plantain: phylogeny of the diverse *Plantagineae* (Lamiales)

**DOI:** 10.1101/2020.07.31.230813

**Authors:** Alexey Shipunov, José Luis Fernández A., Gustavo Hassemer, Sean Alp, Hye Ji Lee, Kyle Pay

## Abstract

The tribe *Plantagineae* (Lamiales) is a group of plants with worldwide distribution, notorious for its complicated taxonomy, still unresolved natural history, and a trend of morphologic reduction and simplification. This tribe includes the plantains (*Plantago*), the small aquatic *Littorella*, and the northern Andean shrubs *Aragoa*. Some *Plantago* lineages exhibit remarkably high diversification rates, which further adds to the complicated classification, and the worldwide distribution of these plants raises numerous questions related to vicariance and dispersal. In this work, we present the broadest phylogeny of the group to date and discuss the evolutionary, morphological, and biogeographical implications of our phylogenetic results, including the description of two new species from the Americas.

## INTRODUCTION

The tribe *Plantagineae* (Lamiales) consists of well-known, worldwide distributed plantains (*Plantago*), small aquatic *Littorella*, and Andean (mostly Colombian) páramo shrubs *Aragoa*. This group of plants is notorious for its complicated taxonomy, still unresolved natural history and a trend of morphologic reduction and simplification (Fernández, 1995; Rahn, 1996; Rønsted el al., 2002; Hassemer et al., 2019). Therefore, the elucidation of the phylogenetic relationships within this tribe is of high importance not only for taxonomic classification but also for phytogeography and our general understanding of plant evolution.

Plantains (ribworts), *Plantago* L., are remarkable plants. They grow almost everywhere all around the world, except for the Antarctic and tropical wet forests (Rahn, 1996; Hassemer et al., 2016). Morphologically, they bear the unusual combination of characters (Linnaeus, 1754): sympetalous 4- merous non-showy flowers developing into circumscissile capsule-like fruit (pyxidium), and monocot- like leaves with arcuate or parallel venation, usually borne in a rosette. Even in times when only sixteen species were described in *Plantago*, botanists mentioned a remarkable similarity between different species (Linnaeus, 1753); this typically results in a significant amount of incorrectly determined specimens, even in leading herbarium collections. Plantains exhibit some of the terminal stages of morphological reduction among Lamiales, and recent research (Preston et al., 2011) demonstrated how this flower reduction could happen within *Antirrhinum*-*Plantago* lineage. There is also a reduction of vegetative characters expressed in many plantains, and only a few species have a branched stem and “dicotyledonous” leaves. In all, evolution towards anemophily resulted in significant morphological convergence with graminoid monocots.

Geographic distribution of *Plantago* is exceptionally broad, and this is due not only to cosmopolitan weeds *P. lanceolata* L. and *P. major* L. On many remote ocean islands, for example, there are unique species of *Plantago*, and some of them exhibit a remarkable tendency to evolve into woody plants (Carlquist, 1970; Iwanycki Ahlstrand et al., 2019). As a whole, *Plantago* likely underwent a rapid (Cho et al., 2004) evolution and recent (Meudt et al., 2015) diversification. Many new *Plantago* species have recently been discovered and described, most of them rare and narrowly distributed (e.g., Hassemer and Rønsted, 2016; Hassemer et al., 2018a; Hassemer 2019). All of the above makes *Plantago* a group of plants with considerable interest for biodiversity conservation (Hassemer et al., 2016). Furthermore, *Plantago* is a complicated genus also from a nomenclatural point of view (Di Pietro et al., 2013; Doweld and Shipunov, 2017; Hassemer, 2018a, 2018b), and many problems remain to be solved.

Aquatic *Littorella* P.J.Bergius (Bergius, 1768) comprises three species with considerably disjunct distribution; they grow in shallow waters of North America Great Lakes, Patagonia, and Northern Europe (here both in lakes and desalinated North and Baltic seas). These plants are morphologically similar to *Plantago*, and Rahn (1996) joined two genera. However, when molecular data started to be available (Hoggard et al., 2003), *Plantago* and *Littorella* were shown as sister clades, and since then nearly all authors preferred to keep the tradition of recognizing the two genera as separate.

Contrary to *Plantago* and *Littorella*, woody shrubs *Aragoa* Kunth (Kunth, 1818) are the local endemics of páramo in Colombia (one species also in Venezuela). Until the first molecular results (Bello et al., 2002), it was never considered to be a sister group for *Plantago* + *Littorella* clade but rather unplaced in “old” Scrophulariaceae (Fernández, 1995). *Aragoa* is relatively speciose, containing more than 20 described species and also several hybrids (Fernández, 1993, 1995), whereas no hybrid species have been described in two other genera. The hybridization might explain the rapid radiation and speciation in this genus (Fernández, 2002). Flowers of *Aragoa* are likely animal-pollinated (Fernández, 1995) but actinomorphic, and leaves are reduced, similarly to two other genera of the group (Bello et al., 2004).

*Plantago*, *Littorella* and *Aragoa* form well-supported (both morphologically and molecularly), stable clade (Bello et al., 2002; Rønsted et al., 2002; Bello et al., 2004; Meudt et al., 2015) which we call hereafter the tribe *Plantagineae* Dumort. (Dumortier, 1829). Multiple detailed morphology-based works were published about *Plantagineae*; most important are Barnéoud (1845), Decaisne (1852), Pilger (1937), Fernández (1995), and Rahn (1996). However, the comprehensive molecular study based on the broad sampling of this whole group is still absent. The broadest at the moment are works of Rønsted et al. (2002) and Hoggard et al. (2003), which included 59 and 27 species, respectively (whereas the group is estimated to include ca. 250 species). To compare further, the GenBank database as of June 2019 contains only about 140 species entries (and this does not account for possible synonymy). This situation has shown signs of improvement lately, and several publications which cover the complicated *Plantago* subg. *Plantago* (e.g., Hassemer et al., 2019; Iwanycki Ahlstrand et al., 2019) and subg. *Coronopus* (Höpke et al., 2019; Hassemer et al., 2017; Hassemer 2018b) are now available. Some recent regional works (Tay et al., 2010; Meudt, 2011, 2012) also improved our knowledge of subg. *Plantago*. However, no recent molecular works are focusing on *Aragoa* diversity, and since Rønsted et al. (2002), nothing significant was published about molecular taxonomy of *Psyllium* and allies (we include here subgenera *Bougueria*, *Psyllium* s.str. and *Albicans*).

Therefore, we consider *Plantagineae* the under-studied group, especially in the molecular aspect. This situation dictates the necessity of the molecular phylogenetic study with the broadest sampling in mind. We continue the line of Rønsted et al. (2002) and employ similar barcoding markers but aim for the greater species coverage, with the ultimate goal to assess all described species of the group and obtain the most detailed picture of their relationships. Together with molecular characters, we employ morphological characters identified in Rahn (1996) and morphometric characters from samples used in molecular studies. Our goal was also to solve multiple problems with the *Plantagineae* taxonomy and geography, whose resolution will improve the overall understanding, help in conservation, ecology, and invasive biology studies, and will ease the identification of *Plantagineae* species which still is a tedious and difficult task, especially for beginners.

## MATERIAL AND METHODS

### PLANT MATERIAL

It is virtually impossible to sample 250 species (Support Table 1) without the help of the herbarium collections. *Plantagineae* is infrequently cultivated, and re-collection from nature, especially herbaceous short-living plants, is a task that is successful only rarely (e.g., Hassemer et al., 2018a). Therefore, while some of our samples were collected into silica gel from the living plants, the majority of work (94%) involved tissue samples taken (with the kind permission of herbarium curators) from plants collected years ago.

**Table 1.** Machine learned placements of the molecularly unsampled species.

Using herbarium samples poses some restrictions. While the purity and concentration of DNA do not significantly suffer from time (Choi et al., 2015), the quality of sequences heavily depends on sample age, collection methods, and the nature of fragment to amplify. With older sample age, more difficult drying process, and longer fragment, our chances to obtain the useful data were significantly lower.

Since not every sample yields reliable DNA, we typically collected multiple samples per species and attempted to extract and sequence it numerous times. Now we have sequences of at least one DNA marker from 220 species (including 192 *Plantago* species). Data from 86 species have been taken from public databases. In all, we were able to increase the amount of available information three-fold (four-fold in Aragoa). Due to the apparent problems with identification (Funk et al., 2018), we always trusted our samples first.

### DNA SEQUENCING

DNA extraction performed using multiple standard protocols, but soon after the start of the project (2011), we decided to stay with NUCLEOSPIN Plant II Kit (MACHEREY-NAGEL GmbH & Co. KG, Düren, Germany) which seems to be is a good trade-off between efficiency and simplicity. We improved this protocol in several points, e.g., increased the lysis time to 30 min and used thermomixer on the slow rotation speed (350 rpm) instead of a water bath. To assess DNA quality, we used Nanodrop 1000 Spectrophotometer (Thermo Scientific, Wilmington, DE, USA), which estimates concentration and purity (the 260/280 nm ratio of absorbance) of samples. Typically, 1.4 ratio was enough to guarantee PCR amplification of smaller markers, whereas the ratio 1.7 was an average in the group. The lower DNA quality was typically obtained from samples presumably collected in wet climates.

Especially low was the quality of our *Aragoa* samples; this might be due to the widespread use of drying cabinets, the theory which indirectly supported with good results which we obtained from samples dried without any help (even without silica gel), just in room conditions in Bogotá. Reversely, we sometimes were able to extract, amplify, and sequence samples collected long ago, e.g., from *Plantago sinaica* Decne. collected in 1834!

Nevertheless, short barcoding DNA markers (Kuzmina and Ivanova, 2011) are the best to amplify for herbarium samples; therefore, our first choice was nuclear ITS2 and chloroplast *trn*L-F spacer and *rbc*L gene. We amplified them following the Barcoding of Life recommendations and protocols (Kuzmina and Ivanova, 2011). Several samples were sequenced with the direct help of Barcoding of Life (“SAPNA” project); this last project provided us also with sequences of mitochondrial COI and plastid *mat*K markers.

Typically, our PCR the reaction mixture had a total volume of 20 μL which contained 5.2 μL of PCR Master Mix (components mostly from Thermo Fisher Scientific, Waltham, Massachusetts supplied with Platinum DNA Taq Polymerase), 1 μL of 10 μM forward and reverse primers, 2 μL of DNA solution from the extraction and 10.8 μL of MQ purified water (obtained from a Barnstead GenPure Pro system, Thermo Scientific, Langenselbold, Germany) in the TBT-PAR water mix (Samarakoon et al., 2013). The latter was developed to improve amplification from the herbarium samples. Thermal cycler programs were mostly 94 °C for 5 min, then 35 cycles of 94 °C for 1 min; 50–52 °C (depending on the primer) for 1 min, 72 °C for 2 min, and finally 72 °C for 10 min. PCR products were sent for purification and sequencing to Functional Biosciences, Inc. (Madison, Wisconsin) and sequenced there under standard Sanger-based protocol. Sequences were obtained, assembled, and edited using Sequencher™ 4.5 (Genes Codes Corporation, Ann Arbor, Michigan, USA).

### PHYLOGENETIC ANALYSES

Subsequent steps use the “Ripeline” workflow. This workflow is the collection of UNIX shell and R (R Core Team, 2019) scripts that automate steps related to sequence selection, quality checking, alignments, gap coding, concatenation, and phylogenetic tree production. Ripeline involves multiple pieces of software, for example, AliView (Larsson, 2014), MUSCLE (Edgar, 2004), APE (Paradis et al., 2004), MrBayes (Ronquist and Huelsenbeck, 2003), ips (Heibl, 2008), shipunov (Shipunov et al., 2019), and phangorn (Schliep, 2011).

With the help of Ripeline, we were able to obtain maximal parsimony (MP) and Bayesian (MB) phylogenetic trees. MP analyses run with the help of the R phangorn package (Schliep, 2011) using parsimony ratchet (Nixon, 1999) with 2000 iterations and then 1000 bootstrap replicates. MB analyses run through the combination of MrBayes 3.2.6, and R ips and shipunov packages (Ronquist and Huelsenbeck, 2003; Heibl, 2008; Shipunov, 2019b). MCMC analysis (2 runs, 4–8 chains) was run for 1,000,000 generations, sampling every 10th generation resulting in 100,000 trees, and checked for the convergence. The first 25% of trees discarded as burn-in and the remaining trees summed to calculate the posterior probabilities. With the aid of R ape package (Paradis et al., 2004), all trees rooted with *Veronicastrum virginicum* (L.) Farw. and *Tetranema roseum* (M.Martens & Galeotti) (= *Tetranema mexicanum* Benth.) as outgroups, or with the *V. virginicum* alone.

Our phylogenetic trees use two data sets. Since all single marker trees were concordant, we used (with the help of Ripeline) the super-matrix approach. Our first data set included multiple barcoding markers available from public databases and our sequencing, which is chloroplast *rbc*L, *trn*L-F, *mat*K, and also nuclear ITS and mitochondrial COI. We call this dataset “broad” since it is relatively rich in data but has a limited sampling along the species dimension (87 entries and 4188 bp, including 656/497/561/1565/909 bp in COI, ITS2, *rbc*L, *trn*L-F and *mat*K, respectively). The second dataset was made with the broadest species coverage but included only ITS2 and *trn*L-F data, which is originated mostly from our sequencing efforts. Below, we designate it as “tall” (273 entries, including some subspecies and forms and 2062 bp length, including the same ITS and *trn*L-F fragment lengths as above).

### MORPHOMETRICS

Ripeline is also capable of using morphological characters, and we employed the updated and expanded morphological dataset from Rahn (1996) to make combined (molecules + morphology) and pure morphological datasets. To make Rahn’s (1996) dataset digital, we OCR’ed and cleaned the text, tables from it were converted into spreadsheets and merged.

To emphasize the weight of morphological characters, we used the hyper-matrix approach (Ashkenazy et al., 2018) and multiplied morphological dataset several times in order to achieve the approximate equality between numbers of molecular and morphological characters. We added characters of seed sculpture (Shipunov, 1998a, 1998b; Shehata and Loutfy, 2006) to the characters used in Rahn (1996), and expanded the dataset with species absent in the last work. In total, our binary morphological matrix has 114 characters and 271 entries.

We also were able to use the measurements of seven most apparent morphometric characters of *Plantago*: petiole, leaf, spike, and scape lengths, maximal leaf width, presence of taproot, and looseness (“gaps”) in the inflorescence. In total, we measured these characters on 405 herbarium samples (same which were in DNA extraction).

Using morphological, morphometric and DNA datasets, we were able to perform the broad spectrum of statistical analyses, including Procrustes analysis of the correspondence between molecular and morphological information (Peres-Neto and Jackson, 2001; Balbuena et al., 2013) and nearest neighbor machine learning (Ripley, 1996) for the placement of under-studied taxa. As an additional source to use in the placement process, we employed the combined molecular + morphology phylogenic tree.

We also employed the recursive partitioning (Venables and Ripley, 2002; Höpke et al., 2019), the machine learning technique which takes the classification and creates binary trees for the rest of data set. The structure of these trees is similar to the dichotomous keys (Therneau et al., 2014). Naturally, results of recursive partitioning are applicable for the construction of the dichotomous keys, which could help in the discrimination of *Plantago* sections and species.

Datasets, scripts, and phylogenic trees used in the preparation of this publication are available from the first author’s Open Repository here: http://ashipunov.info/shipunov/open/plantago.zip. Ripeline is available on Github: https://github.com/ashipunov/Ripeline. We encourage readers to reproduce our results and develop our methods further. All sequences were deposited into the GenBank.

In the paper, we followed the “appropriate citation of taxonomy” (ACT) principle (Seifert et al., 2008) and cited names of the most supra-species groups (Reveal, 2012).

## RESULTS

### PLANTAGINEAE IN GENERAL

All trees based on “broad” and “tall” datasets returned the stable (*Aragoa*, (*Littorella*, *Plantago*)) topology (Fig. 1), typically with the longest branch leading to *Aragoa*. As our maximum parsimony (MP) trees do not differ significantly from Bayesian (MB) trees, we hereafter present the results (Fig. 1–5) based mostly on the second type of analysis.

**Figure 1.**
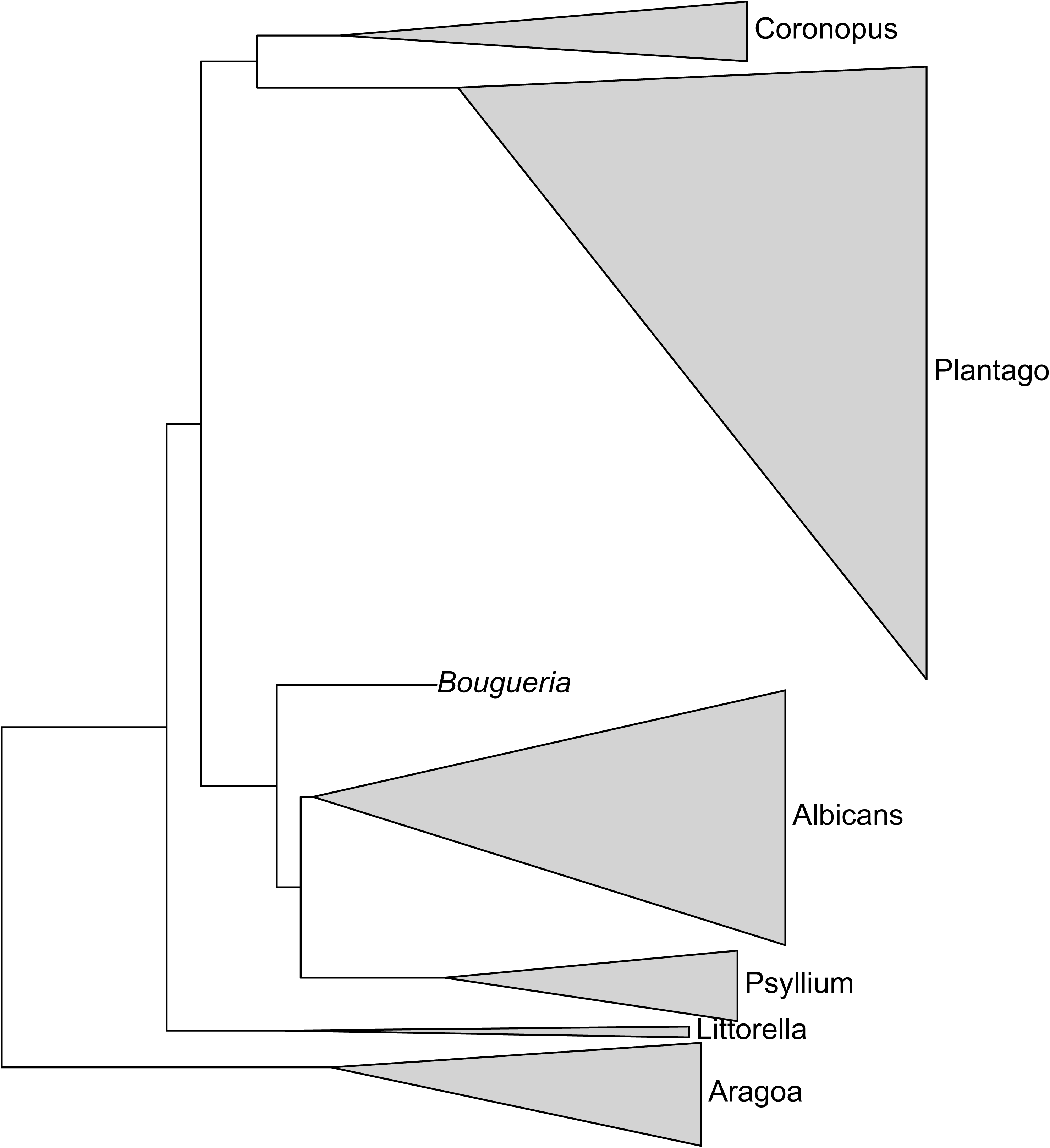
Phylogeny of the *Plantagineae*: the general arrangement of clades (genera and subgenera). Each triangle is the result of concatenation applied to the branches of the Bayesian (MB) phylogenetic tree.

### ARAGOA

Generally, the stability of subclades is not high in *Aragoa* (Fig. 2). The most stable is a placement of *Aragoa lucidula* S.F.Blake as a sister to all other studied *Aragoa* species. Within the rest of the subtree, the majority of species make one clade with subdivisions mostly without high support.

**Figure 2.**
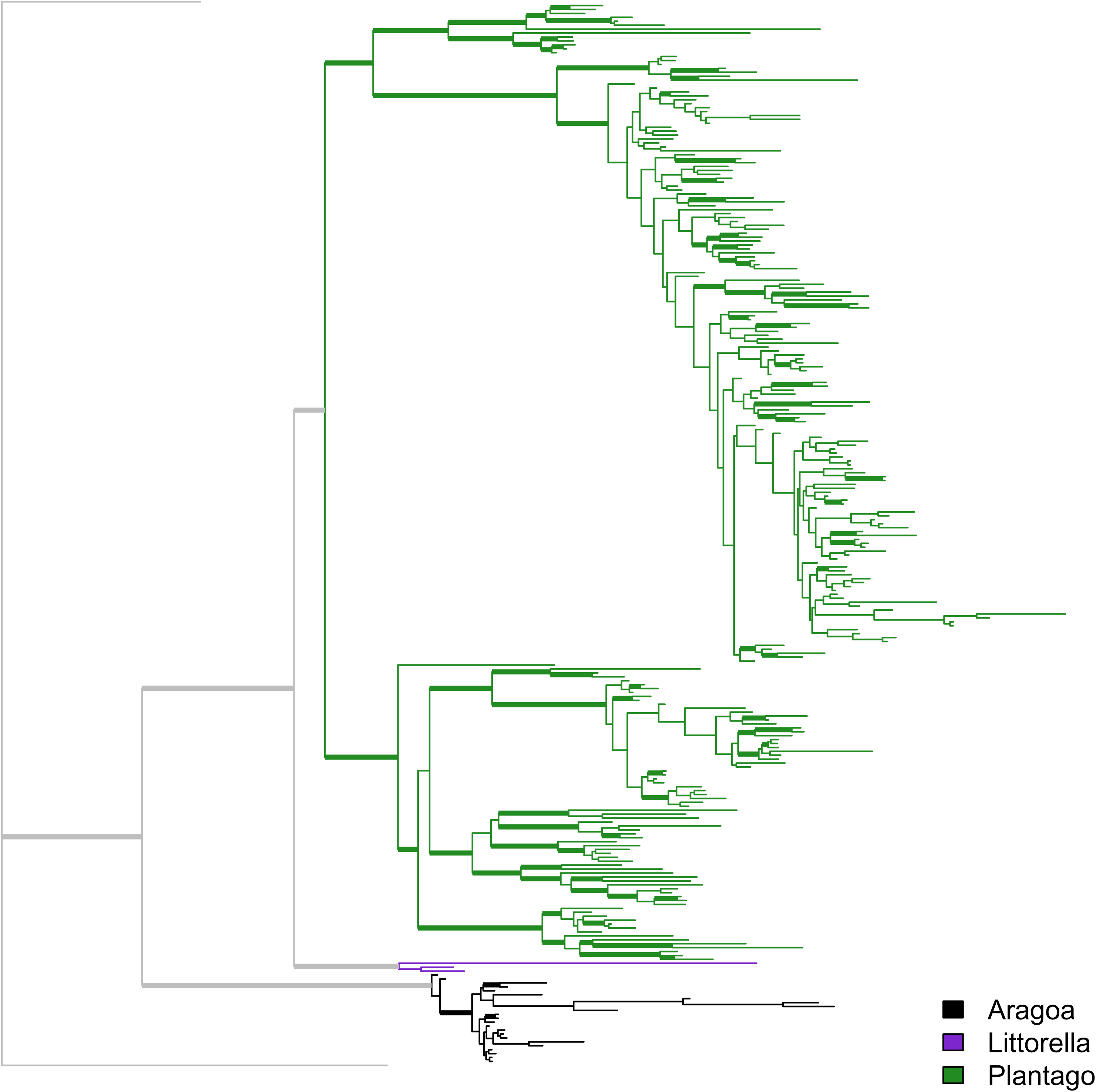
The overview of the MB tree based on the “tall” dataset. Branches with support > 90% thickened.

Morphologically outstanding *A. dugandii* Romero forms the clade with *A. lycopodioides* Benth. and *A. occidentalis* (all branches here are significantly longer than in other parts of *Aragoa* tree). The other unusual species, *A. perez-arbelaeziana* Romero, forms a clade with *A. romeroi* Fern.Alonso (Fig. 4, 5A).

### LITTORELLA

Three (two in the “broad” dataset) species of *Littorella* make the stable group where European *L. uniflora* (L.) Asch. is sister to American *L. americana* Fernald and *L. australis* Griseb. ex Benth. & Hook.f.

### *PLANTAGO* IN GENERAL

There is relatively high support for three major subdivisions of *Plantago*, which correspond with subgeneric rank. The topology is robustly (*Psyllium* s.l., (*Coronopus*, *Plantago*)), or in more detail, ((*Bougueria*,(*Psyllium* s.str., *Albicans*)), (*Coronopus*, *Plantago*)). Subgenera *Plantago*, *Coronopus*, and *Psyllium* clade form the remarkable “three-ridge” phylogenetic density surface (Fig. 3).

**Figure 3.**
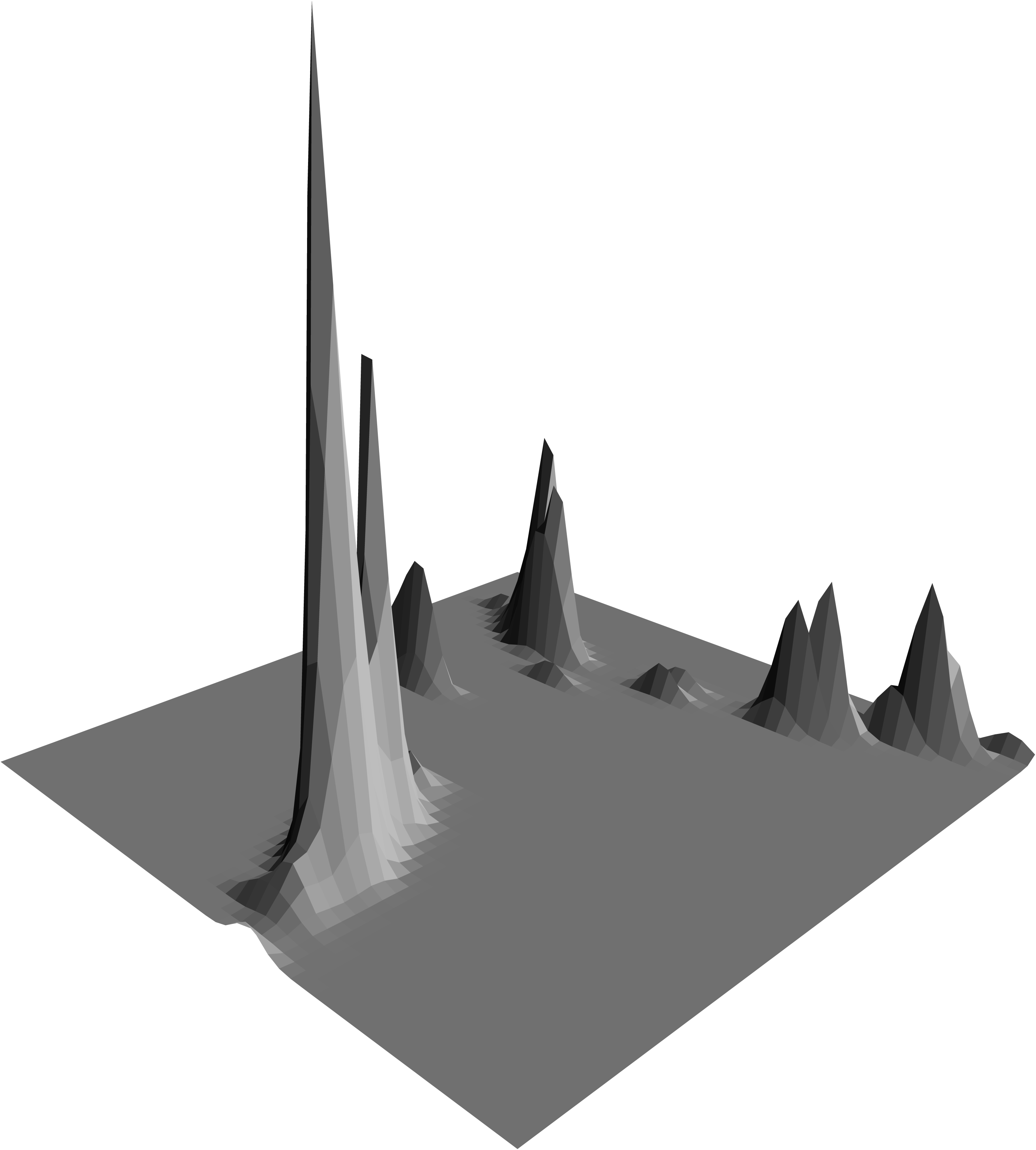
Density surface of the cophenetic space, based on the MB trees from the “tall” dataset. This surface is the result of multidimensional scaling of the cophenetic distances between tree tips; density of points estimated with 2D binned kernel. “Ridges” reflect areas with the highest density and correspond well with three major subgeneric divisions of *Plantago*. Note the three-fold structure of the “phylogenetic surface”: tallest corresponds with subg. *Plantago*, close and behind it, is subg. *Coronopus* whereas subg. *Psyllium* and allies form the rightmost “ridge.”

### PLANTAGO SUBG. PLANTAGO

Only trees originated from the “broad” dataset have relatively high support for clades within this group, whereas “tall” trees have the reliable support only for some terminal clades (Fig. 2, 4, 5B-D).One of the most stable groups consists of *P. media* L., *P. canescens* Adams, *P. arachnoidea* Schrenk ex Fisch. & C.A.Mey., *P. krascheninnikovii* C.Serg., *P. maxima* Juss. ex Jacq., *P. perssonii* Pilg. and *P. schwarzenbergiana* Schur. Tetraploid, xeromorph variant of *P. media* described as *P. urvillei* Opiz (*P. media* subsp. *stepposa* (Kuprian.) Soó), typically does not branch with *P. media* s.str.

*Plantago krascheninnikovii* from the Urals is habitually similar to the inland forms of *P. maritima* L.— (however, it lacks the key feature of subg. *Coronopus*, i.e., pilose corolla tube). On our trees, it groups with *P. arachnoidea* from Central Asia. Chinese *Plantago perssonii* Pilg. (including *P. lorata* (J.Z.Liu) Shipunov described from Central Asia: Shipunov, 2000a) robustly groups with *P. arachnoidea*. Morphologically unusual *P. reniformis* Beck from Balkans frequently also groups here with low support.

Another stable group consists of species from sect. *Micropsyllium*: Palearctic *P. polysperma* Kar. & Kir., *P. tenuiflora* Waldst. & Kit., and Nearctic *P. elongata* Pursh, *P. heterophylla* Nutt. and *P. pusilla* Nutt. Here we noted that geographically isolated, perennial *P. tenuiflora* from Öland (first described as separate species *P. minor* Fr.) does not group with typical *P. tenuiflora* but instead groups with *P. polysperma*.

The less stable but relatively consistent group forms around polymorphic *P. asiatica* L. from mainland China and Japan, including *P. schneideri* Pilg., *P. centralis* Pilg., and *P. cavaleriei* H.Lév. We found typical *P. asiatica* on Hawaii Island, thus extending the range of this East Asian species to mid-Pacific. However, all “*P. asiatica*” from mainland USA are either *P. major* or *P. rugelii* Decne. (Shipunov, 2017, 2019a).

*Plantago hakusanensis* Koidz. is a Japanese alpine endemic species with a distinct morphology. On our trees, it is branched closely to *P. asiatica*. In PE herbarium, we discovered the Yunnan sample labeled as “*Plantago zhongdainensia*” (*nomen nudum*), which morphologically might be considered similar to both *P. hakusanensis* and *P. asiatica*. Unfortunately, DNA data is not available from this sample.

Proximal to *P. asiatica* is also morphologically distinct *P. hasskarlii* Decne. from Java mountains. Another species from Southeast Asia, *P. incisa* Hassk., groups outside of *P. asiatica* clade(s). The unusual form collected from China is morphologically somewhat similar to the *P. densiflora* J.Z.Liu (synonymized with *P. asiatica* in the “Flora of China,” Li et al., 2011). However, this form, “*Plantago* sp. Hupeh1” has typically 4–6 large black seeds (and also large fruits), which is not in agreement with *P. densiflora* protologue. On our trees, it groups with *P. depressa* Willd. and allies (e.g., *P. komarovii* Pavlov and *P. camtschatica* Link). Besides, on “broad” trees, *P. depressa* robustly groups together with *P. macrocarpa* Cham. & Schltdl.; this grouping is also present on “tall” trees with less support.

American *P. eriopoda* Torr., *P. rugelii*, *P. sparsiflora* Michx., and *P. tweedyi* A.Gray robustly supported as a clade on “broad” trees. Here belong also two samples collected in Chihuahua desert (BRIT) from northern Mexico; these plants have many morphological differences from other species in this group but cluster together with *P. eriopoda* and *P. tweedyi*. *Plantago rugelii*, which morphologically is hard to tell from *P. major*, doest not group with this last species on any tree; these two species found to belong to different sections (Hassemer et al., 2019).

*Plantago major* does not branch closely to *P. asiatica*, which was pointed out in Hassemer et al. (2019). Instead, *P. major* s.l. groups with *P. japonica* Fr. & Sav., *P. cornuti* Gouan, *P. gentianoides* Sibth. & Sm. and *P. griffithii* Decne., albeit with low support. Sequences from polyspermous form of *P. major* (Morgan-Richards and Wolff, 1999) described as *P. uliginosa* F.W.Schmidt, are identical to the typical *P. major*. *Plantago griffithii*, which is frequently considered a form of *P. gentianoides* groups with the last species (in a strict sense) on our trees, but this grouping is unstable.

*Plantago pachyphylla* A.Gray and *P. hawaiensis* (A.Gray) Pilg. (both from Hawaii) group together, and also with *P. aundensis* P.Royen from New Guinea. Alpine form of *P. pachyhylla* from Kauai (labeled in HUH as “*Plantago nubicola* Tessene,” nomen nudum) clusters outside of *P. hawaiensis* + *P. pachyphylla* from Hawaii island.

Two New Zealand species, *P. triandra* Berggr. and *P. unibracteata* Rahn, always cluster together outside of the rest of subg. *Plantago*.

Our phylogenic trees, especially from the “tall” dataset, also provide the primary ground for the placement of little-studied or previously molecularly not studied forms, for example, for *P. laxiflora* Decne. This South African species is morphologically unusual for the region and groups outside of African species. Other African and Madagascan species, i.e., *P. africana* Verdc., *P. longissima* Decne., *P. palmata* Hook.f., *P. remota* Lam. and *P. tanalensis* Baker, tend to group with small support.

Most of *Plantago* sect. *Virginica* species do not group with high support. However, we note that Andean *P. oreades* Decne. always branches outside of the *P. australis* Lam. group. Another Peruvian form from this section was listed by Knud Rahn (MO herbarium note) as possible new species; on our trees, it groups with different members of the section, including South American *P. tomentosa* Lam. The second “unknown” from Peru, sect. *Virginica* sample from NY with a long stem (unusual in subg. *Plantago*) frequently groups with *P. tenuipala* (Rahn) Rahn from Columbia.

*Plantago firma* Kunze ex Walp. was typically considered as strictly Chilean species, but we have found its samples collected in Peru (USM). All Chilean and Peruvian *P. firma* samples robustly group together, and then with another species, bipolarly distributed *P. truncata* Cham. & Schltdl.

While there is a little confidence among branches which belong to the rest of sect. *Virginica*, we were able to place in that group molecularly those species which have not been sampled before, namely *P. argentina* Pilg., *P. berroi* Pilg., *P. buchtienii* Pilg., *P. dielsiana* Pilg., *P. floccosa* Decne., *P. jujuyensis* Rahn, *P. orbignyana*, *P. penantha* Griseb., *P. tenuipala* (Rahn) Rahn, *P. ventanensis* Pilg. and *P. venturii* Pilg.

Knud Rahn’s series *Oliganthos* species (*P. barbata* G.Forst., *P. correae* Rahn, *P. pulvinata* Speg., *P. sempervivoides* Dusén, and *P. uniglumis* Wallr. ex Walp.) were sequenced the first time as a totality. On our “tall” trees, the group does not have high support but clusters together with *P. moorei* Rahn, *P. tehuelcha* Speg. and *P. fernandezia* Bertero ex Barnéoud, all from South America’s Cone and surrounding islands.

Most of the Australian species form a low supported but relatively stable grade; here we were able to place some under-researched species: *P. antarctica* Decne., *P. depauperata* Merr. & L.M.Perry, *P. drummondii* Decne., *P. gunnii* Hook.f., *P. polita* Craven (New Guinea) and *P. turrifera* B.G.Briggs & al.

### PLANTAGO SUBG. CORONOPUS

On this stable trees (Fig. 4, 5B), the topology always supports the subdivision of sects. *Maritima* and *Coronopus*. Within sect. *Maritima*, we were able to place with confidence the rare Central Asian *P. eocoronopus* Pilg. (as a sister to the whole group) and North African *P. rhizoxylon* Emb. We detected the presence of the “true” *P. maritima* in South Africa (PRE herbarium); these samples are molecularly not different from the rest of *P. maritima*.

**Figure 4.**
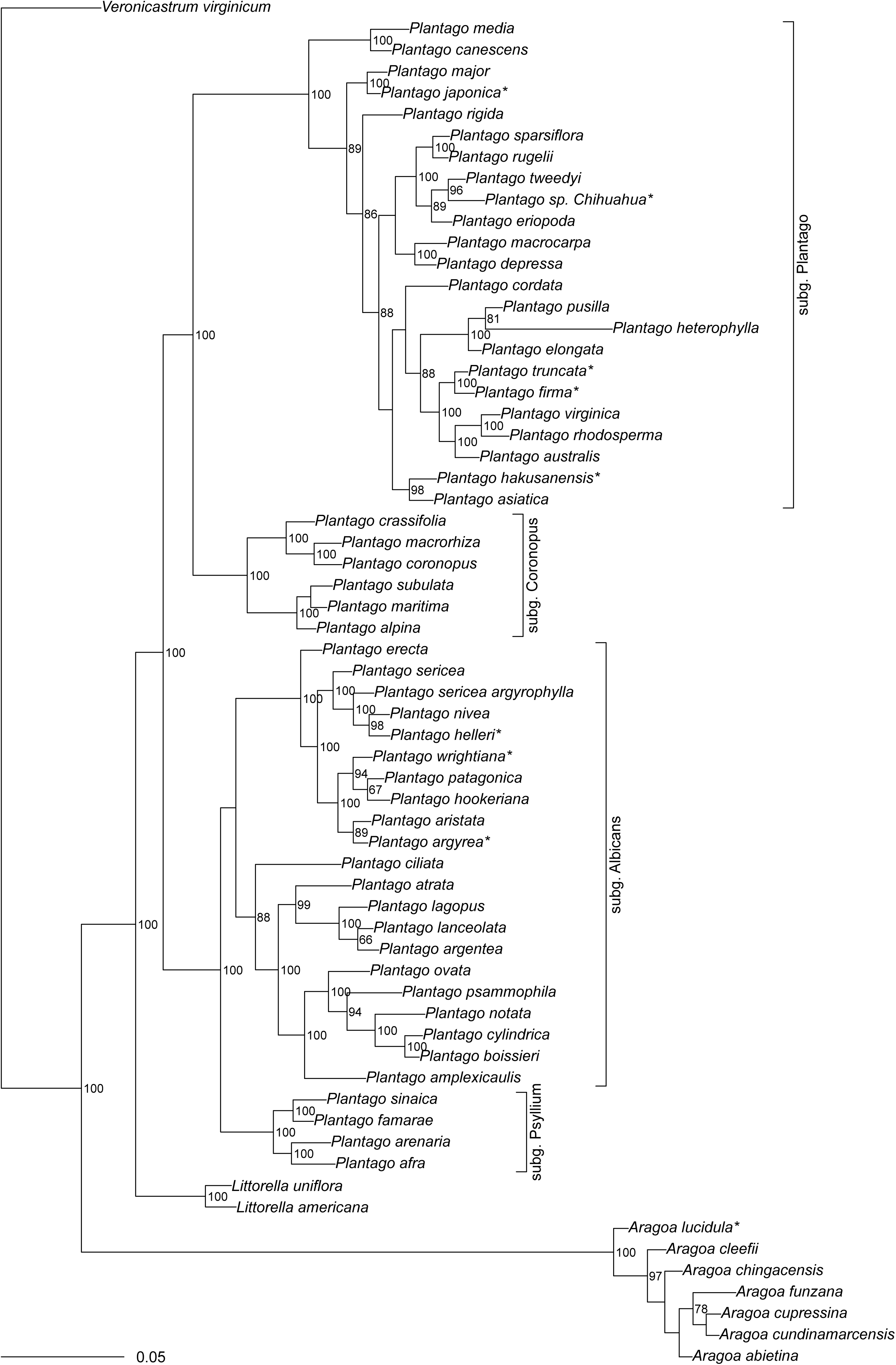
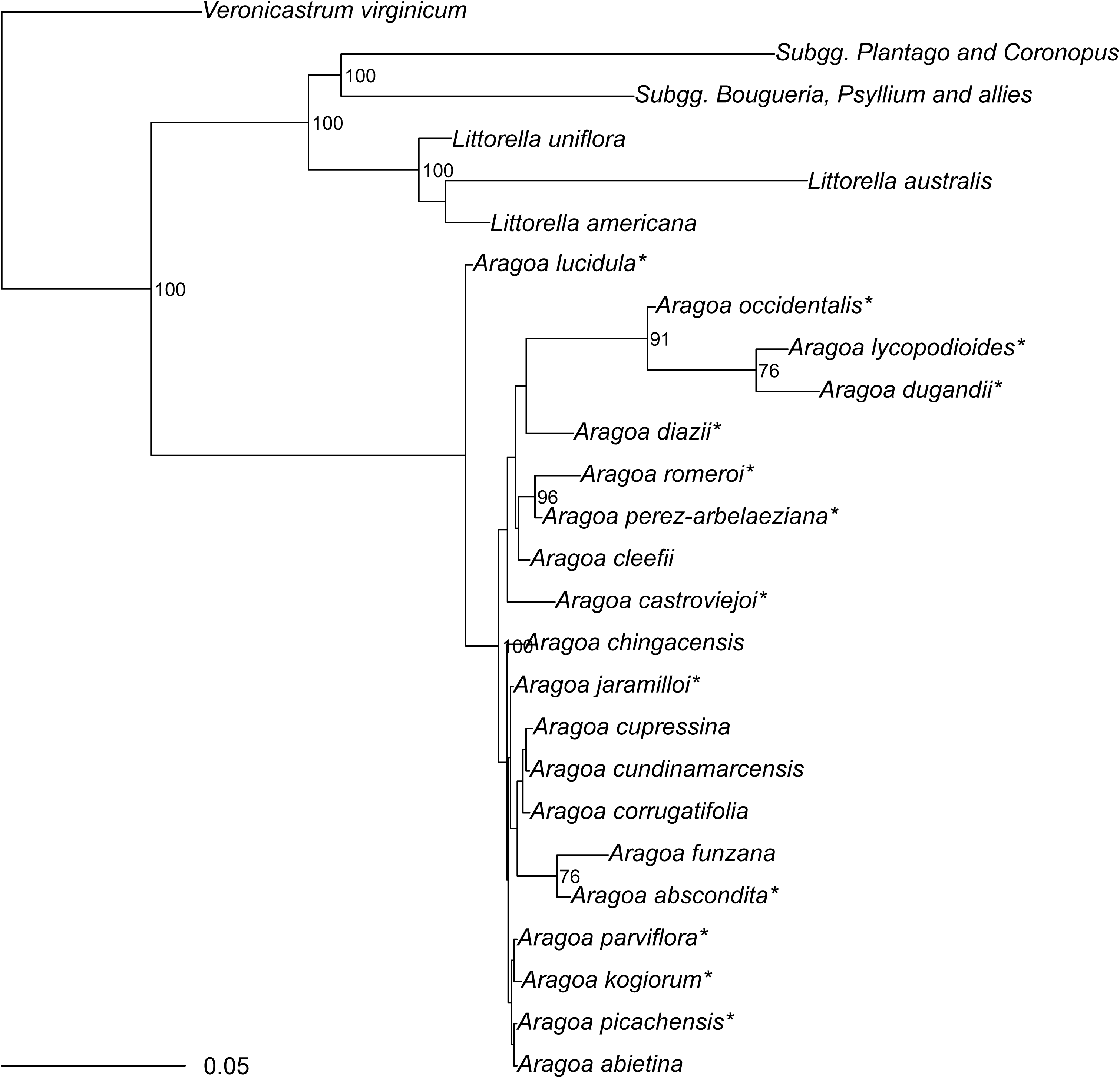
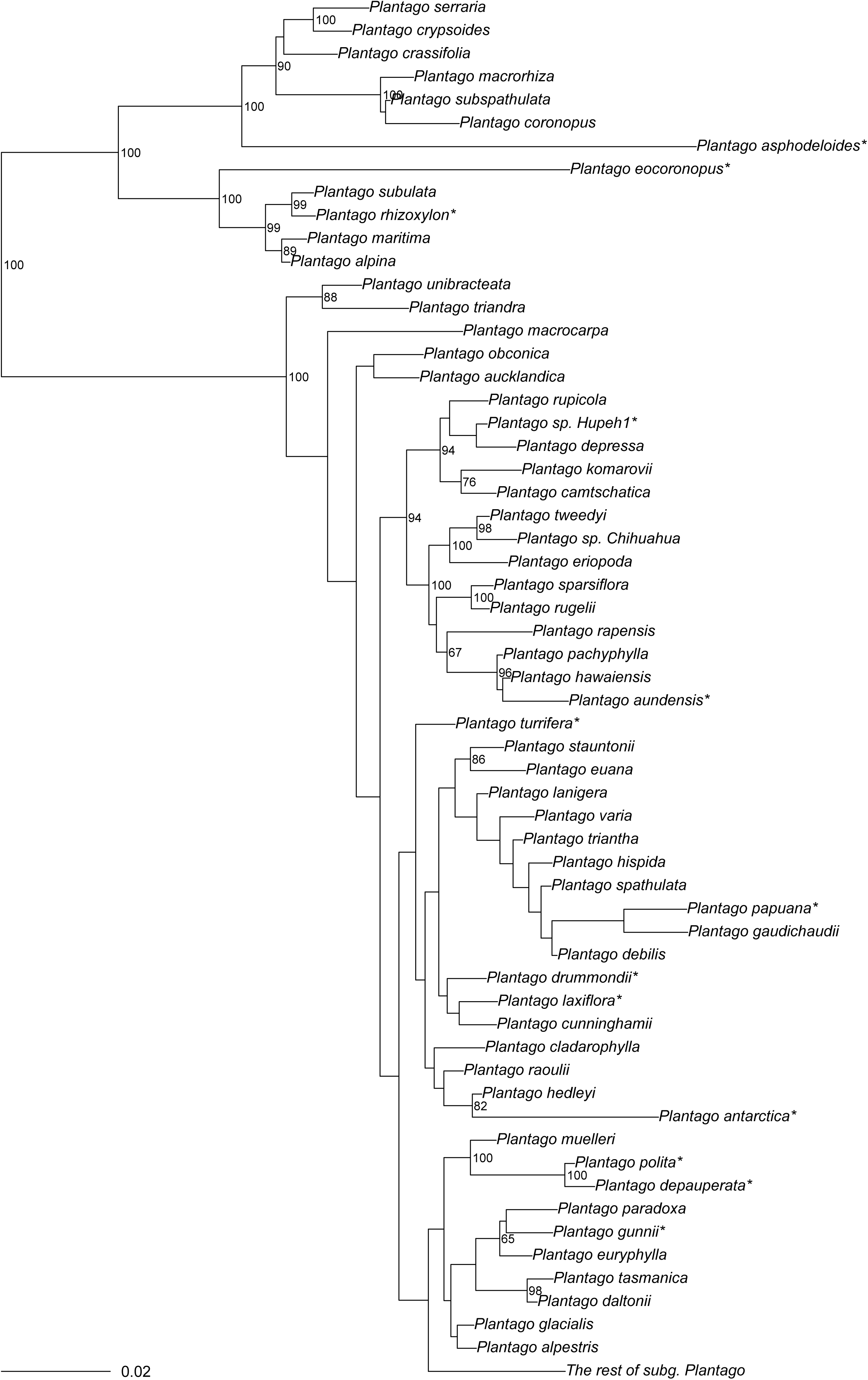
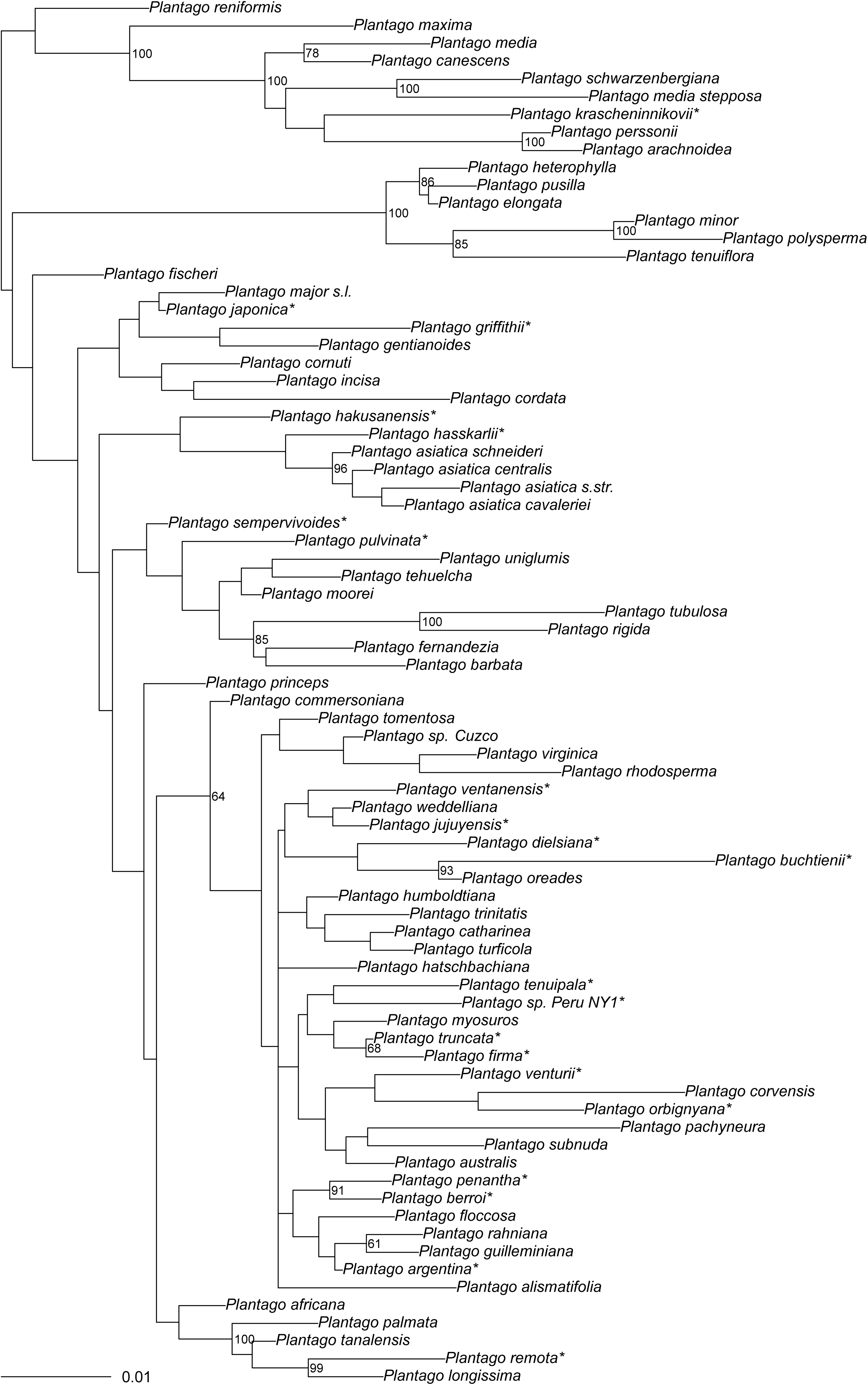
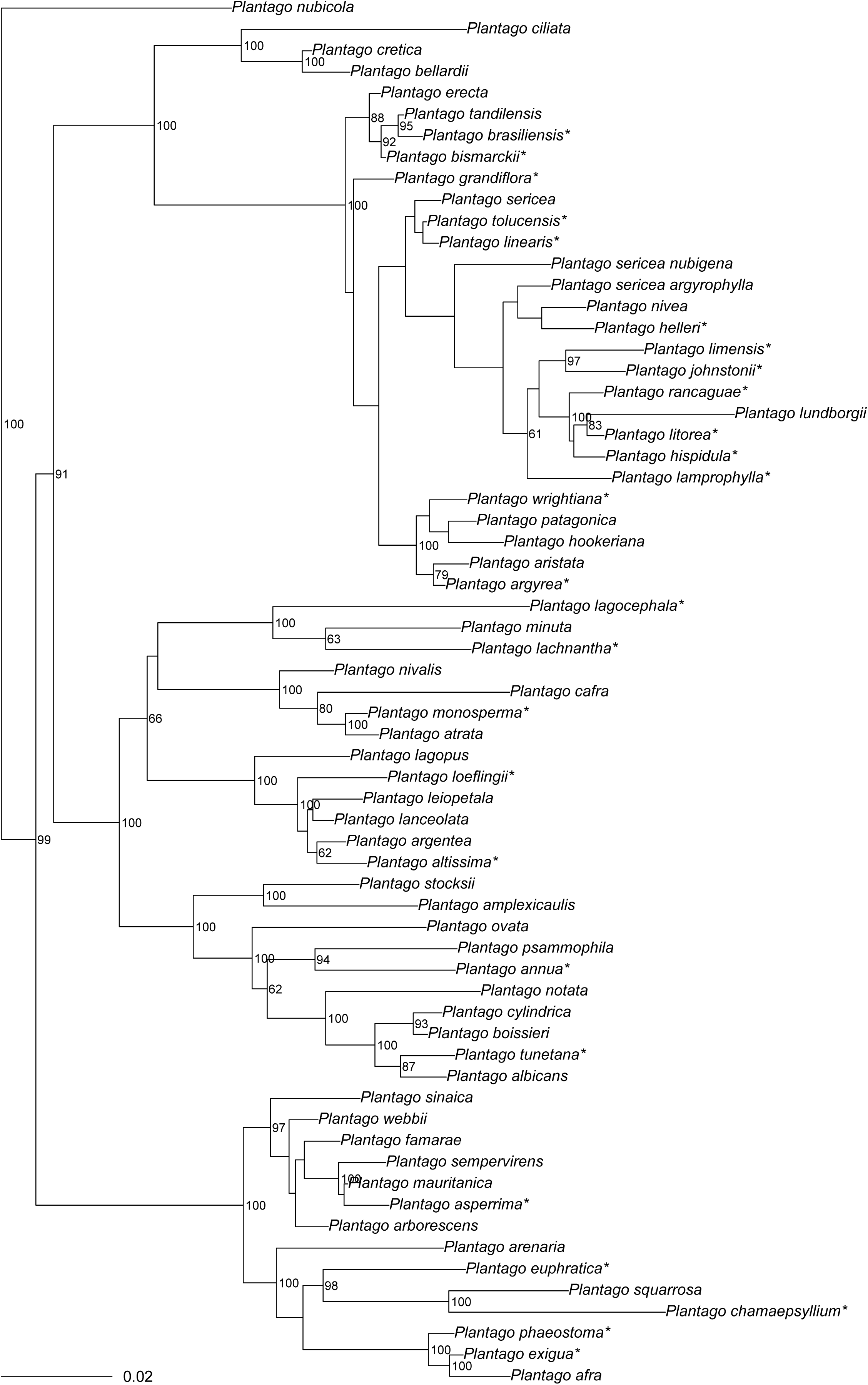
The phylogeny of *Plantagineae* obtained from the “broad” dataset (based on MB trees). Stars (*) mark species which have not been barcoded before. Numbers on nodes are Bayesian posterior probabilities (%).

Macaronesian *P. asphodeloides* Svent. is the sister to other species from sect. *Coronopus*, and North African *P. crypsoides* Boiss. is sister to Mediterranean *P. serraria* L.

### PLANTAGO SUBG. PSYLLIUM AND ALLIES

Within this stable group (Fig. 4, 5D), *P. nubicola* (Decne.) Rahn (which sometimes regarded as a separate genus *Bougueria*) is always branching basally. The following topology is prevalent: (*Psyllium* s.str., (“American clade,” “*Plantago ciliata* clade,” “Mediterranean clade”)); these we will describe in detail below.

*Psyllium* s.str. forms a robust, relatively long branch that split between mostly annual species with non-linear bracts (e.g., *P. squarrosa* Murray) and mostly perennial, woody species with narrow bracts (e.g., *P. arborescens* Poir.).

“*Plantago ciliata* clade” on “tall” trees is sister to “American clade”, whereas on “broad” trees *P. ciliata* Desf. is sister to “Mediterranean clade” (with lower support). This clade includes *P. ciliata* and two successfully sampled species from the sect. *Hymenopsyllium*, i.e., *P. cretica* L. and *P. bellardii* All.

“American clade” is, in essence, Rahn’s sect. *Gnaphaloides*. *Plantago erecta* E.Morris is variably at the base of this group, and *P. helleri* Small branches close to the southern *P. nivea* Kunth. The rest of North American species form a stable clade (which therefore roughly corresponds with Rahn’s ser. *Gnaphaloides*), where *P. aristata* Michx. and *P. argyrea* E.Morris form a subgroup.

On “tall” trees (where sampling is reliable), species from Central and South America form the *P. tandilensis* (Pilg.) Rahn + *P. brasiliensis* Sims + *P. bismarckii* Nederl. clade, *P. grandiflora* Meyen clade, *P. sericea* Ruiz & Pav. grade (incl. *P. lamprophylla* Pilg., *P. nivea*, *P. helleri*, *P. linearis* Kunth, and *P. tolucensis* Pilg.) and ser. *Hispiduleae* clade. The latter also includes *P. johnstonii* Pilg. and samples of *P. litorea* Phil. collected in Peru (thus extending the range of this Chilean species). Samples of some *P. sericea* subspecies do not branch together with the bulk of *P. sericea* samples (Fig. 5D).

**Figure 5.**
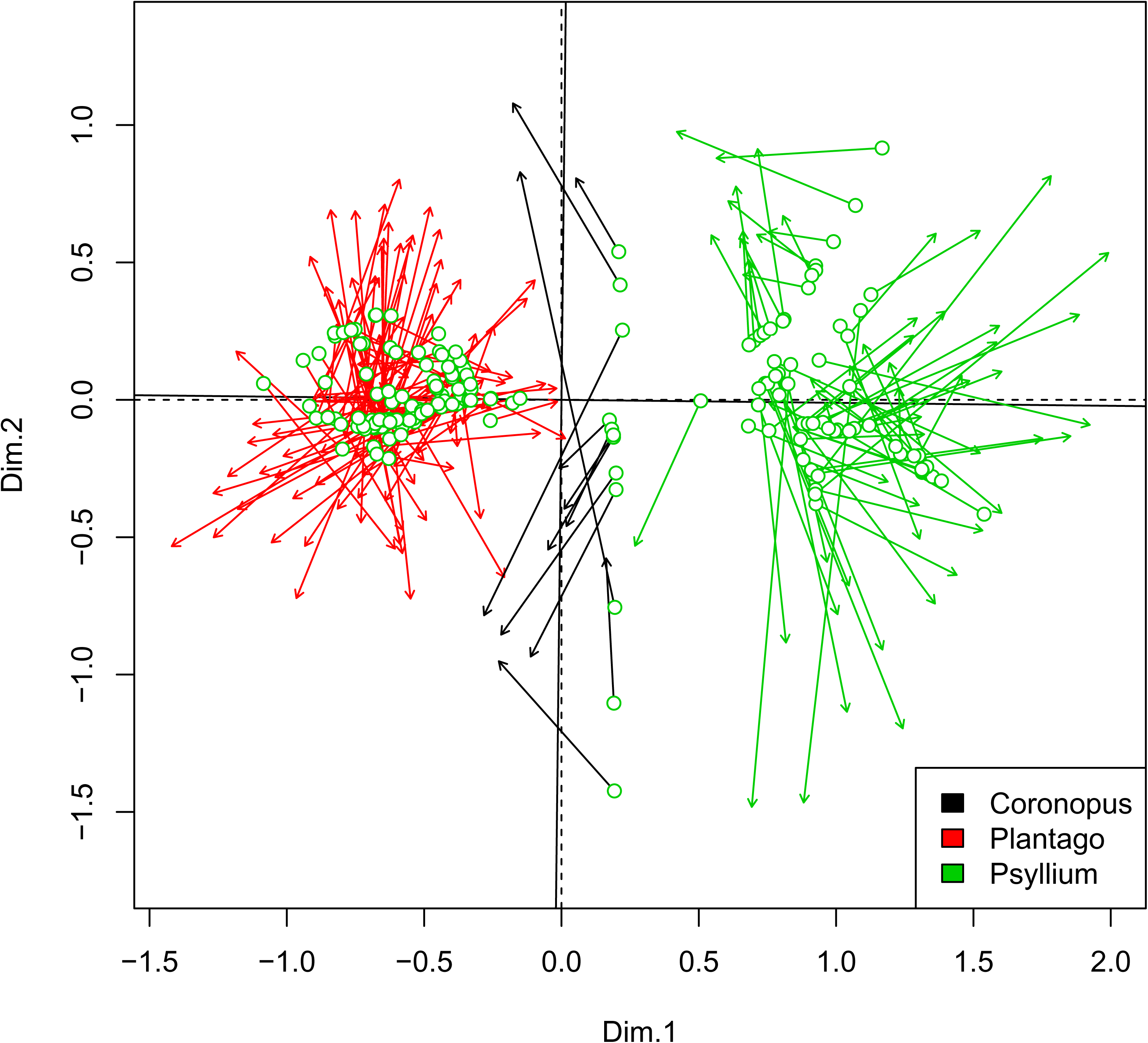
The phylogeny of *Plantagineae* obtained from the “tall” dataset (based on MB trees). (A) *Aragoa* and *Littorella* (B) *Plantago* subgenera *Coronopus* and *Plantago* (first part); (C) subgenus *Plantago* (second part); (D) subgenus *Psyllium* and allies. Stars (*) mark species which have not been barcoded before. Numbers on nodes are Bayesian posterior probabilities (%).

“Mediterranean clade” corresponds with sects. *Montana*, *Lancifolia*, and *Albicans* (except *P. ciliata*). The first subclade formed with members of the first two sections plus *P. lagocephala* Bunge and two species from the sect. *Albicans* ser. *Minutae*: *P. minuta* Pall. and *P. lachnantha* Bunge. Sections *Montana* and *Lancifolia* represented as proposed by Rahn (1996) except for *P. loeflingii* L. (it groups with sect. *Lancifolia* instead of sect. *Montana*).

“Mediterranean clade” 2nd subclade consists mostly of species from the sect. *Albicans*. *Plantago amplexicaulis* Cav. (sect. *Bauphula*) and *P. stocksii* Boiss. ex Decne. (sect. *Albicans* ser. *Ciliatae*) group together on the base of this group. The next branch(es) is *P. ovata* Forssk. and *P. psammophila* Agnew & Chal.-Kabi + Ethiopian *P. annua*. The rest of this subclade consists of species from ser. *Albicantes* and *Ciliatae*, plus *P. notata* Lag.

### MORPHOLOGICAL AND COMBINED ANALYSES

The Procrustes analysis allows for the embedding of two multivariate datasets (Peres-Neto et Jackson, 2001; Balbuena et al., 2013) and related statistical tests. Our molecular and morphological datasets are significantly correlated (correlation = 0.7748, significance = 0.001 based on 999 permutations), but individual placements are variably shifted (Fig. 6).

**Figure 6.**
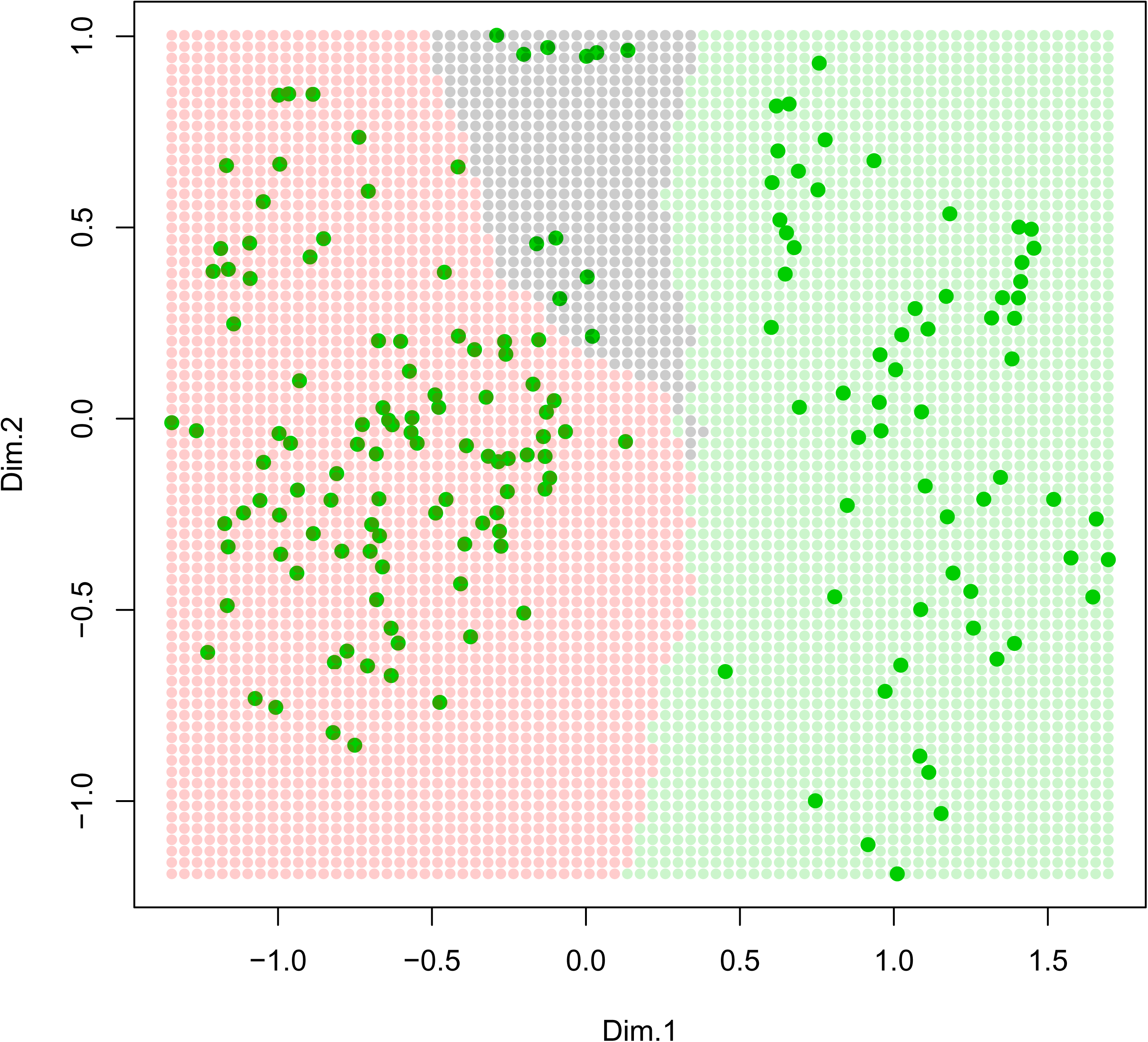
Scatterplot of the data points from joint molecular and morphological datasets (genus *Plantago* only) after the Procrustes superimposition. Differences in location of each species designated with arrows. These arrows start from the location defined by molecular data and target to the location defined by morphological data. The angle between solid and dashed axes reflects the overall Procrustean distance.

Even after intensive sampling, some species of the group still lack the molecular information. There are also species where only ITS2 sequences are available. With *k-*nearest neighbor machine learning (Fig. 7), we obtained the section/series placements of these *Plantago* species. More than half of them placed with high (> 90%) bootstrap confidence (Table 1). In the case of *Aragoa*, we only operated with an existent classification (Fernández, 1995) combined with phylogenic trees, and our placements here structured as trios of the most closely related species (Table 1).

**Figure 7.**
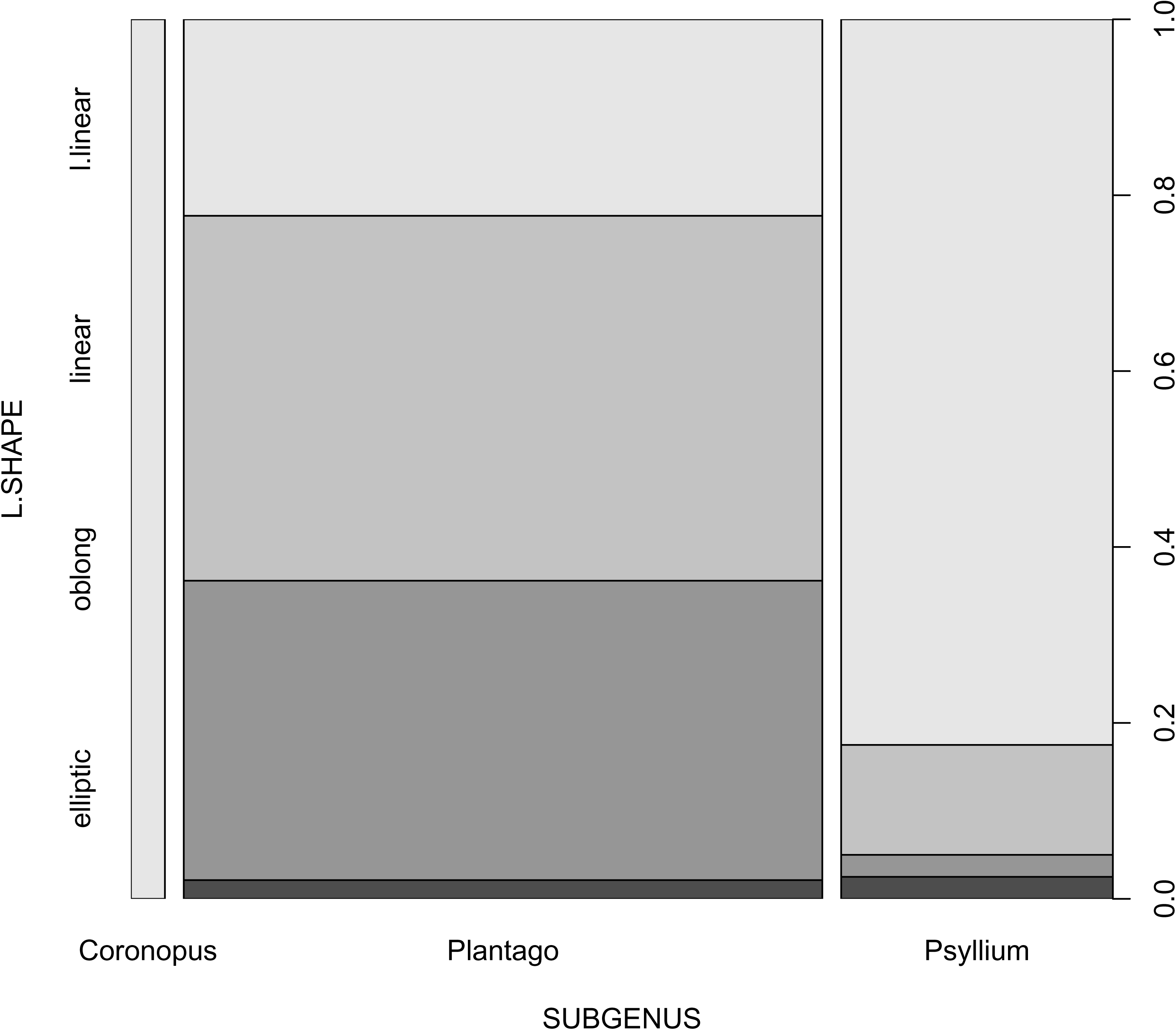
*k*-nearest neighbors (*k*-NN) plane predicts *Plantago* subgenera from the combined (molecular and morphological) data. The combined data was projected into a 2-dimensional plane. Solid dots correspond to subgenera *Plantago* (red), *Coronopus* (black), and *Psyllium* with allies (green). For each location shown as a semi-transparent dot, subgeneric placement learned with the *k*-NN algorithm, and the dot was colored following this prediction. Now, if any new species appear which corresponds with any of these semi-transparent dots, its subgeneric placement is already predicted.

Chi-squared tests returned the appropriate p-values when comparing leaf shapes (morphometric dataset) with subgenera, sections, and macro-regions (0.0005, 0.016, and 0.0055 respectively) and typically large effect sizes (corrected Cramer’s V 0.39, 0.34 and 0.26 respectively, see also Fig. 8). At the same time, the relative sizes of the stalk and spike were not significant. There is also a support for the pattern of gapped spike *vs*. sections (p-value 0.0004 and 0.48 corrected Cramer’s V).

**Figure 8.**
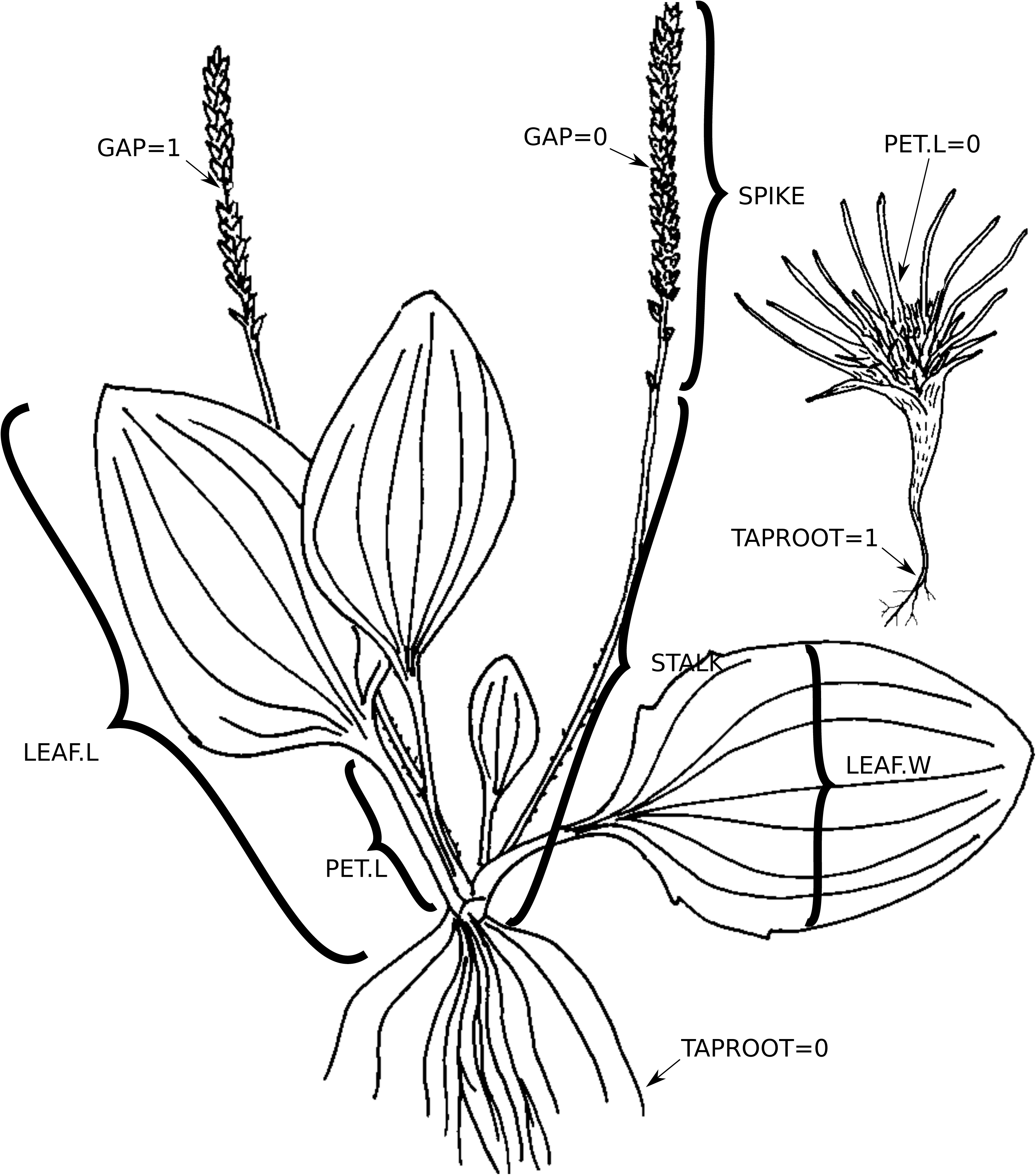
*Plantago* leaf shapes *vs*. subgenus. This plot derived from the cross-tabulation, which uses a morphometric dataset for leaf shapes.

We used the average or maximum Spearman correlation between morphological matrices and phylogenetic trees based either on a “tall” dataset or molecular-morphological dataset to determine the “molecular weight” of morphological characters. Most “heavy” among morphometric characters Fig. 9A) was the presence of taproot, and then the length of leaves (Fig. 9B). The top 10 binary morphological characters (Fig. 9C) were: seed surface type 4 (with elongated ridges: Shipunov, 1998b), long corolla (> 4 mm or > 3 mm) lobes, opposite leaves, presence of pedicel, truncated base of leaf blade, presence of glandular hairs, elongated stem, antrorse hairs on the stalk, and presence of non-glandular hairs with the strongly refracted walls.

**Figure 9.**
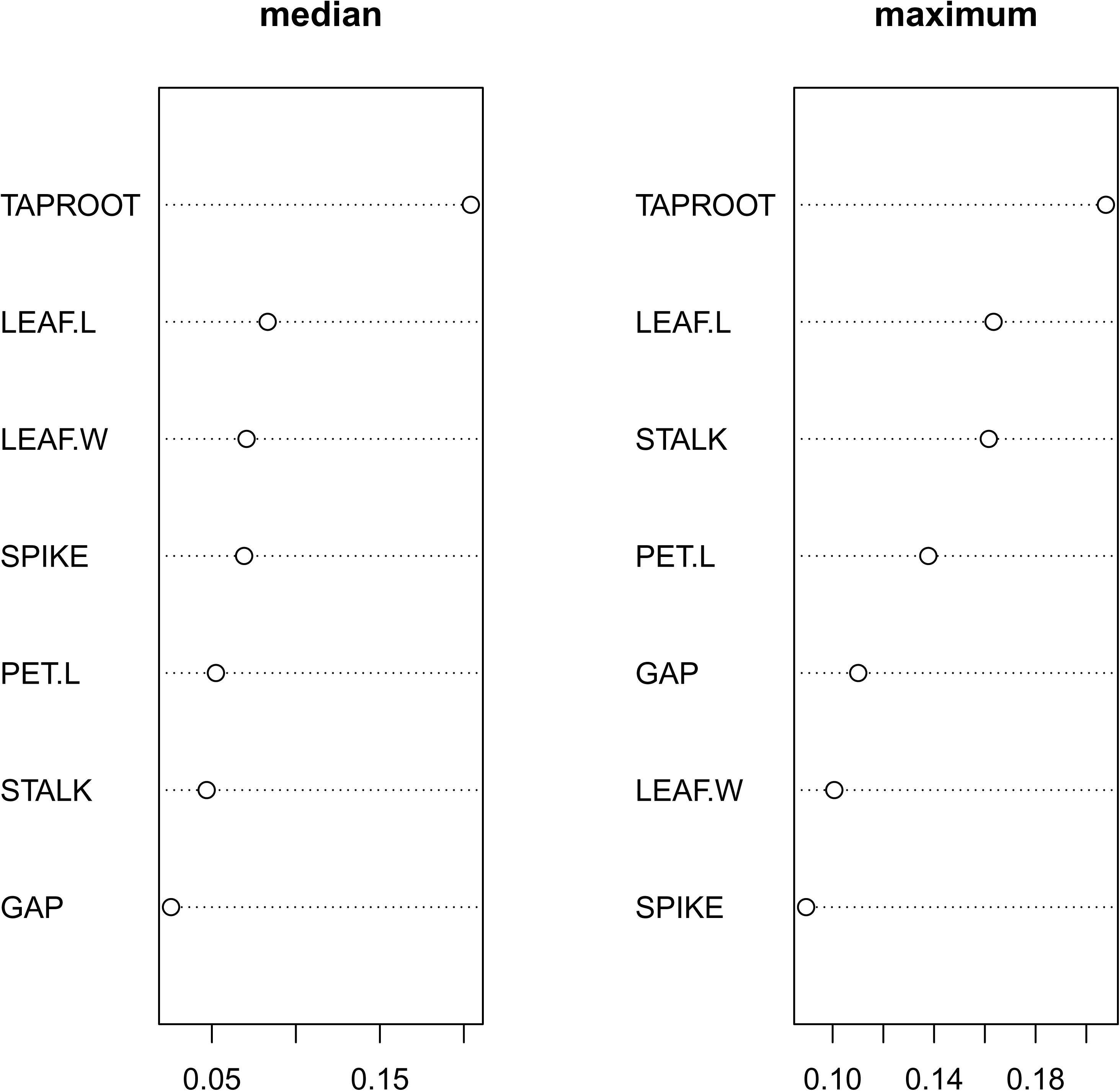
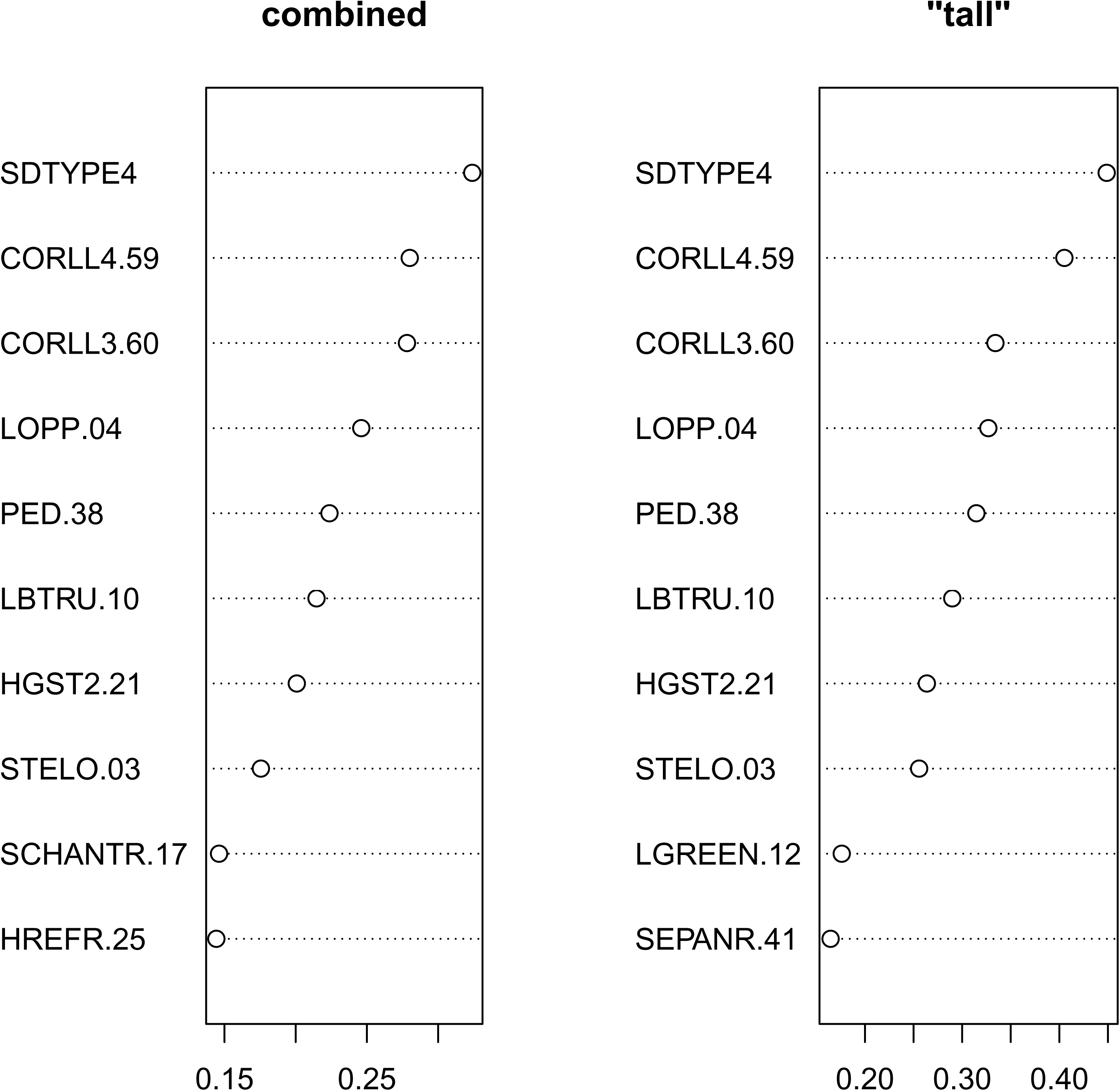
Morphological characters in *Plantago*: (A) morphometric measurements of seven “spot” characters; (B) “molecular weights” (median and maximum average Spearman correlations on 1000 bootstrap replicates) of morphometric characters; (C) “molecular weights” (average correlations with “tall” tree and combined molecular-morphology tree) of binary characters, character abbreviations explained in the text and Support Table 4.

We used recursive partitioning (Venables et Ripley, 2002) to construct the classification trees of the group (Fig. 10A-B). With binary morphological binary characters, we employed three runs, excluding characters used in the previous run. The resulting recursive classification trees employed 20, 20, and 19 characters (out of 115) and had 25.3%, 35.5%, and 48.1% misclassification errors, respectively. With morphometric characters, the resulted tree used all seven characters and returned a 75.5% misclassification error.

**Figure 10.**
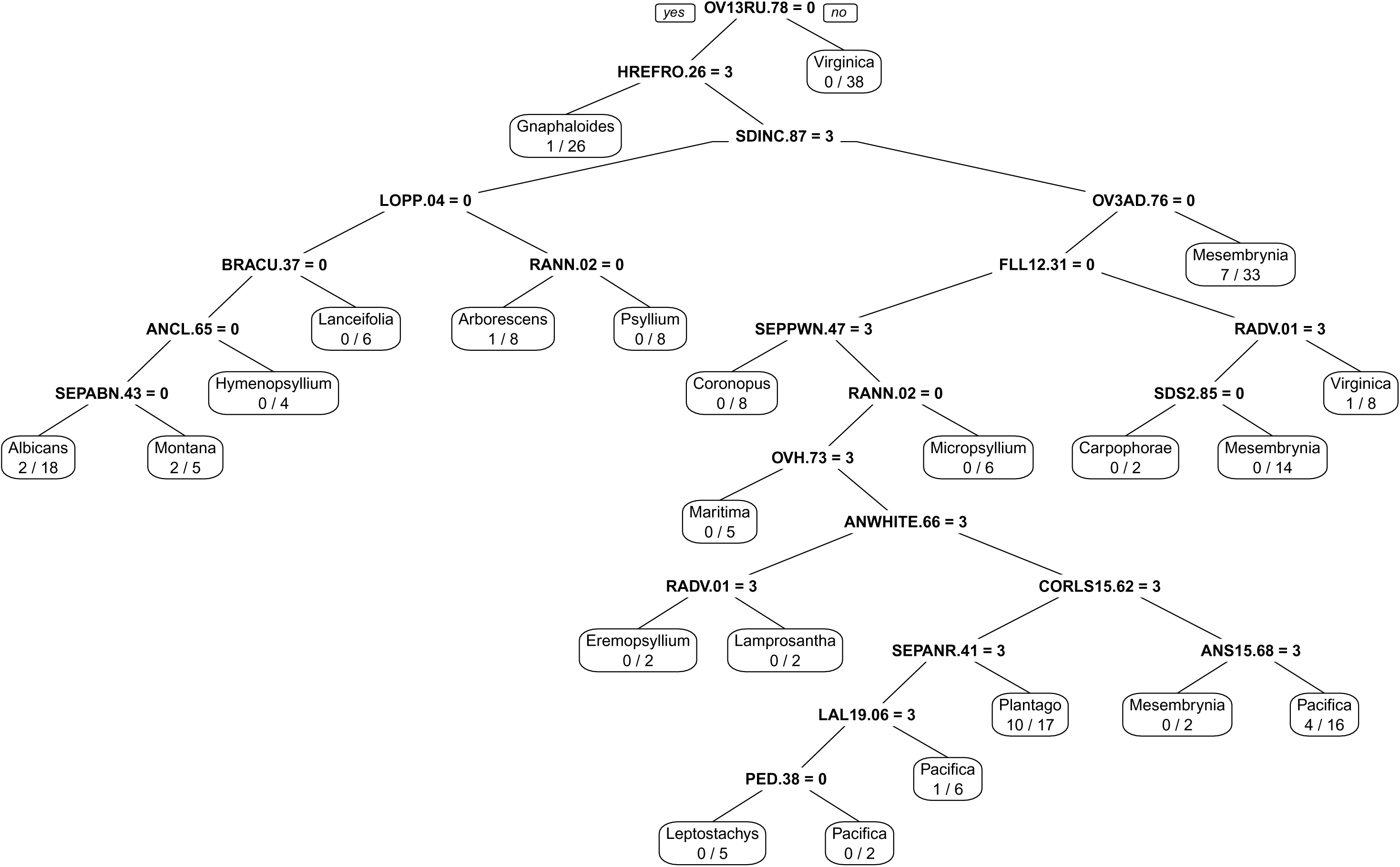
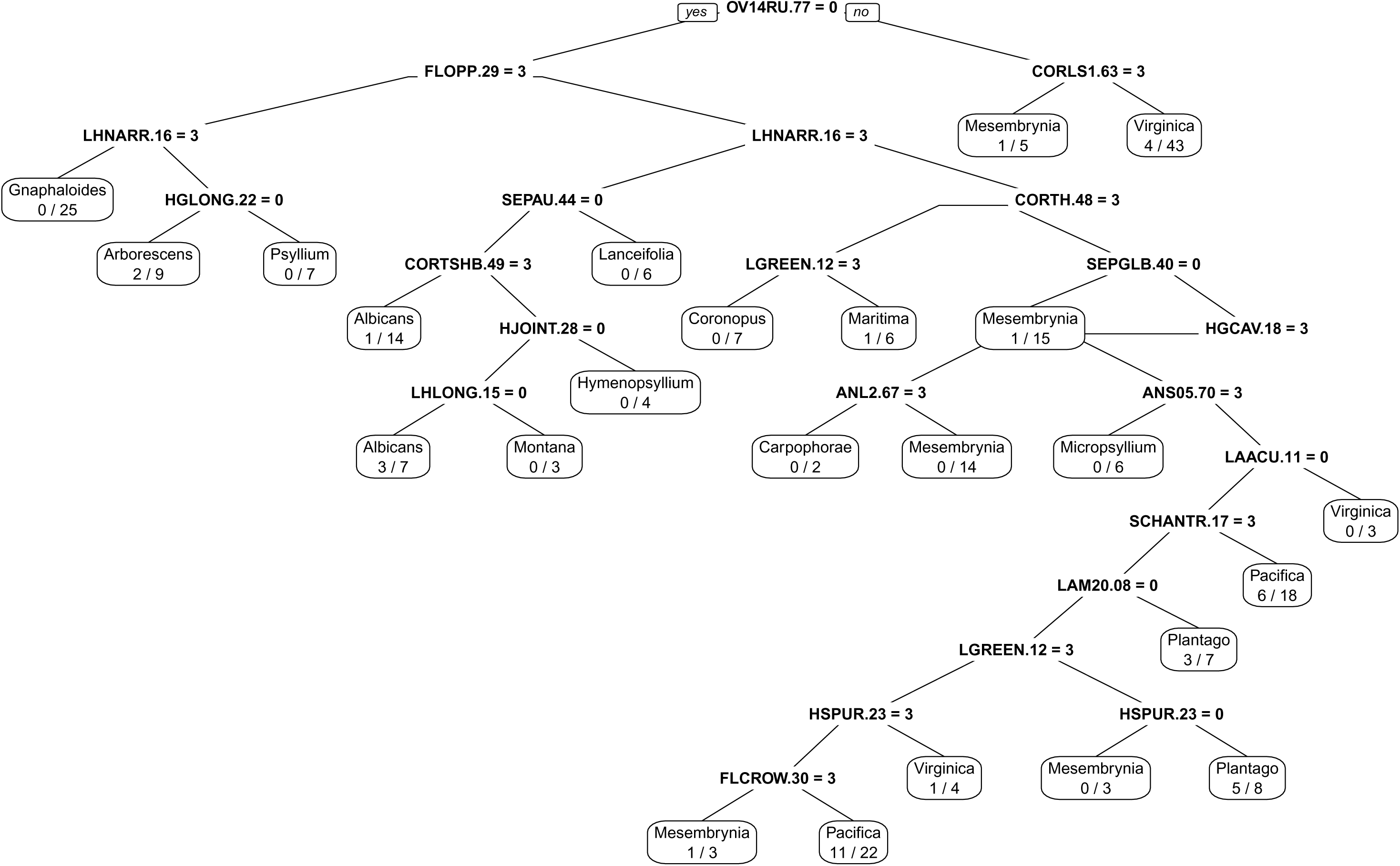
Recursive partitioning of *Plantago* sections with morphological characters, the prototype of diagnostic key: (A) first run and (B) second run on binary morphological characters. Character

## DISCUSSION

### PLANTAGINEAE

There is unequivocal support for the *Aragoa* (*Littorella* (*Plantago*)) structure of the group phylogeny (Fig. 1, 2, 4). This structure is concordant with the current understanding of the evolution of the tribe, including the evolution of flower symmetry (Preston et al., 2011). Reduced leaf morphology might also be explained with this phylogeny.

Bello et al. (2012) included only three species (plus two hybrids) of *Aragoa* and did not resolve their relationships. Here for the first time, the significant part of the *Aragoa* diversity was reviewed using the data obtained in molecular research (Fig. 4, 5A). In general, there is some support for grouping obtained in the research solely based on morphology (Fernández, 1995).

However, some morphologically unusual species like *A. dugandii* with relatively broad leaves, and *A. perez-arbelaeziana* with long tubular flowers do not make separate clades but are clustered together with more “typical” species (*A. lycopodioides* and *A. romeroi*, respectively, both from the main clade). The relatively broad, patent leaves and the simple inflorescence of *A. dugandii*, together with relatively large flowers, might be therefore interpreted as an adaptation to environments where moisture loss is not so critical as for many other species of the genus. As for *A. perez-arbelaeziana*, the presence of long, pendulous, yellowish corollas, unusual in the genus, can be interpreted as a result of recent adaptive radiation to a specific type of pollinator (hummingbirds: Fernández, pers.obs.) which does not entail the significant modifications in vegetative structures.

The machine learning placement of three unsampled *Aragoa* species resulted in the selection of possible “candidate neighbors” (Table 1) from the same main *Aragoa* clade.

*Littorella*, with three distinct species lineages, is robustly supported as a sister group of *Plantago.* Our data (Fig. 4, 5A) agree with a view of morphologically and ecologically outstanding *Littorella* as a separate generic lineage (Hoggard et al., 2003).

*Plantago* taxonomy is the most complicated part of our research. In general, there are no significant conflicts with the most recent studies of the genus based on morphology summarized in Rahn (1996). However, there are numerous findings and differences to emphasize.

All our analyses reproduce ((*Coronopus*, *Plantago*), *Bougueria*, (*Psyllium*, *Albicans*)) backbone (Fig. 4, 5B-D). Based on our trees, it is possible to keep the latter three subgenera as such based on strong molecular and morphological support. Alternatively, it is also possible to lump two last groups in one subg. *Psyllium* (Rønsted et al., 2002). Apart from the molecular evidence, this *Psyllium* s.l. union has reliable morphological support: two (rarely one) ovules and seeds, cotyledons perpendicular to the placenta, placenta side of seed deeply concave, hairs with a basal cell shorter than broad, leaves are often linear and spike usually relatively short (Rahn, 1996). The third alternative would be to accept this union as a separate genus, *Psyllium* Mill. s.l. (Shipunov, 1998a, 2000a), but this will significantly decrease the nomenclatural stability in the group.

### PLANTAGO SUBG. PLANTAGO

In general, our trees in this part (Fig. 5B, 5C) do not provide a clear, well-resolved picture as in plastome studies (Hassemer et al., 2019). However, they contain new information about possible placements of previously not molecularly studied species, and also about species which were not included in plastome research. In several cases (Hassemer et al., 2019), the position of species in the specific section was “inferred based on the accumulated knowledge.” Many of such species are now more formally distributed between these sections (Table 1), and this is reflected in our working classification of *Plantagineae* (Support Table 1).

For example, it is mentioned in Hassemer et al. (2019) that “…based on our phylogeny it is impossible to infer the position of the five unsampled American species in Rahn’s (1996) series *Oliganthos* (*P. barbata*, *P. correae*, *P. pulvinata*, *P. sempervivoides*, and *P. uniglumis*)”. Our current trees (Fig. 5C) place these species (except *P. correae* for which we have no data) in one clade, which also includes *P. rigida*, *P. tubulosa*, *P. moorei*, *P. tehuelcha*, and *P. fernandezia*, the placement which is well justified geographically. Morphologically unusual *P. sempervivoides* is sister to the remainder of the group.

Two sect. *Carpophorae* species are sister to *P. fernandezia* + *P. barbata*. This group likely has an Andean origin, and *P. fernandezia* might, therefore, arrive at Juan Fernández from South America (Stuessy et al., 2017; Iwanycki Ahlstrand et al., 2019). Position of *P. fernandezia*, *P. tehuelcha*, and sect. *Carpophorae* species are different on the plastome trees (Hassemer et al., 2019), but these trees have low support exactly in these parts.

Another complicated group was not resolved entirely, but we provide several insights for the placements and phylogeny in general of species around polymorphic and widespread *P. asiatica* (Matsuo, 1989, Ishikawa et al., 2006; Ishikawa et al., 2009). Most of these forms cluster together (Fig. 4, 5C) and separately from the *P. major* clade; thus, morphological similarity here does not justify taxonomic closeness. Japanese endemic *Plantago hakusanensis* is either within this group or branches basally; the same is true for *P. hasskarlii*. Relations of *P. hakusanensis* and *P. hasskarlii* mandate more in-depth research.

Molecular data from *Plantago japonica*, together with ecology and morphology, suggests that this species is likely distinct from *P. major* (Matsuo, 1989; Ishikawa et al., 2009).

Samples from the Hubei province of China (“*Plantago* sp. Hupeh1”) have an appearance of *Plantago asiatica* but branch near *P. komarovii* + *P. camtschatica* clade (Fig. 5B). As the allopolyploid origin of tetra-or hexaploid *P. asiatica* and *P. rugelii* was suggested by Ishikawa et al. (2009), we cannot exclude the possibility of the allopolyploid origin of these Hubei samples (with ITS kept from the sect. *Pacifica* parent). We believe that a thorough study of Chinese, Korean, and Japanese *Plantago* subg.

*Plantago* species is required before reaching any robust conclusions. In the light of Ishikawa et al. (2009) report, the recent historical origin of *P. rugelii* (which branches closely to *P. sparsiflora* on our trees but morphologically hardly distinguishable from *P. major*) might also be justified.

There are seven endemic *Plantago* species described from New Guinea (*P. aundensis*, *P. depauperata*, *P. montisdicksonii* P.Royen, *P. papuana* P.Royen, *P. polita*, *P. stenophylla* Merr. & L.M.Perry and *P. trichophora* Merr. & L.M.Perry), but DNA extraction from available samples failed in 90% of cases. Nevertheless, we were able to place four of these species (Fig. 5B): *P. aundensis* with Hawaiian *P. pachyphylla* s.l. and *P. hawaiensis* (the third Hawaiian species, *P. princeps* does not hold a stable position on our trees); *P. depauperata* and *P. polita* with Australian *P. muelleri* Pilg.; and *P. trichophora* with Australian *P. gaudichaudii* Barnéoud. Still, much more sampling is needed, and Hawaiian species also deserve a closer look (Hassemer et al., 2019).

Morphologically unusual samples (Fig. 11) from Chihuahua (Mexico) are physiognomically similar to *P. gaudichaudii* from Australia and *P. remota* from South Africa, but we believe that it is the clear example of morphological convergence. On our trees, these samples belong to sect. *Pacifica* clade and branch together with Midwestern *P. eriopoda* and *P. tweedyi* from the Rocky Mountains region (Fig. 5B).

The same *Pacifica* clade includes on “broad” trees *Plantago macrocarpa*, the Northern Pacific seashore species. On the “tall” trees (Fig. 5B), the placement of this species is not stable.

*Plantago krascheninnikovii* is superficially similar to the inland forms of *P. maritima* and therefore treated as a member of subg. *Coronopus* (Shipunov, 2000b). Known populations of this rare Urals species do not typically form the ripe seeds (Shipunov, 1998a), thus preventing the correct placement on the base of morphology. Here we the first time confirm its similarity with other *Lamprosantha* species, for example, Eastern European *P. schwartzenbergiana* (Fig. 5C).

The southern tetraploid forms similar to *Plantago media* but with thick, erect, grayish leaves (Shipunov, 1998a, 2000b), often regarded as *P. urvillei* Opiz or *P. media* subsp. *stepposa* (Kuprian.) Soó, are distant from *P. media* s.str. (Fig. 5C), thus necessitating the separation of this taxon. However, the diversity of *P. media* s.l. is still far from being fully understood (Palermo et al., 2010), and more research is needed to draw robust taxonomic conclusions. Here should be noted that *P*. *media* subsp. *stepposa* must not be mixed with the similarly looking shade, mesophytic plants of *P. media* (Shipunov, 1998a).

Small perennial sect. *Micropsyllium* plantains from Öland described as *P. minor* Fries are morphologically distinct and geographically isolated from other species of this section. However, common garden experiments (Tsinger, 1905) led to the conclusion that they are conspecific with *P. tenuiflora* while our Öland sample is proximal to *P. polysperma*. We believe that more research on Öland plantains is needed to understand the taxonomic status of these forms better.

Andean *Plantago oreades* Decne. with distinct morphology (narrow leaves, long inflorescences, broad bracts, 1–3 seeded fruit, thick roots) was nevertheless included into broadly understood *P. australis* (Rahn, 1974). On our trees (Fig. 5C), it almost always separate from the other *P. australis* samples. Therefore, we propose here to re-establish this species.

There are many local endemics in sect. *Virginica* (Hassemer, 2019a). Knud Rahn labeled morphologically unusual samples (MO) collected in the Cuzco area (Peru) as possible new species (Fig. 12). These samples are always separate on our trees (Fig. 5C).

### PLANTAGO SUBG. CORONOPUS

Our trees (Fig. 4, 5B) provide one of the most comprehensive phylogenies for the subg. *Coronopus*, and are in concordance with the recent work of Höpke et al. (2019). We were able to place those species which have not been the subject of molecular studies. The most interesting are positions of Canarian *P. asphodeloides* Svent as sister to the rest of species from sect. *Coronopus*, and *P. eocoronopus* Pilg. as sister to the rest of the sect. *Maritima*. The latter species is the rare Afghan plant, practically absent in collections. Pilger (1937) guessed this species to be ancestral for the section, and now we can support this view on the base of both molecules and morphology (Shipunov, 2000a). All our “*P. schrenkii*” C. Koch samples from the Arctic are identical to *P. maritima* (Shipunov, 2015).

### PLANTAGO SUBG. PSYLLIUM AND ALLIES

Generally, this part of our trees (Fig. 4, 5D) is the most congruent with the classification of Rahn (1996). Our data is also congruent with the most complete (until now) sampling of the group (Rønsted et al., 2002) and provides robust support for many sub-groupings, which is reflected in our working classification (Supplement Table 1).

Among other results, we found that likely extinct *P. johnstonii* (Hassemer et al., 2018a) branches close to the coastal annual *P. limensis* and therefore belongs not to ser. *Brasilienses* but to ser. *Hispiduleae*. It is possible then that the perennial life form of the former species is the result of adaptation to the “Lomas” microclimate.

The most recent review of the sect. *Lancifolia* (Hassemer, 2019b) is in agreement with our results but also provides new insights for the classification of this Mediterranean taxon. More research is needed to understand the relations of rare endemic species in this group.

### MORPHOLOGICAL AND COMBINED ANALYSES

With the Procrustes analysis (Fig. 6), we found that the overall “picture of diversity” is retained between morphological and molecular approaches. In other words, the correspondence between these analyses is high and allows us to run the combined analyses. These analyses, in turn, allow for the *k*- nearest neighbor (Fig. 7) placement of several species which might be otherwise *incertae sedis* in our working classification (Table 1, Supplement Table 1).

Morphometric characters that reflect general bio-morphological features of species should be assistive in identification. Our analysis provided several insights into this field. We found several repetitive morphometric patterns, “refrains” (Meyen, 1987) within sections and subgenera (Fig. 8); this is additional evidence that morphometric characters should work better within sections (Höpke et al., 2019). We also found several morphological and morphometric characters most correlated with molecular phylogenies (Fig. 9). The morphometric characters are especially interesting because the analysis was performed on the *same* samples and not on higher units like species descriptions. Most notable is the importance of the presence of taproot, which is another argument for collecting plantains always with carefully preserved underground parts. Among the binary morphological characters, attention should be paid on the research of seed surface characters (Shipunov, 1998b), highly correlated with molecular data (Fig. 9C).

Producing of identification keys is a complicated task in plantains. These keys must take into account the high variability and overlapping of the most distinctive characters used in *Plantago* taxonomy (Hassemer et al., 2019). Therefore, it might be desirable to employ here results of machine learning techniques such as recursive partitioning. Our partitioning trees (Fig. 10A-B) allow the distinguishing with minimal possible errors on the base of the few most informative characters. As the identification power of trees was relatively high, we decided to provide the dichotomous key for sections based on three runs of classification tree with binary characters and one run with morphometric characters. This prototypic key might serve as a framework for the future development of comprehensive keys for the whole group:

1. Ovary with 1–3 ovules and a rudiment of an upper compartment on the adaxial side of the placenta. Corolla lobes longer than 1 mm. Flowers are mostly cleistogamous; corolla lobes form a beak … sect. *Virginica*

– Ovary structured otherwise. Corolla lobes short or long. Flowers are mostly chasmogamous, corolla in most (but not all) species does not form a beak … 2.

2. Non-glandular hairs with joints are strongly refracting, walls between cells oblique. Hairs on leaves narrow, less than 0.04 mm … sect. *Gnaphaloides*

– Strongly refracting joints absent. Hairs on leaves (if present) variable … 3.

3. The inner side of the seed is deeply concave … 13.

– The inner side of the seed is not deeply concave … 4.

4. Ovary with a third compartment at the top on the adaxial side of the placenta, or with a rudiment of it, seen as a thickening at the apex on the posterior side of the ripe placenta. If this compartment absent, then there are few flowers in the inflorescence; no adventitious roots and seeds are longer than 2 mm. Sepals are glabrous on the back … sect. *Mesembrynia*

– Ovary without the third compartment, other character combinations are different … 5.

5. Less than four flowers per inflorescence. Carpophore present … sect. *Carpophorae*

– Inflorescence with more than 12 flowers. Carpophore absent … 6.

6. Posterior sepals with the membranaceous, very conspicuous wing on the back. Leaves are usually remaining green on drying. Corolla tube hairy. Annuals; leaves are often dentate or even dissected … sect. *Coronopus*

– Posterior sepals without a conspicuous wing on the back. Leaves dry differently. Corolla tube hairy or glabrous. Annuals or perennials; leaves with the whole margin or sometimes dentate … 7.

7. Annuals. Anthers usually less than 0.5 mm long … sect. *Micropsyllium*

– Perennials. Anthers are longer than 0.5 mm … 8.

8. Ovary hairy. Corolla tube hairy. Leaves usually do not remain green on drying … sect. *Maritima*

– Ovary glabrous. Corolla tube glabrous. Leaves usually remain green on drying … 9.

9. Anthers white both when fresh and dried … 10.

– Anthers not white … 11.

10. Root system mostly of primary and secondary roots. … sect. *Lamprosantha*

– Root system mostly of adventitious roots … sect. *Eremopsyllium*

11. Corolla lobes longer than 1.5 mm. Ovary with four or fewer ovules. Leaf width usually less than 25 mm … sect. *Pacifica*

– Corolla lobes shorter than 1.5 mm. Ovary usually with four or more ovules. Leaf width more than 25 mm … 12.

12. Anterior sepals distinctly narrower than posterior, and differently shaped … sect. *Leptostachys*

– Anterior and posterior sepals similar … sect. *Plantago*

13. Leaves opposite or in whorls of three … 14.

– Leaves alternate … 15.

14. Perennials, typically without long glandular hairs. Inner bracts narrow. Seeds longer than 3 mm … sect. *Arborescens*

– Annuals, with long glandular hairs. Inner bracts are broad. Seeds shorter than 3 mm … sect. *Psyllium*

15. Bract with the upper part scarious, acuminate. Some species with anterior sepals united for more than half of their length … sect. *Lancifolia*

– Bract without scarious, acuminate upper part. Anterior sepals always free … 16.

17. Connective of anther very large, about as long as the pollen sacs. Plants densely hairy (leaf surface hardly visible), cells of non-glandular hairs jointed by a common wall with crown-like elongations … sect. *Hymenopsyllium*

– Connective of anther smaller. Plants are not densely hairy, cells of hairs without crown-shape elongations … 18.

18. The nerve of anterior sepals well developed. Corolla lobes are slightly hairy on the back. The concave inner side of the seed covered by a ragged, white membrane, except for two areas to the right and left of the center. Leaves are usually remaining green on drying … sect. *Albicans*

*–* The nerve of anterior sepals present at base only, distal part scarious. Corolla lobes not hairy. White membrane on seeds absent. Leaves usually darken on drying. … sect. *Montana*

## ACKNOWLEDGMENTS

We are grateful to the curators of B, BO, BRIT, C, CAS, COL, CONC, F, HUH, IBSC, JBB, KW, LE, M, MA, MHA, MO, MW, NBG/SAM, NY, PE, PRE, SGO, SP, SPF, TI, TNS, UC/JEPS, US, USM herbarium collections (Thiers, 2019) for the permission to work with their herbarium material, and to Barcoding of Life Consortium and personally to Maria L. Kuzmina (University of Guelph, Guelph, Ontario, Canada) for the help with sequencing North American species of *Plantago*. Our special thanks to Polina Volkova, Jim Keesling and Gabor Sramkó for the help with *Plantago* samples, and Ekaterina Shipunova and the staff of the Subdirección Científica, Jardín Botánico de Bogotá José Celestino Mutis (Bogotá, Colombia) for the help with *Aragoa* samples. We thank the Department of Biology of Minot State University for the financial support, and Minot State University students for the help with morphometric observations. From May 2014, this research was supported by North Dakota INBRE.

## SUPPORT MATERIALS

Support Table 1. Working classification of *Plantagineae*.

Support Table 2. Vouchers of *Plantagineae* samples. Samples used for morphometric measurements labeled with star* (please note that not all measured samples yielded the DNA of the reliable quality).

Support Table 3. GenBank accession numbers of *Plantagineae* samples.

Support Table 4. Binary morphological characters used.

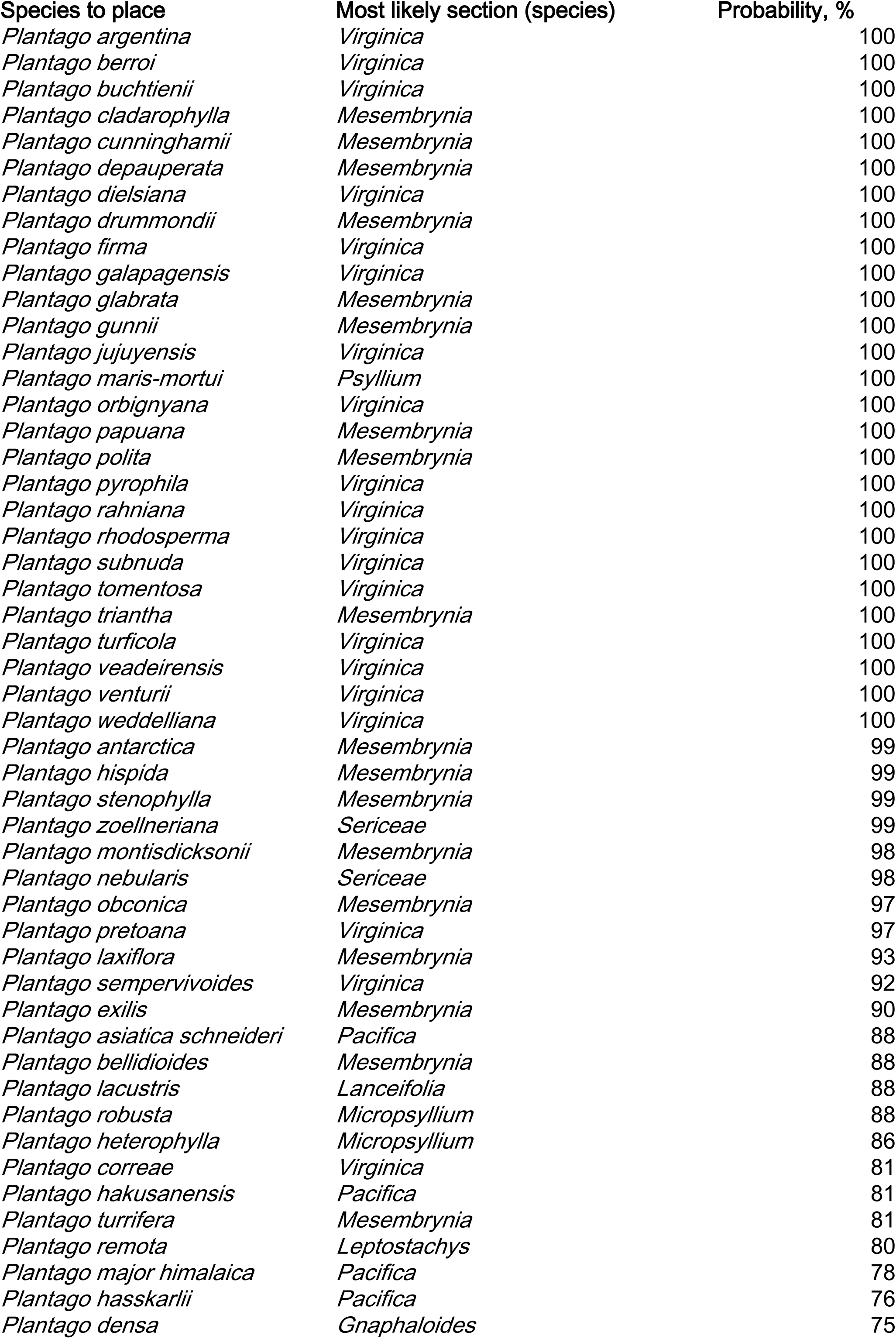
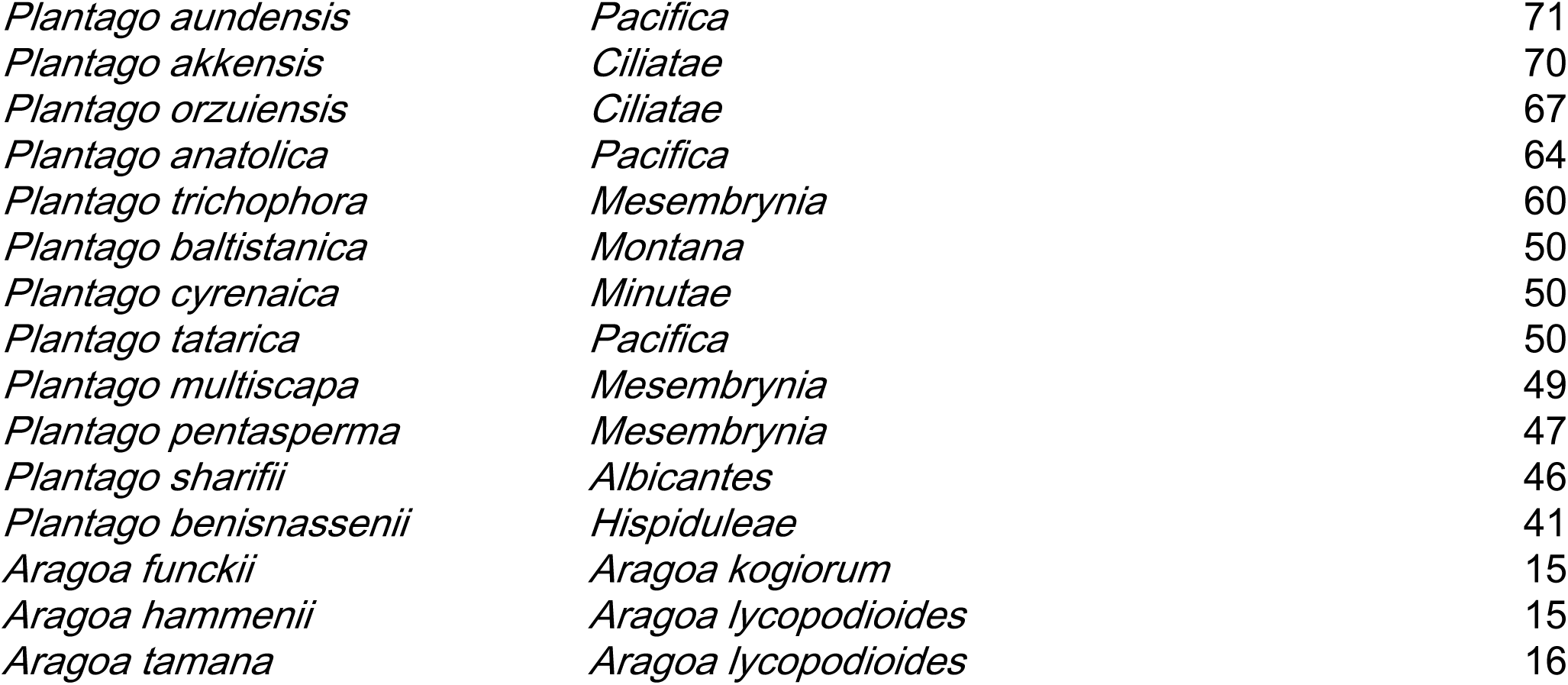

| Species to place | Most likely section (species) | Probability, % |
| --- | --- | --- |
| *Plantago argentina* | *Virginica* | 100 |
| *Plantago berroi* | *Virginica* | 100 |
| *Plantago buchtienii* | *Virginica* | 100 |
| *Plantago cladarophylla* | *Mesembrynia* | 100 |
| *Plantago cunninghamii* | *Mesembrynia* | 100 |
| *Plantago depauperata* | *Mesembrynia* | 100 |
| *Plantago dielsiana* | *Virginica* | 100 |
| *Plantago drummondii* | *Mesembrynia* | 100 |
| *Plantago firma* | *Virginica* | 100 |
| *Plantago galapagensis* | *Virginica* | 100 |
| *Plantago glabrata* | *Mesembrynia* | 100 |
| *Plantago gunnii* | *Mesembrynia* | 100 |
| *Plantago jujuyensis* | *Virginica* | 100 |
| *Plantago maris-mortui* | *Psyllium* | 100 |
| *Plantago orbignyana* | *Virginica* | 100 |
| *Plantago papuana* | *Mesembrynia* | 100 |
| *Plantago polita* | *Mesembrynia* | 100 |
| *Plantago pyrophila* | *Virginica* | 100 |
| *Plantago rahniana* | *Virginica* | 100 |
| *Plantago rhodosperma* | *Virginica* | 100 |
| *Plantago subnuda* | *Virginica* | 100 |
| *Plantago tomentosa* | *Virginica* | 100 |
| *Plantago triantha* | *Mesembrynia* | 100 |
| *Plantago turficola* | *Virginica* | 100 |
| *Plantago veadeirensis* | *Virginica* | 100 |
| *Plantago venturii* | *Virginica* | 100 |
| *Plantago weddelliana* | *Virginica* | 100 |
| *Plantago antarctica* | *Mesembrynia* | 99 |
| *Plantago hispida* | *Mesembrynia* | 99 |
| *Plantago stenophylla* | *Mesembrynia* | 99 |
| *Plantago zoellneriana* | *Sericeae* | 99 |
| *Plantago montisdicksonii* | *Mesembrynia* | 98 |
| *Plantago nebularis* | *Sericeae* | 98 |
| *Plantago obconica* | *Mesembrynia* | 97 |
| *Plantago pretoana* | *Virginica* | 97 |
| *Plantago laxiflora* | *Mesembrynia* | 93 |
| *Plantago sempervivoides* | *Virginica* | 92 |
| *Plantago exilis* | *Mesembrynia* | 90 |
| *Plantago asiatica schneideri* | *Pacifica* | 88 |
| *Plantago bellidioides* | *Mesembrynia* | 88 |
| *Plantago lacustris* | *Lanceifolia* | 88 |
| *Plantago robusta* | *Micropsyllium* | 88 |
| *Plantago heterophylla* | *Micropsyllium* | 86 |
| *Plantago correae* | *Virginica* | 81 |
| *Plantago hakusanensis* | *Pacifica* | 81 |
| *Plantago turrifera* | *Mesembrynia* | 81 |
| *Plantago remota* | *Leptostachys* | 80 |
| *Plantago major himalaica* | *Pacifica* | 78 |
| *Plantago hasskarlii* | *Pacifica* | 76 |
| *Plantago densa* | *Gnaphaloides* | 75 |
| *Plantago aundensis* | *Pacifica* | 71 |
| *Plantago akkensis* | *Ciliatae* | 70 |
| *Plantago orzuiensis* | *Ciliatae* | 67 |
| *Plantago anatolica* | *Pacifica* | 64 |
| *Plantago trichophora* | *Mesembrynia* | 60 |
| *Plantago baltistanica* | *Montana* | 50 |
| *Plantago cyrenaica* | *Minutae* | 50 |
| *Plantago tatarica* | *Pacifica* | 50 |
| *Plantago multiscapa* | *Mesembrynia* | 49 |
| *Plantago pentasperma* | *Mesembrynia* | 47 |
| *Plantago sharifii* | *Albicantes* | 46 |
| *Plantago benisnassenii* | *Hispiduleae* | 41 |
| *Aragoa funckii* | *Aragoa kogiorum* | 15 |
| *Aragoa hammenii* | *Aragoa lycopodioides* | 15 |
| *Aragoa tamana* | *Aragoa lycopodioides* | 16 |

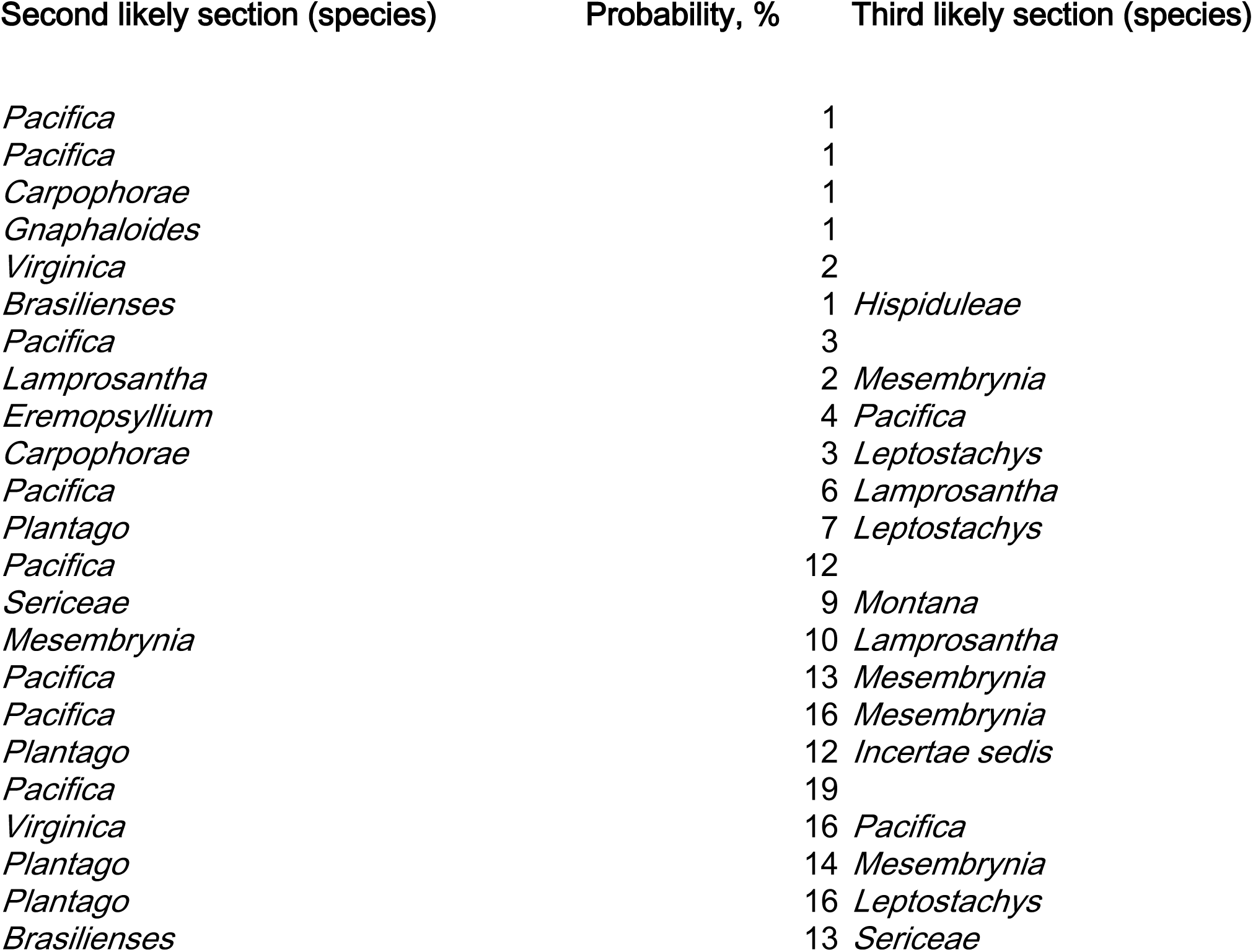
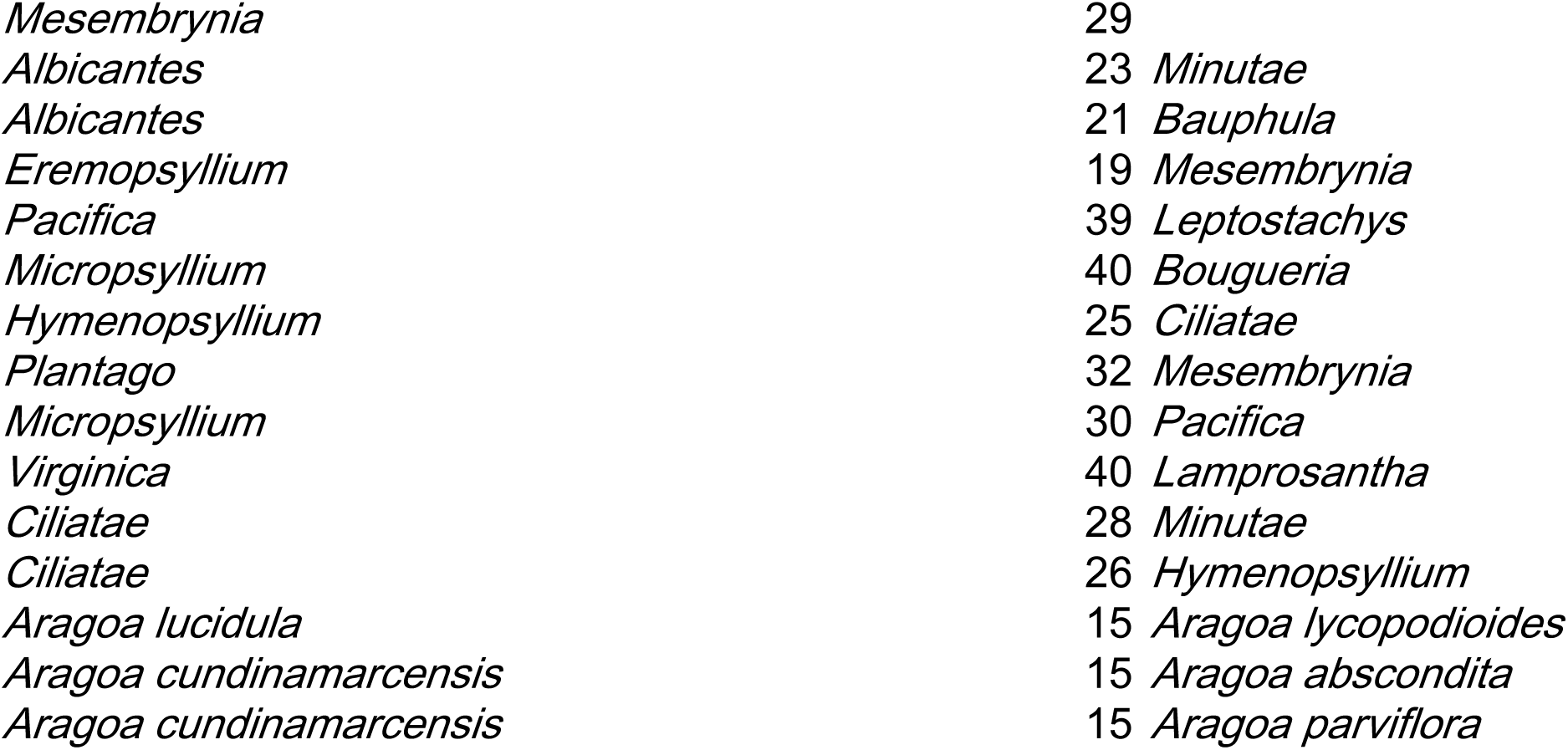

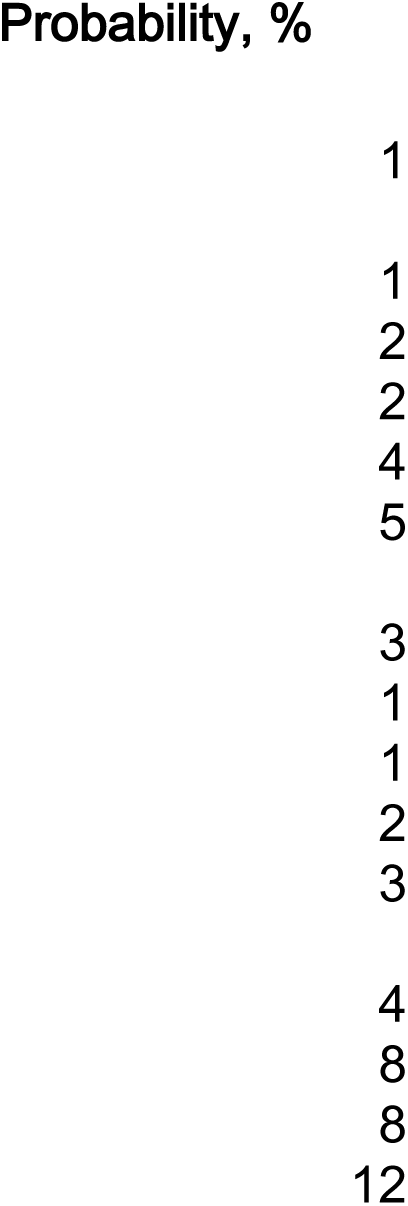
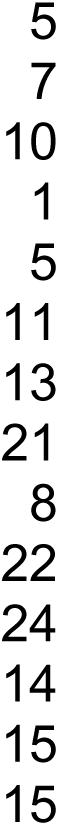

