## Supplementary Tables and Figures for "How to map a plantain: phylogeny of the diverse *Plantagineae* (Lamiales)": photographs_of_plants_sampled_in_the_field.pdf

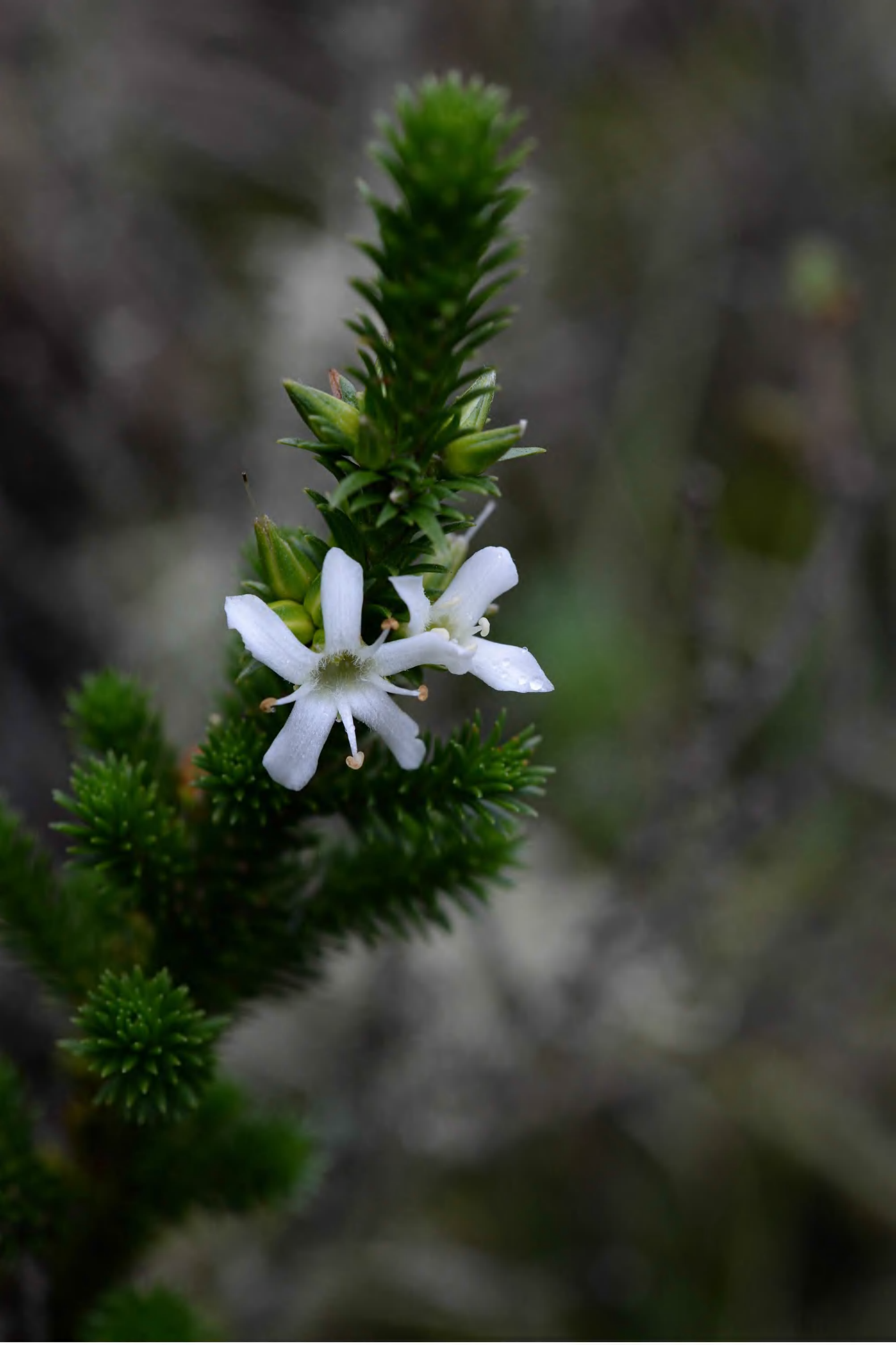

aragoa_abietina_p_1273_abs_9406.jpg

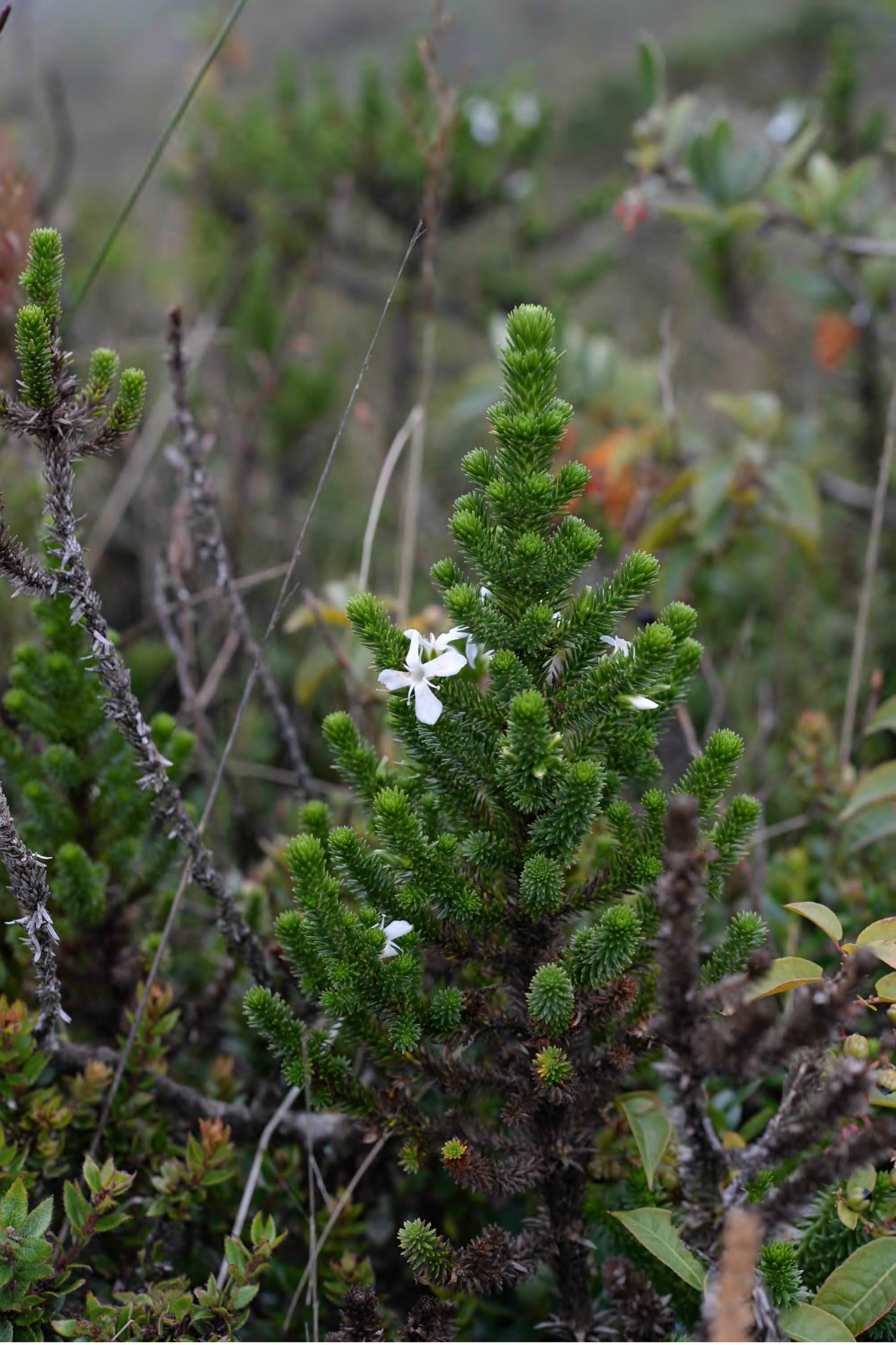

aragoa_abietina_p_1280_abs_9609.jpg

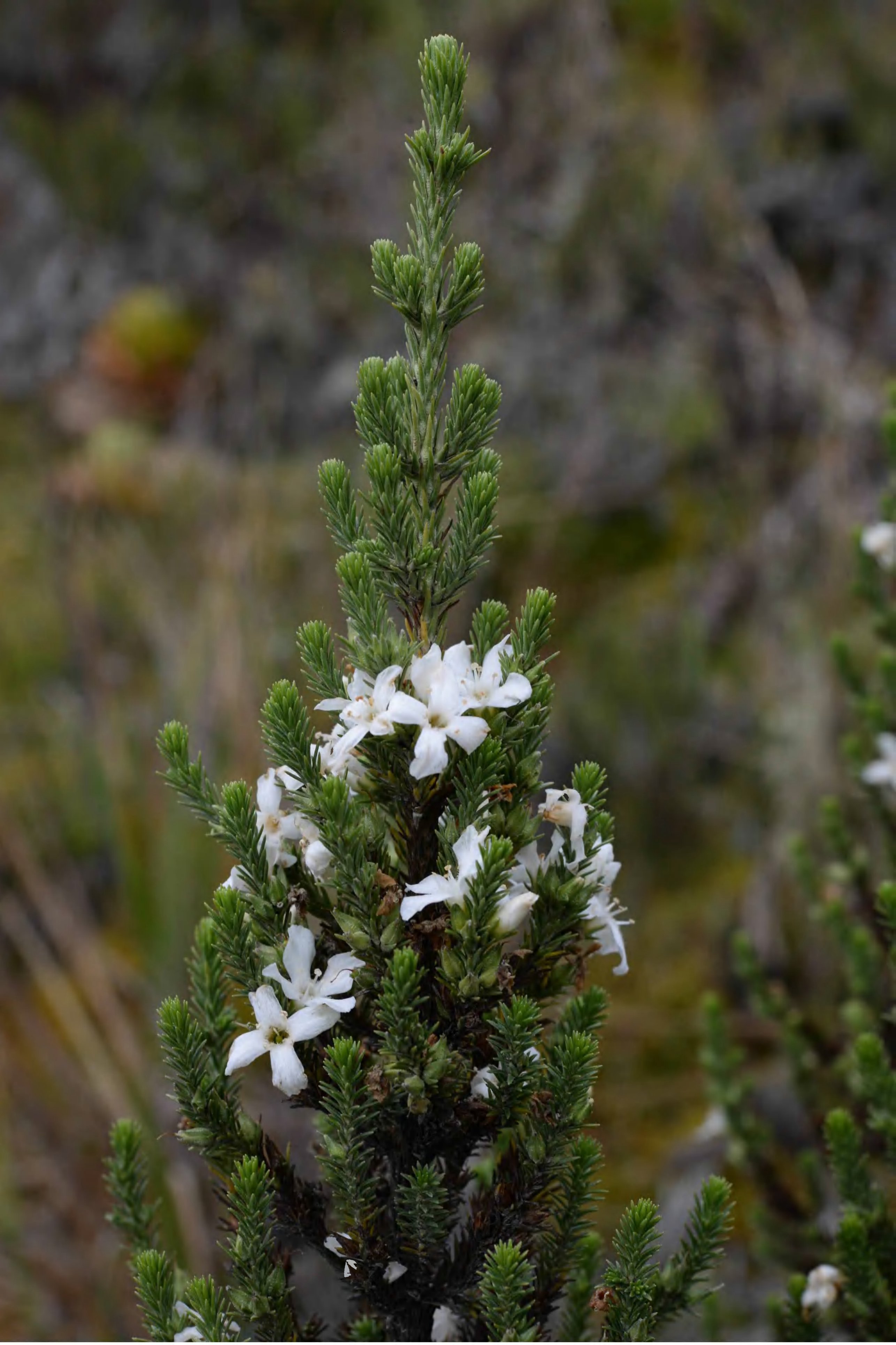

*aragoa_corrugatifolia_p_1282_bbs_0134.jpg*

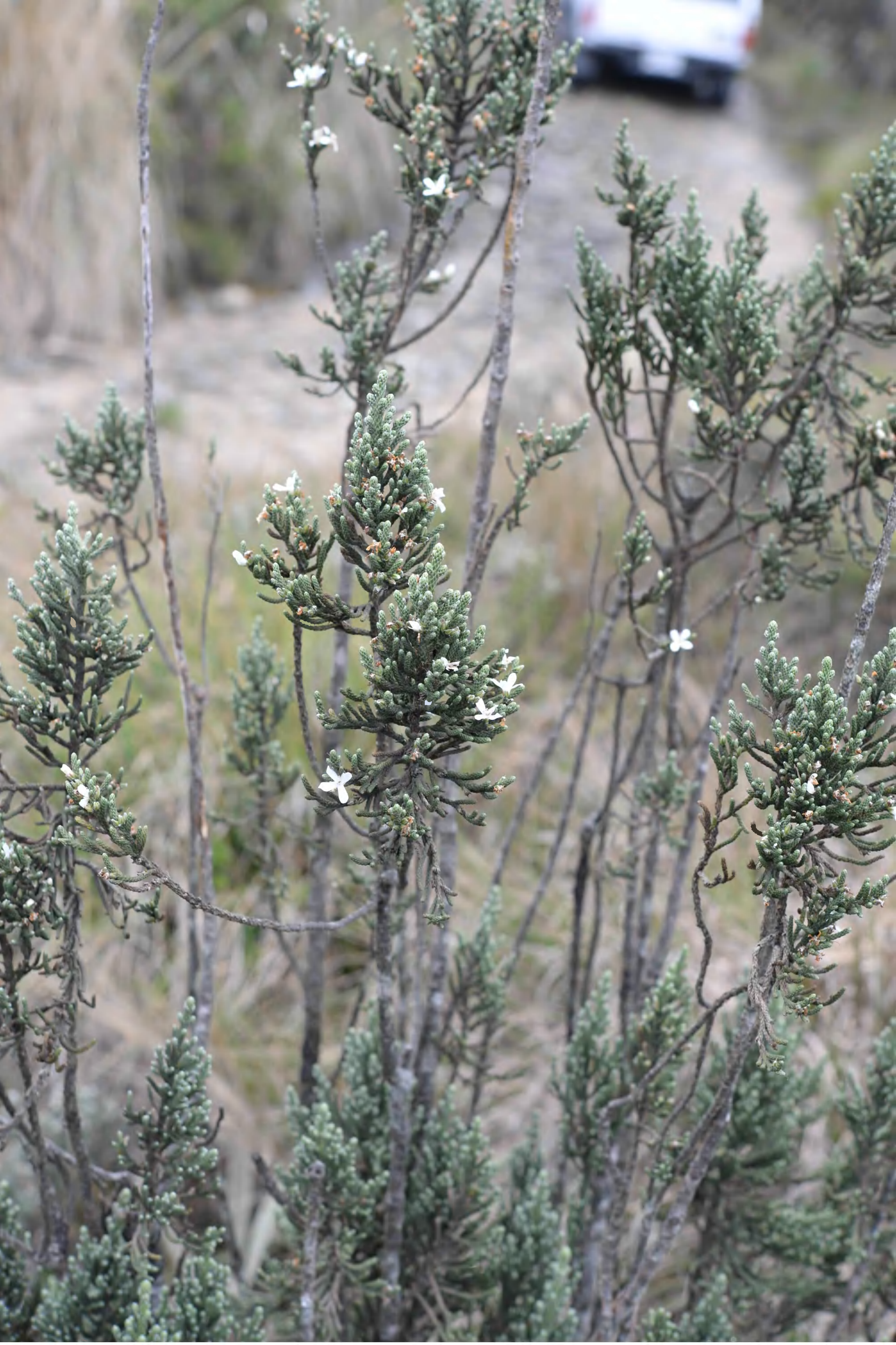

aragoa_cupressina_p_1275_abs_9587.jpg

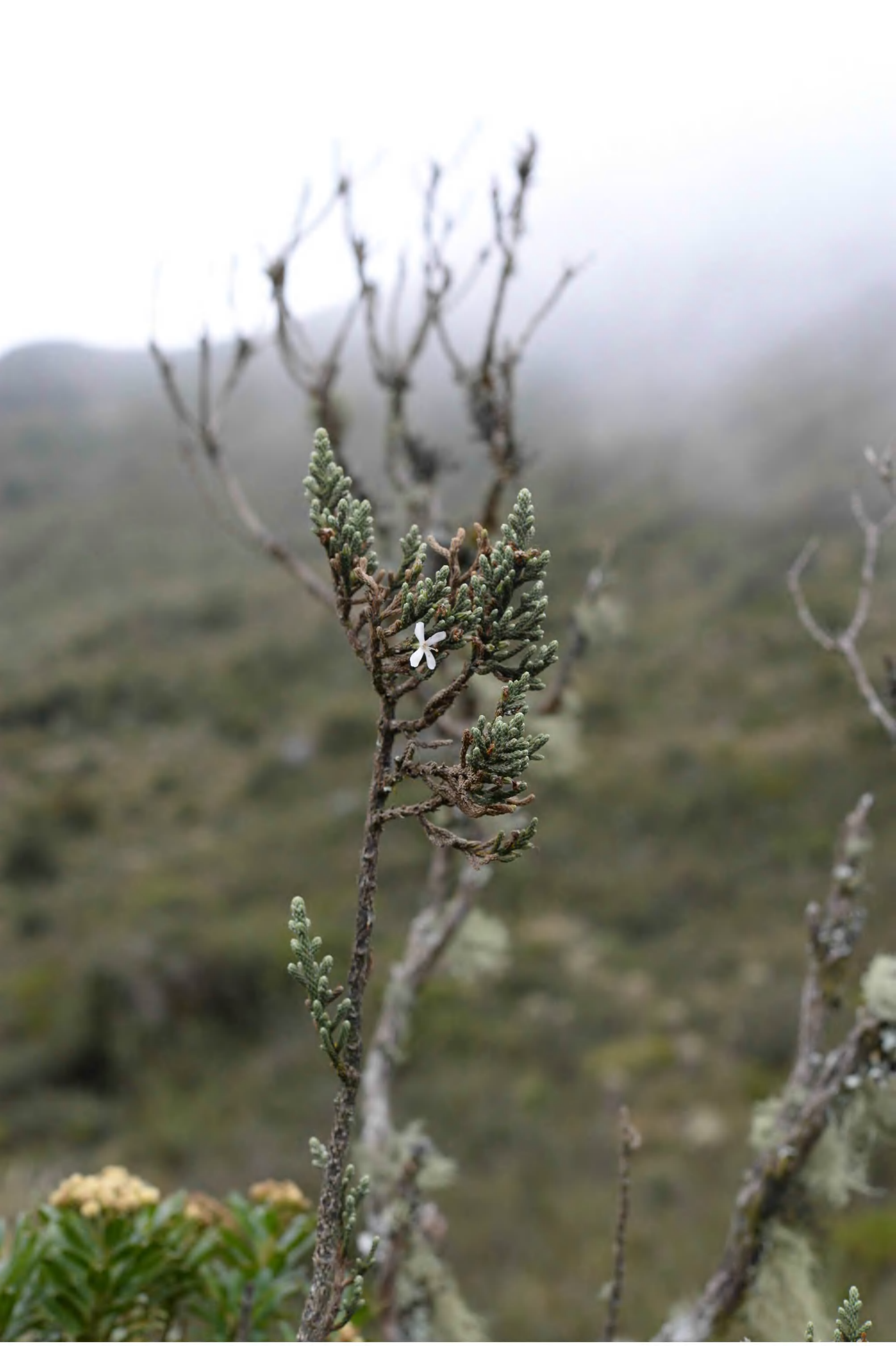

*aragoa_cupressina_p_1276_abs_9705.jpg*

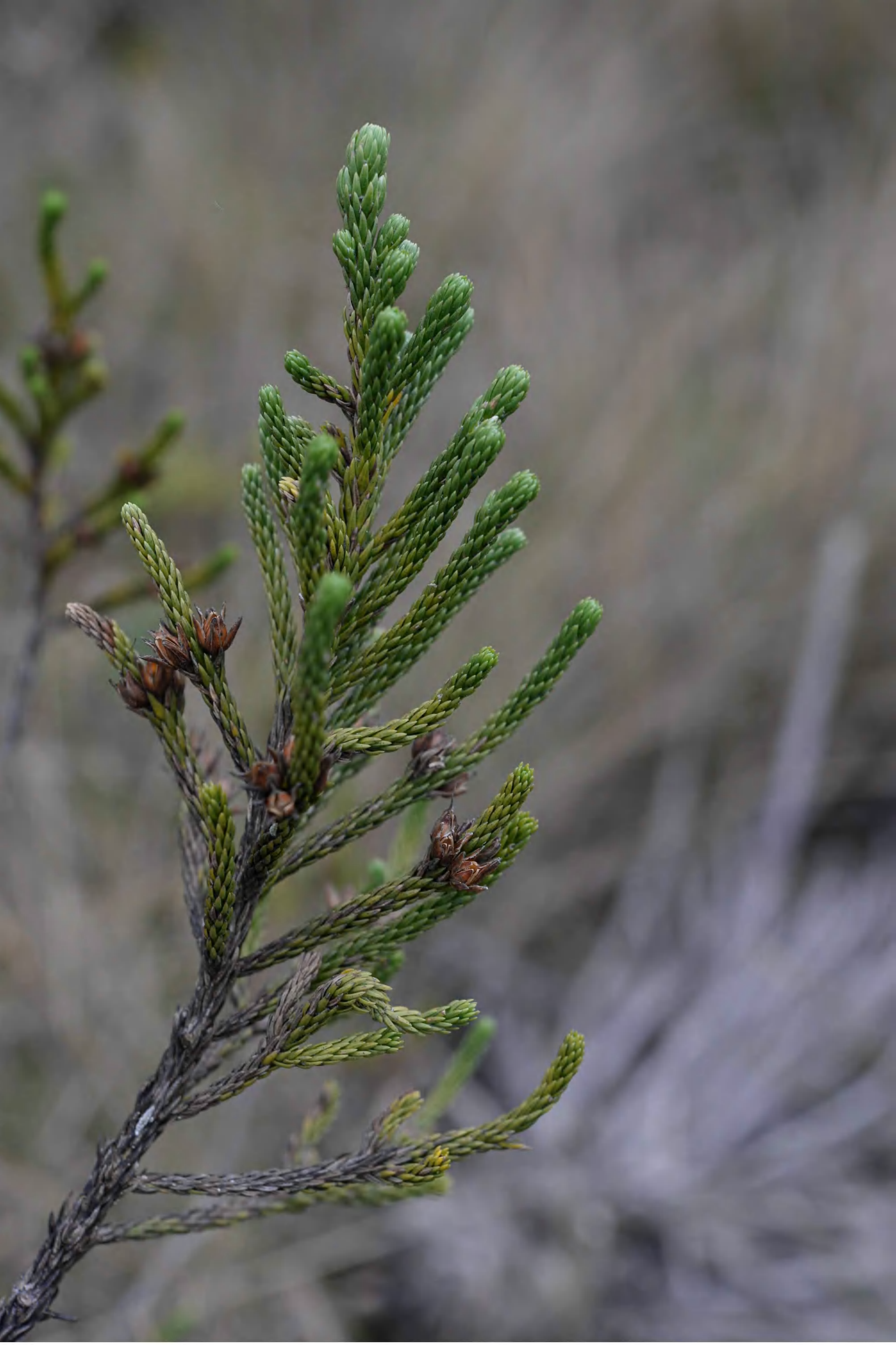

aragoa_jaramilloi_p_1277_abs_9630.jpg

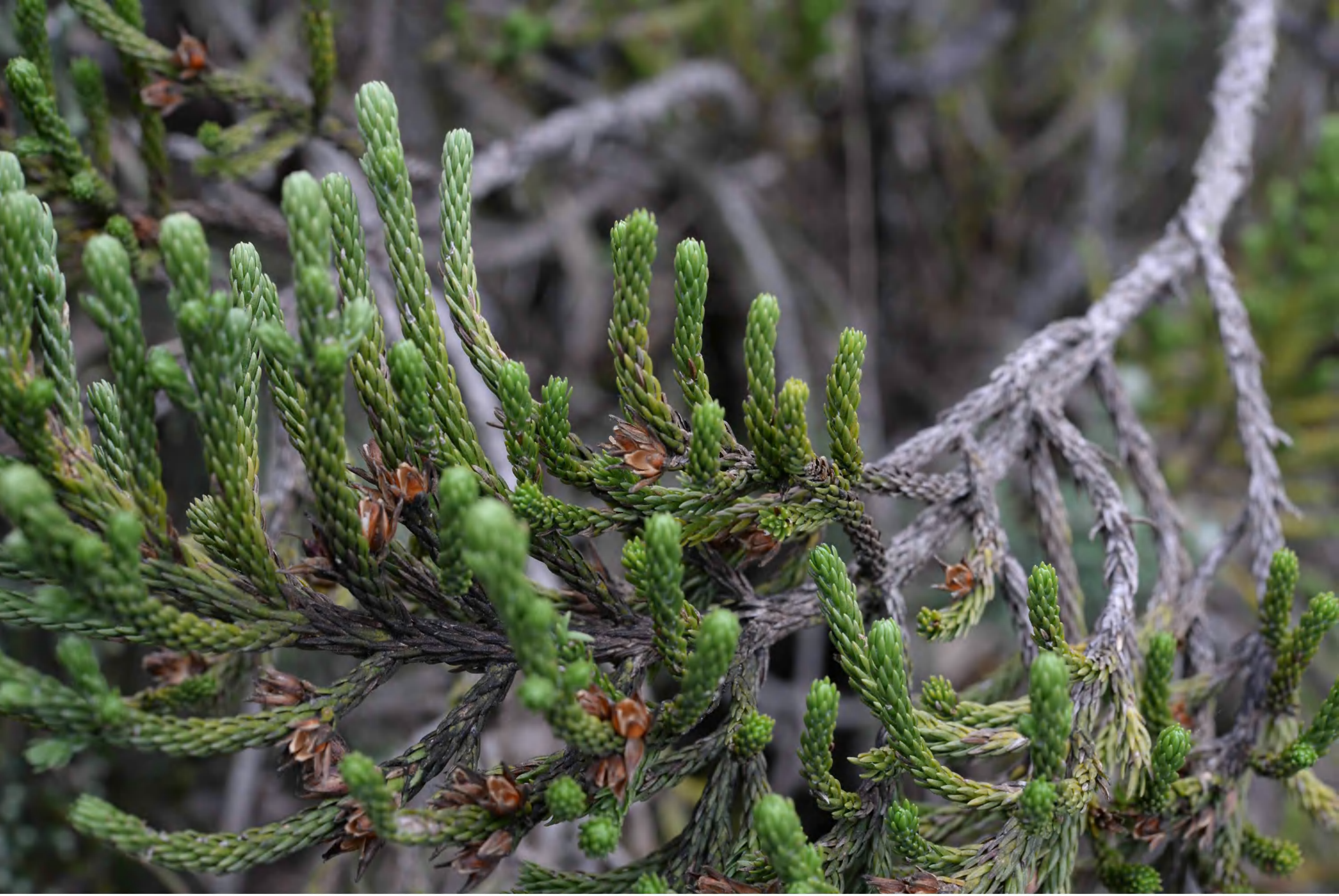

*aragoa_jaramilloi_p_1278_abs_9880.jpg*

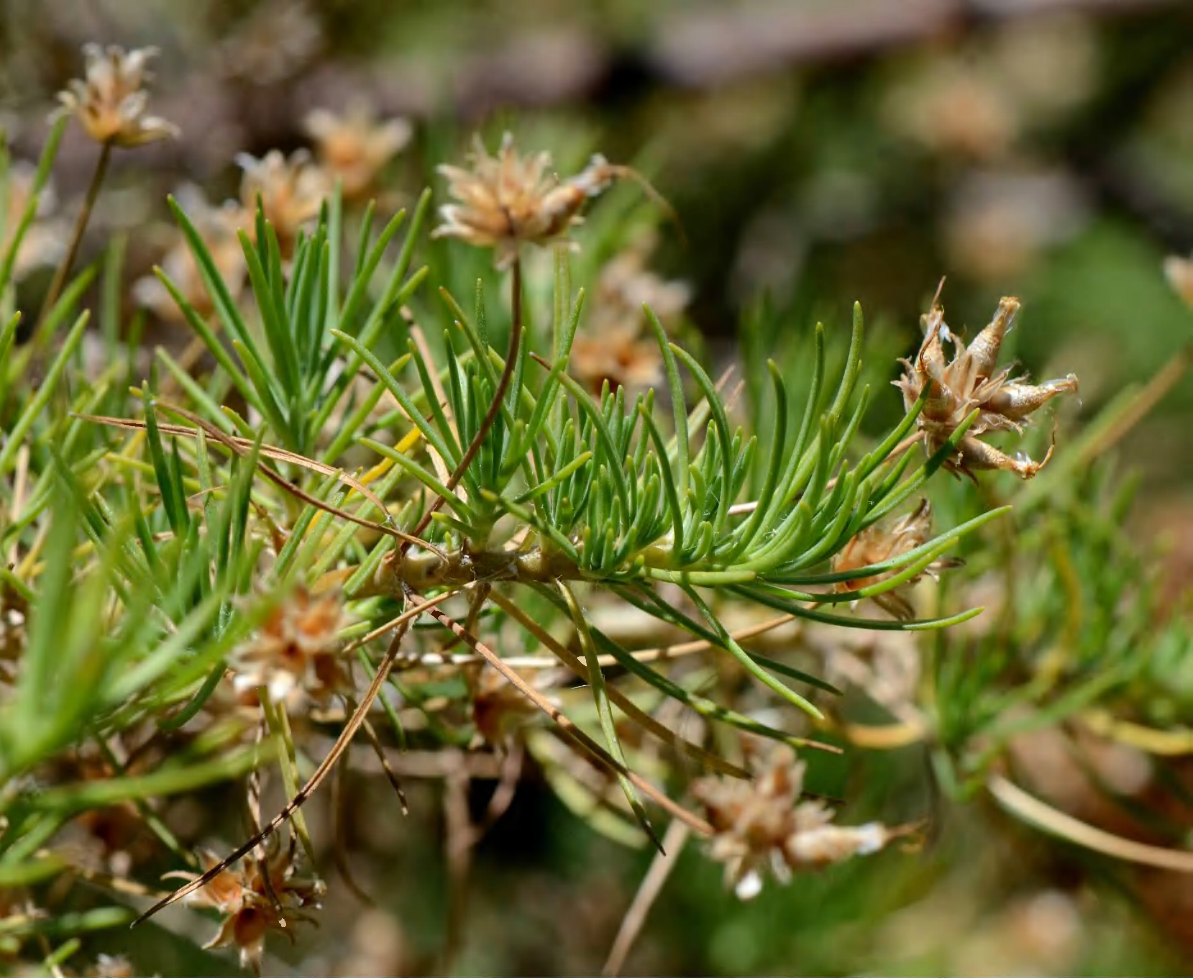

plantago_arborescens_p_0000_bbb_0218.jpg

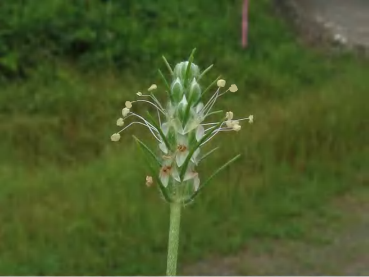

plantago_aristata_p_1
501_image016.jpg

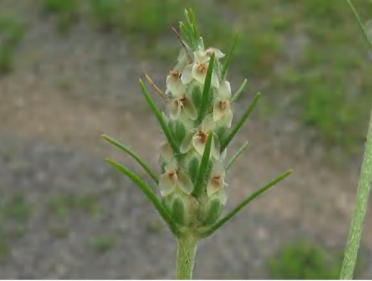

plantago_aristata_p_1502_image017.jpg

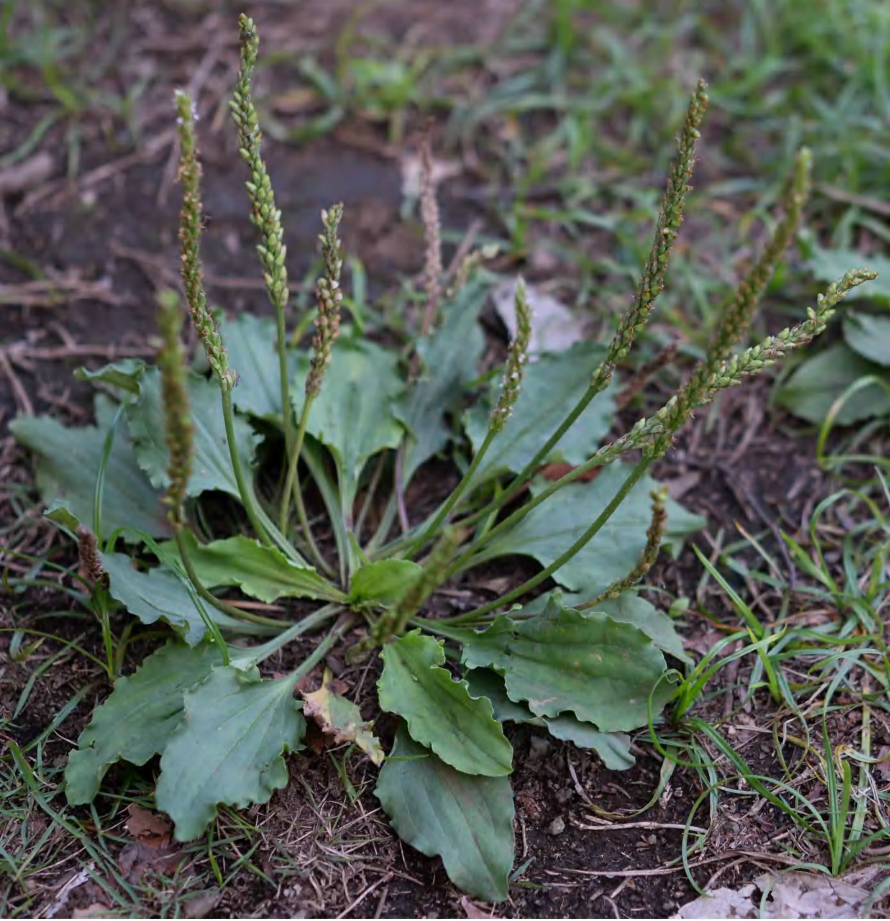

*plantago_asiatica_p_1503_abs_0276.jpg*

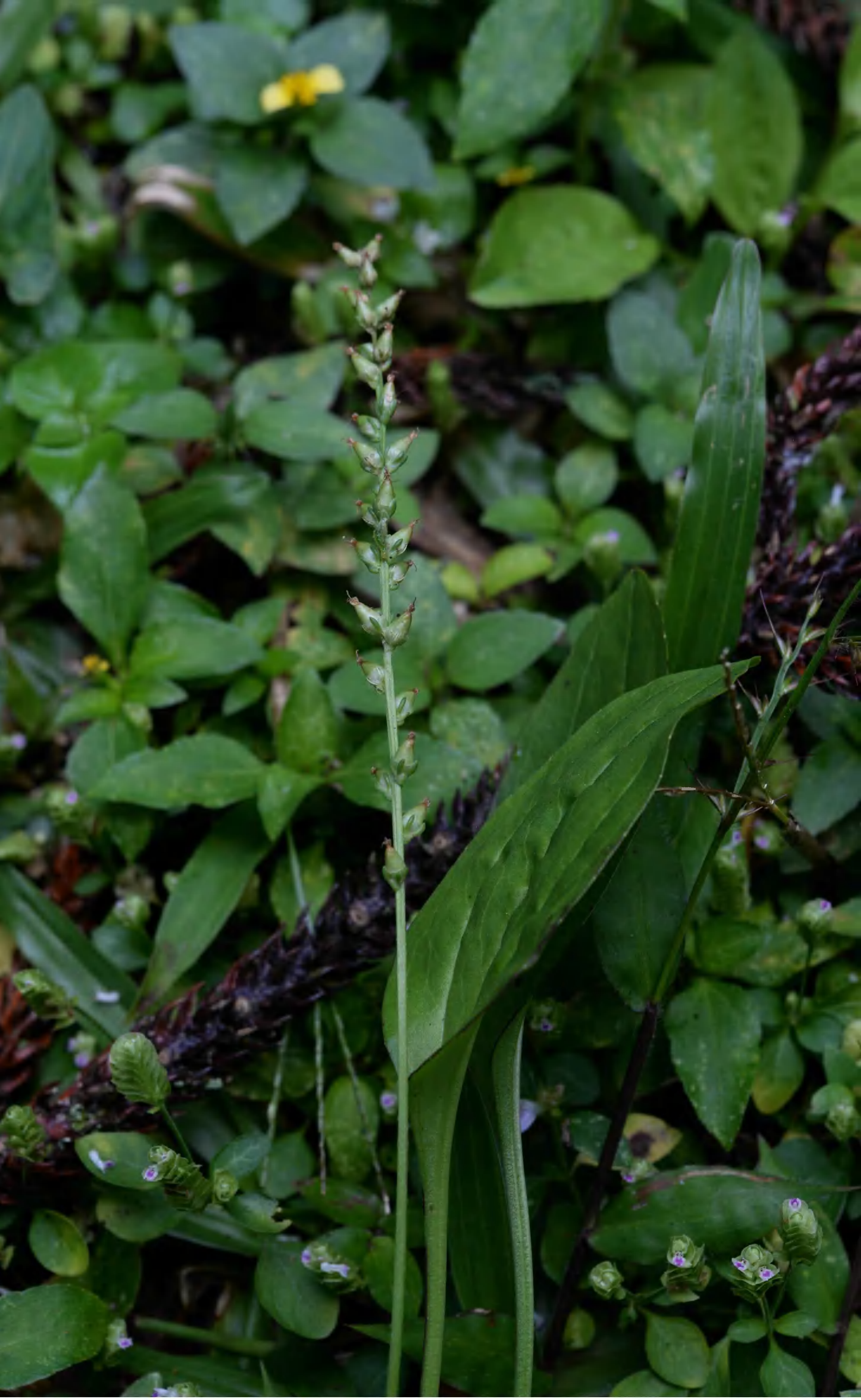

plantago_asiatica_p_1504_abs_0656.jpg

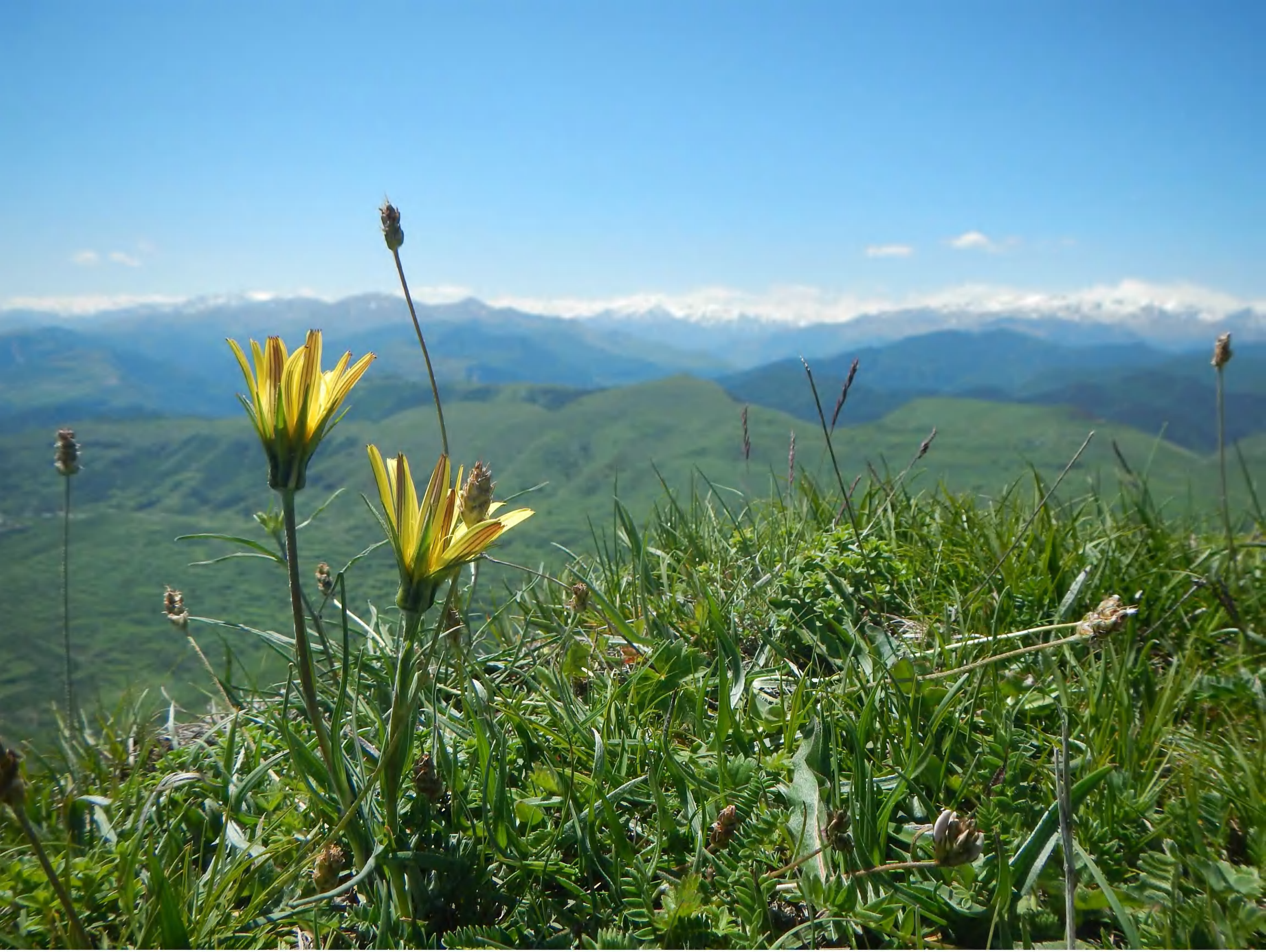

plantago_atrata_p_1506_2.jpg

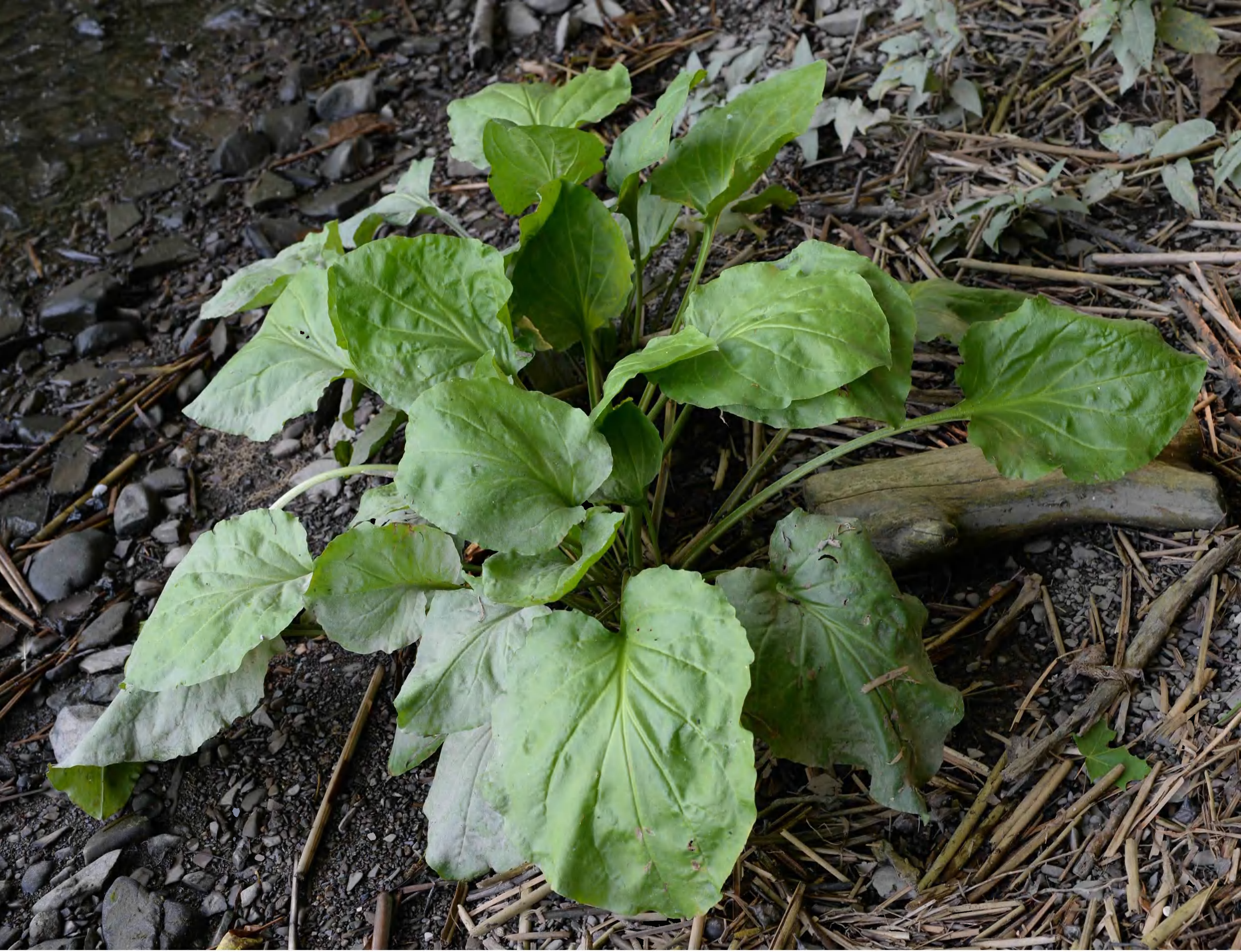

plantago_cordata_p_1507_abs_3028.jpg

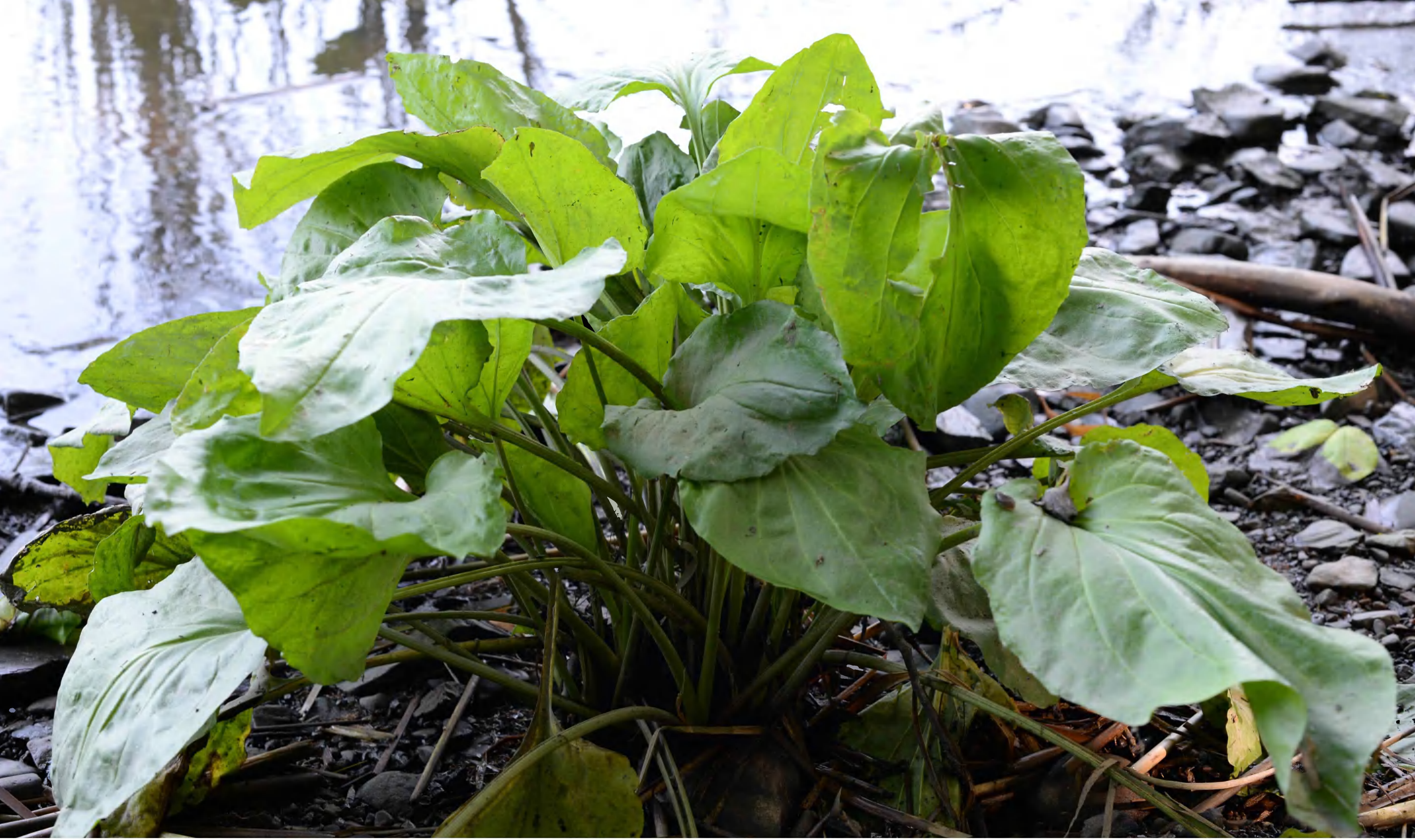

plantago_cordata_p_1508_abs_3057.jpg

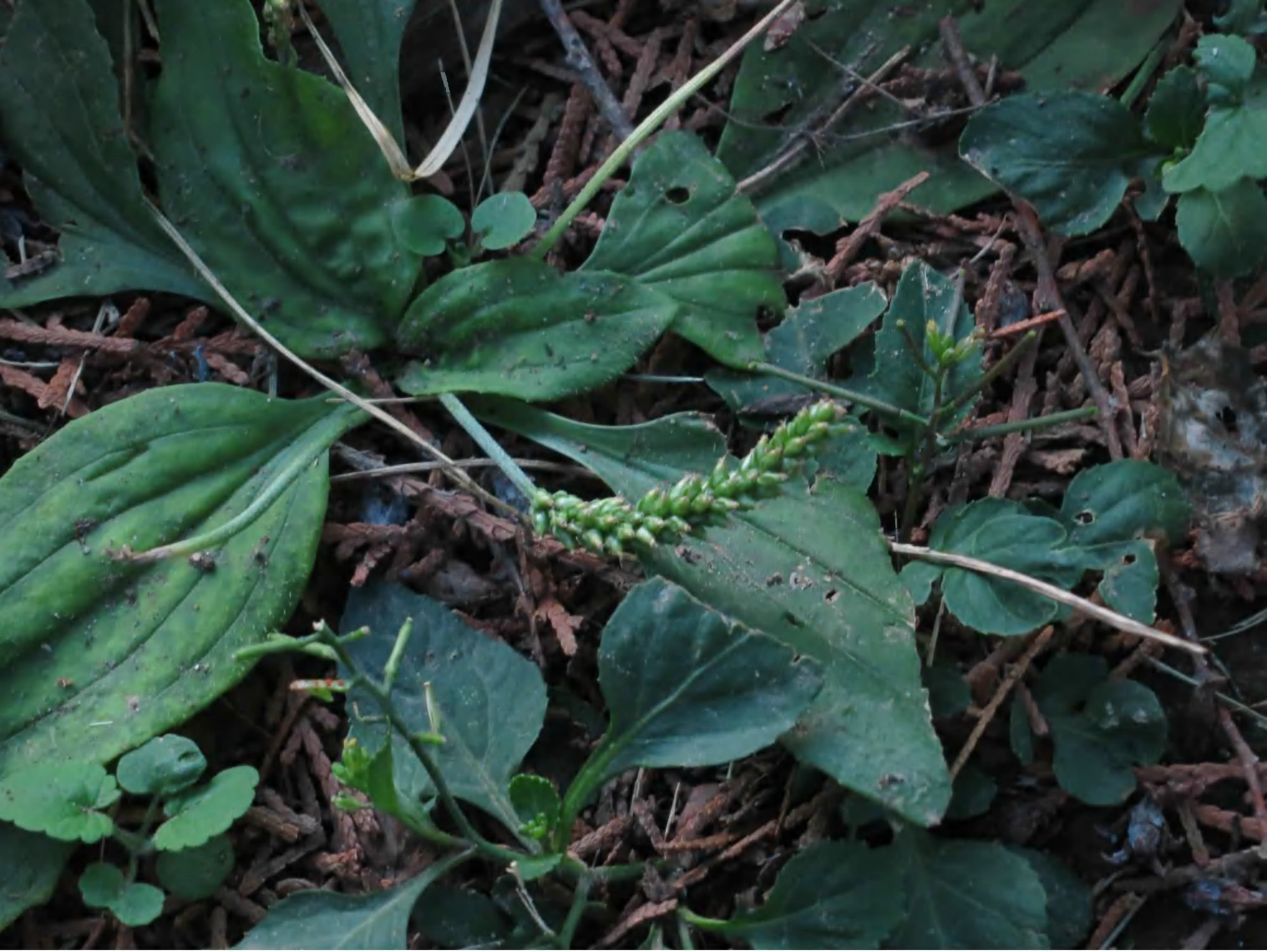

*plantago_depressa_p_1505_img_4897.jpg*

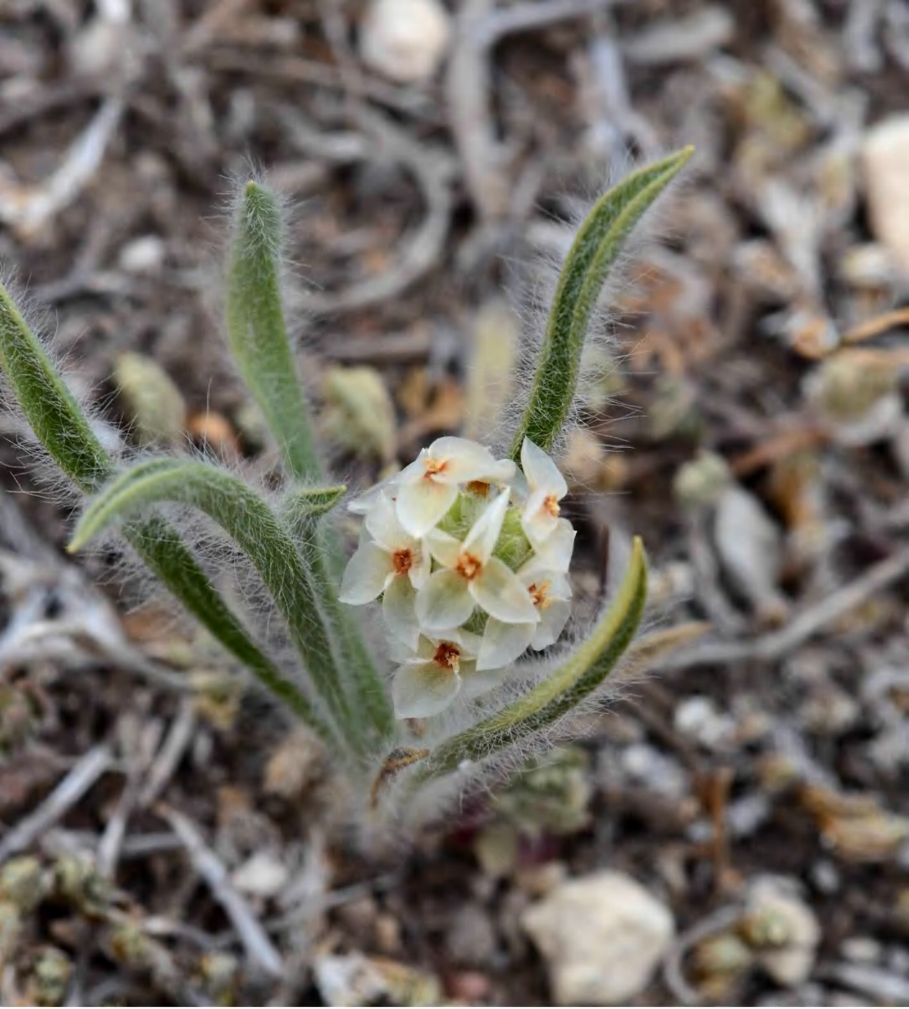

plantago_hellerii_p_0739_bbb_8046.jpg

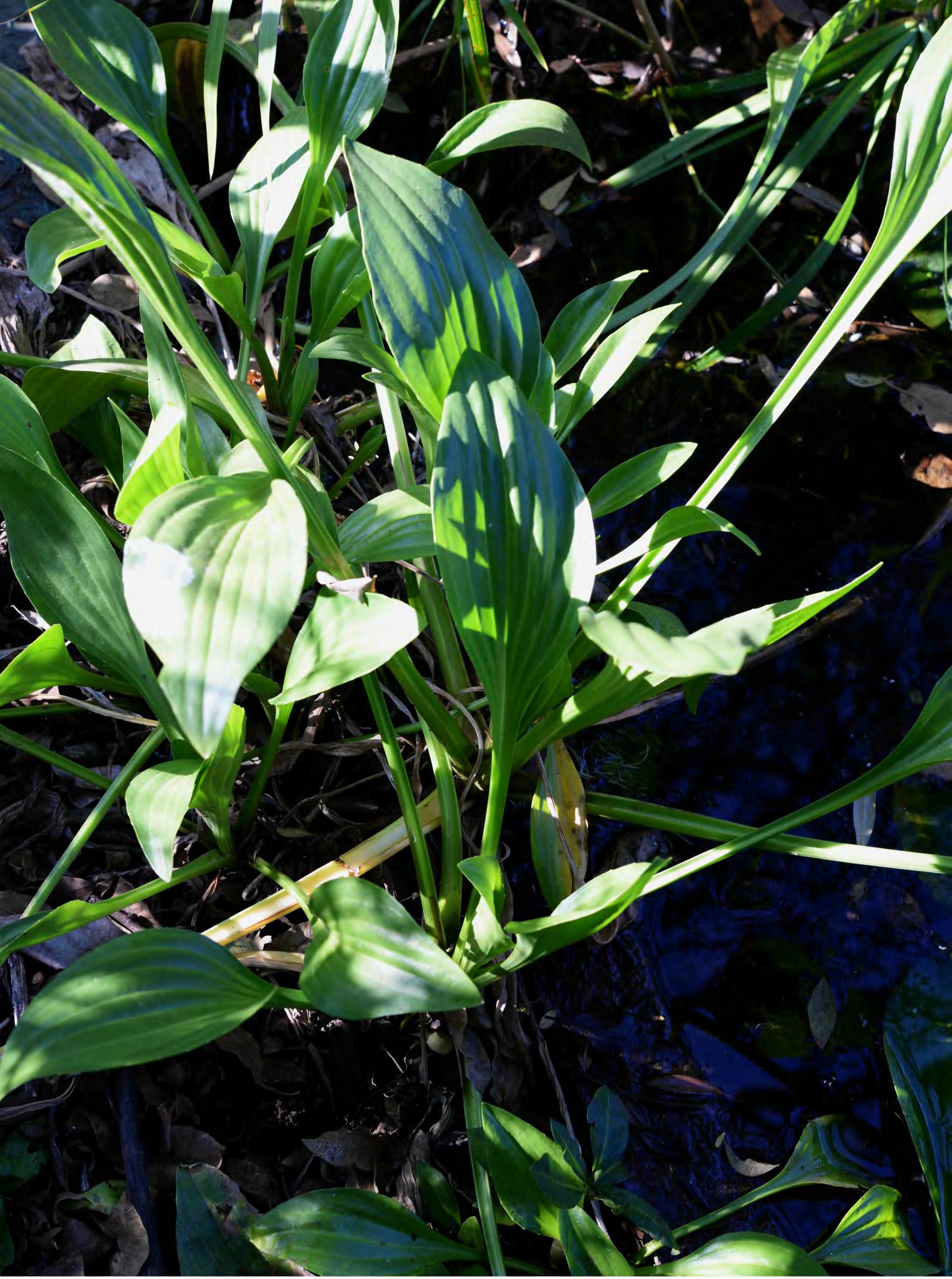

plantago_longissima_p_1121_abs_2297.jpg

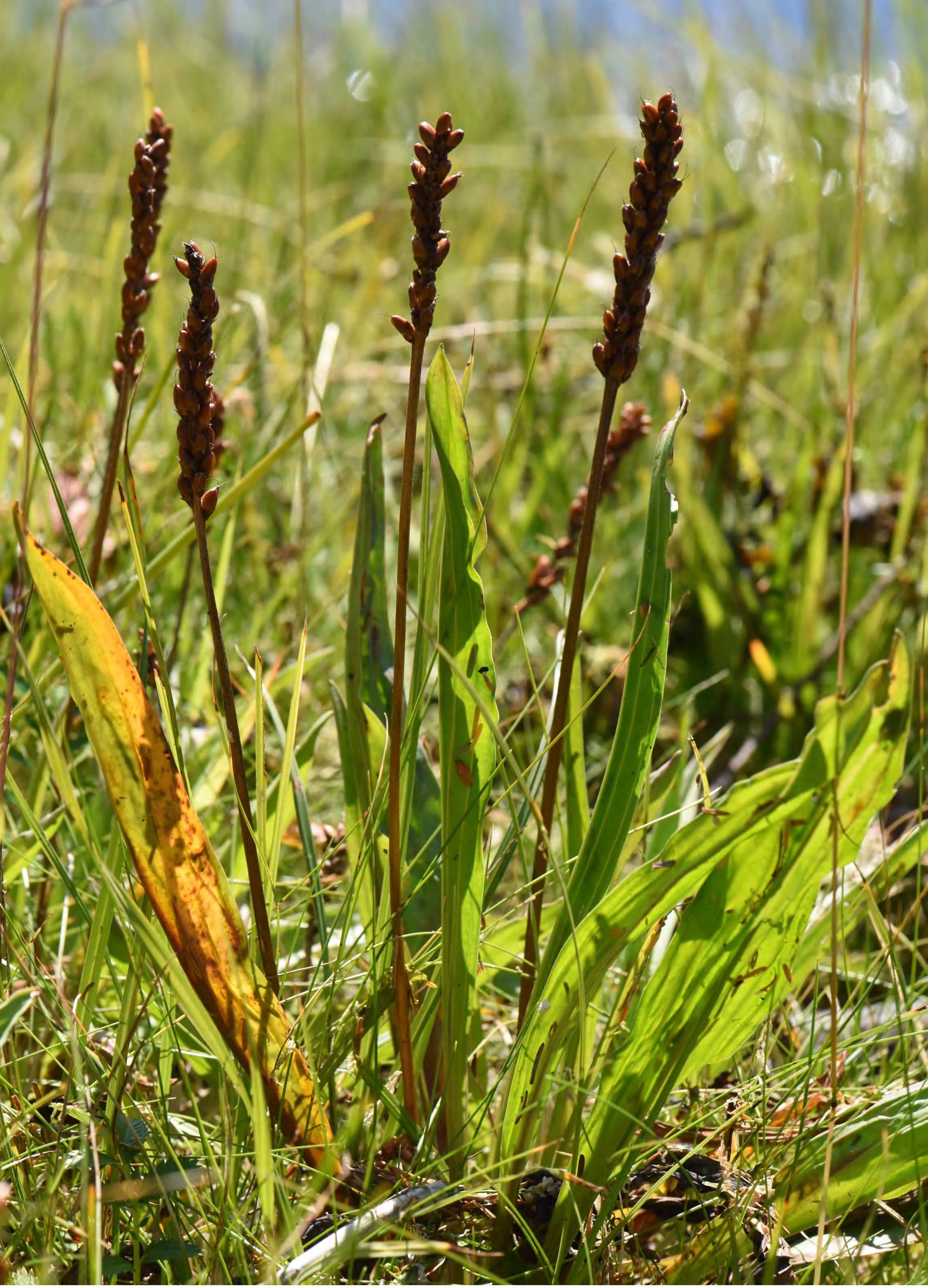

plantago_macrocarpa_p_1509_ycv_3522.jpg

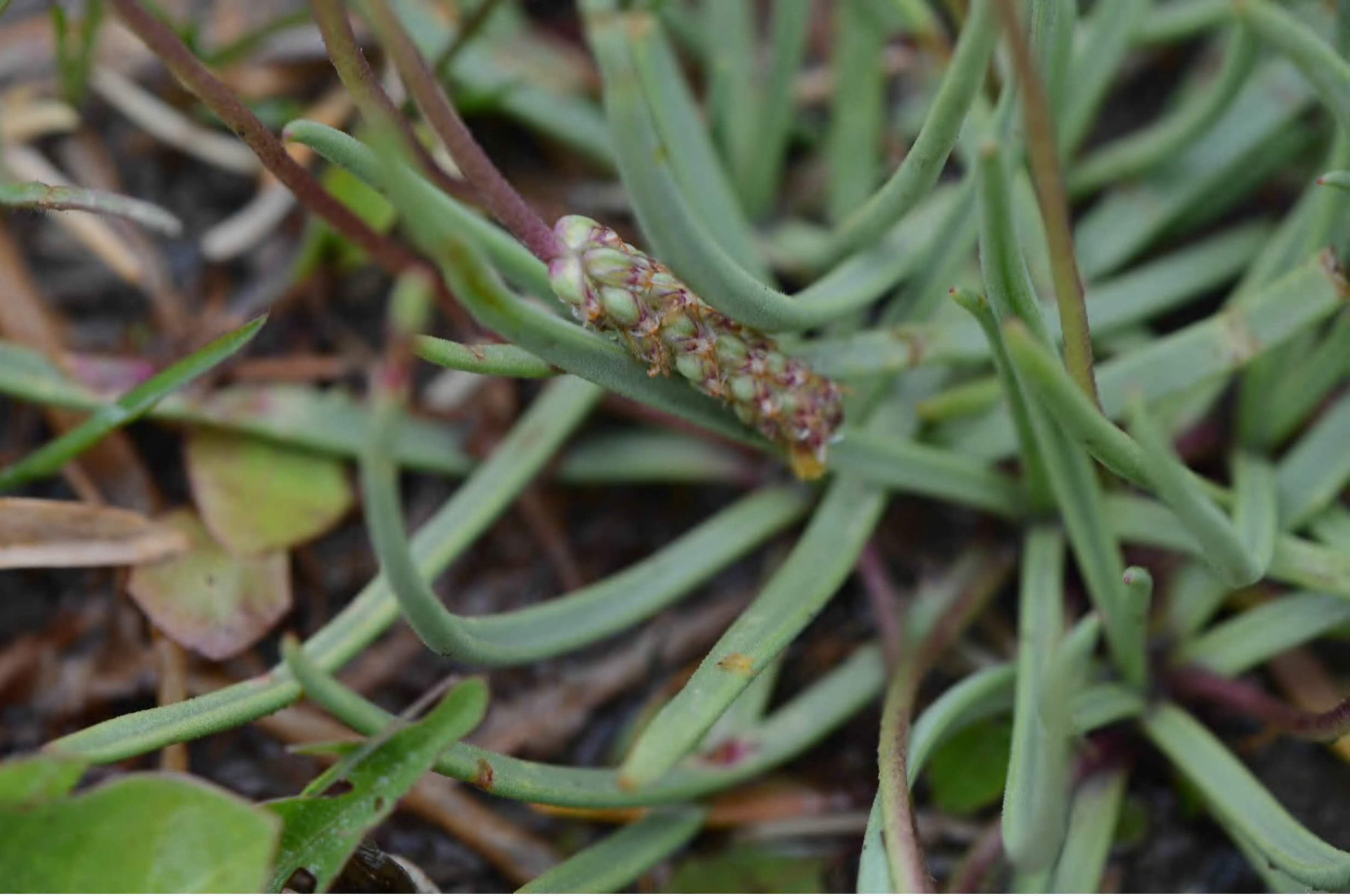

plantago_maritima_p_1203_bbb_2425.jpg

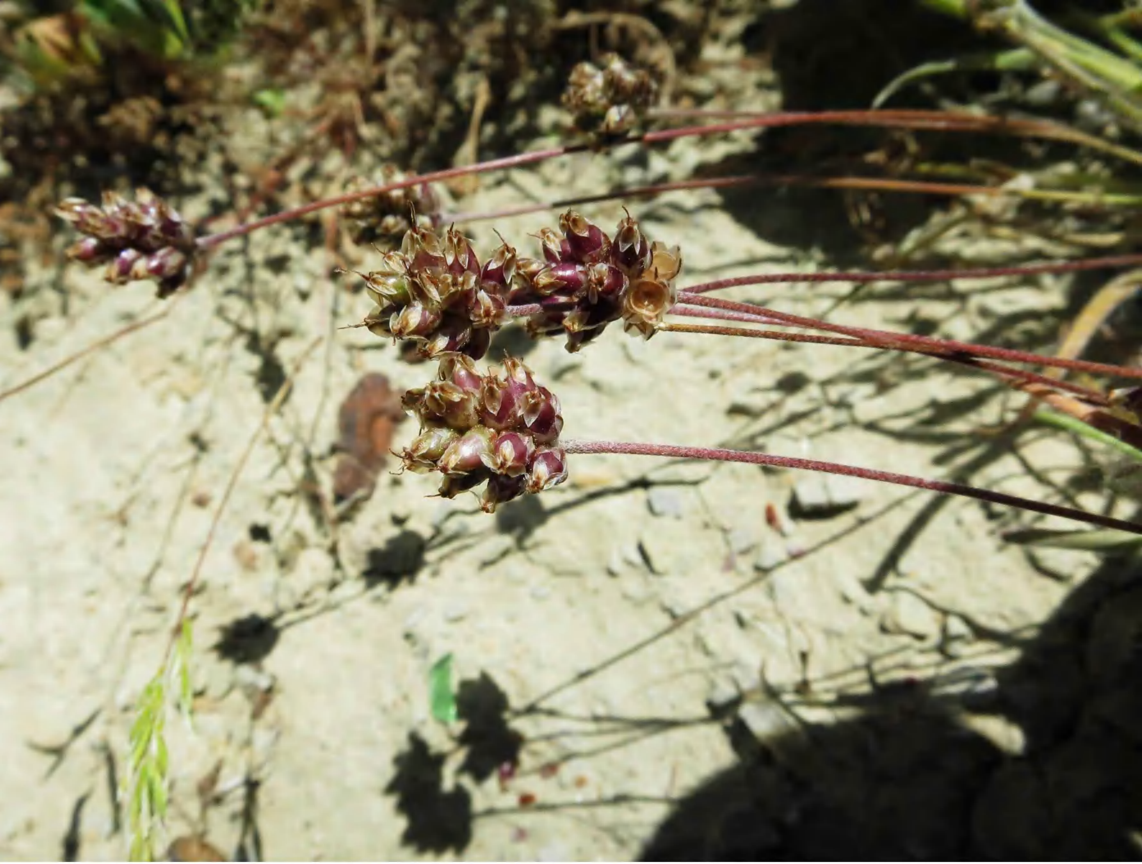

plantago_ovata_p_1271_img_1150.jpg

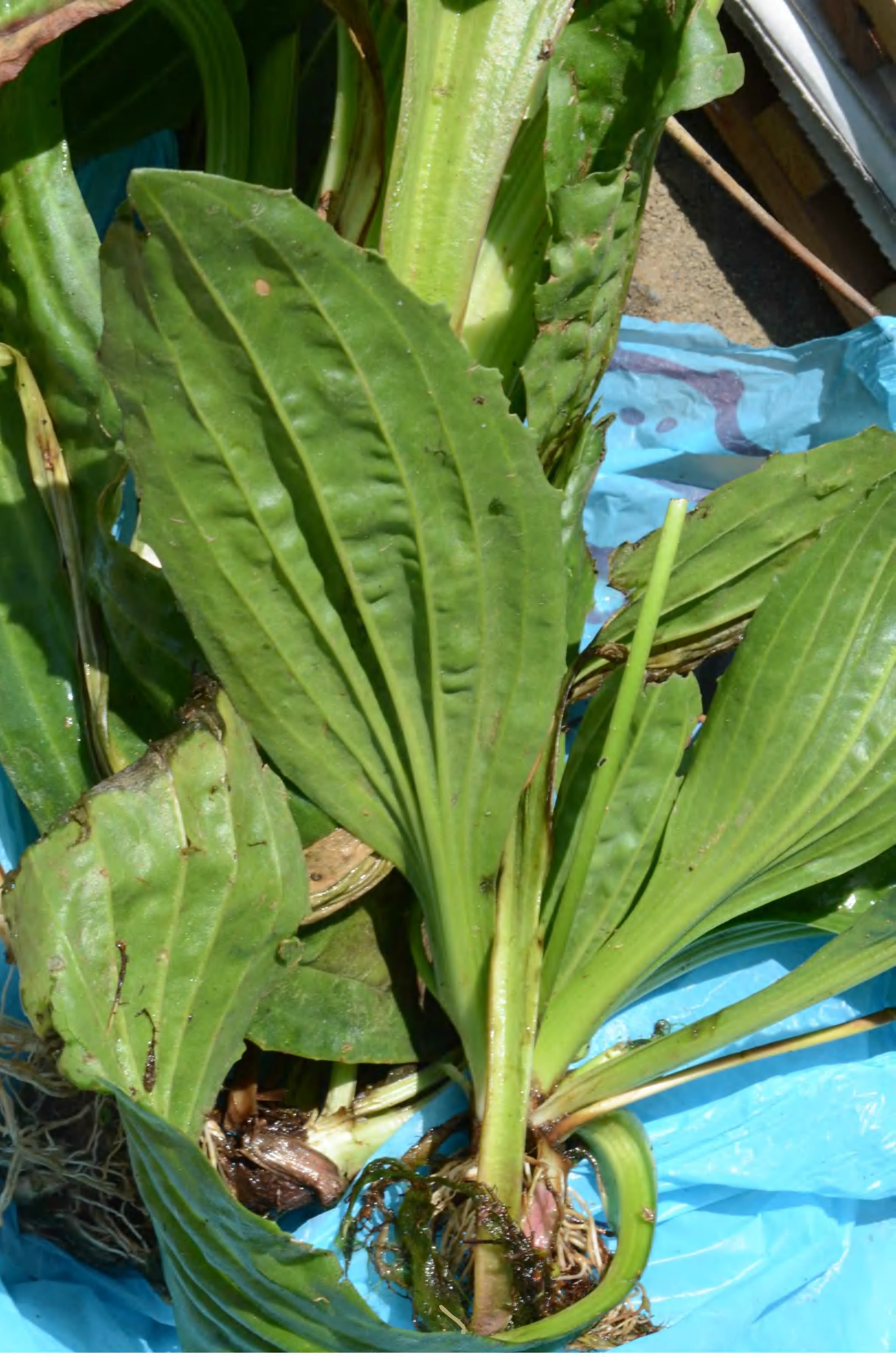

plantago_pachyneura_p_1223_bbb_3581.jpg

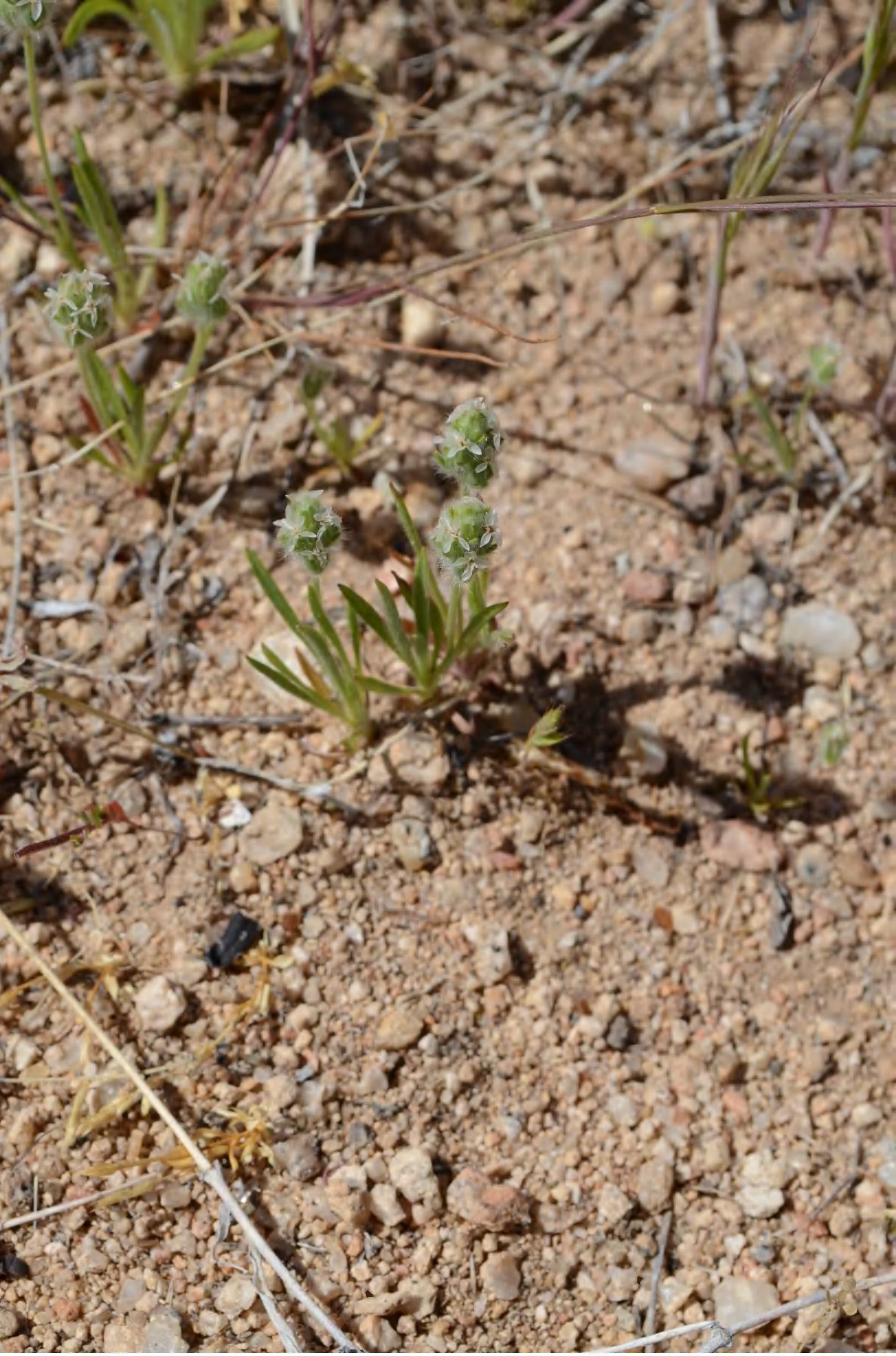

plantago_patagonica_az_dsc_6345.jpg

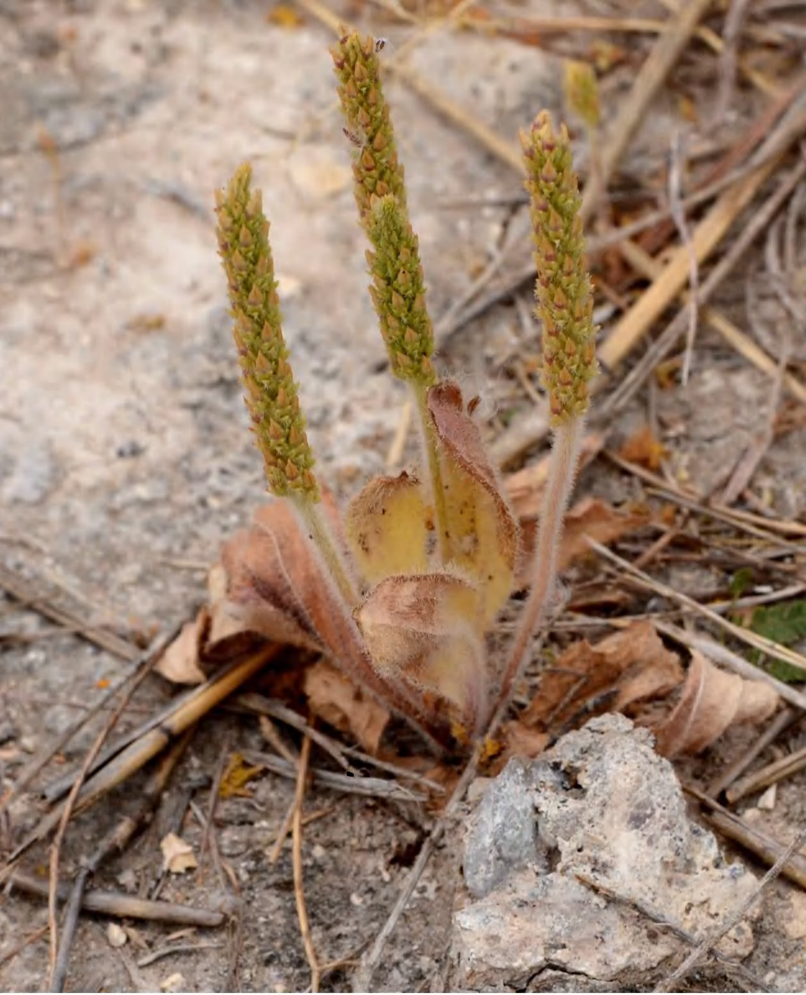

plantago_rhodosperma_p_0740_bbb_7888.jpg
