## Supplementary Tables and Figures for "How to map a plantain: phylogeny of the diverse *Plantagineae* (Lamiales)": 01_plantagineae.pdf

### Plantagineae Dumort.*

2020-05-26

#### Genus 1. Aragoa Kunth

##### **Sectio Ciliatae** Fern.Alonso

1(1). *Aragoa lucidula* S.F.Blake

##### **Sectio Aragoa**

1(2). *Aragoa abietina* Kunth
2(3). *Aragoa abscondita* Fern.Alonso
3(4). *Aragoa castroviejoi* Fern.Alonso
4(5). *Aragoa cleefii* Fern.Alonso
5(6). *Aragoa corrugatifolia* Fern.Alonso
6(7). *Aragoa cundinamarcensis* Fern.Alonso
7(8). *Aragoa cupressina* Kunth
8(9). *Aragoa dugandii* Romero
9(10). *Aragoa funckii* Fern.Alonso
10(11). *Aragoa hammenii* Fern.Alonso
11(12). *Aragoa kogiorum* Romero
12(13). *Aragoa lycopodioides* Benth. ex Oliver
13(14). *Aragoa occidentalis* Pennell
14(15). *Aragoa parviflora* Fern.Alonso & Castrov.
15(16). *Aragoa perez-arbelaeziana* Romero
16(17). *Aragoa picachensis* Fern.Alonso
17(18). *Aragoa romeroi* Fern.Alonso
18(19). *Aragoa tamana* Fern.Alonso
19(20). *Aragoa ×chingacensis* Fern.Alonso[1]
20(21). *Aragoa ×diazii* Fern.Alonso[2]
21(22). *Aragoa ×funzana* Fern.Alonso[3]
22(23). *Aragoa ×jaramilloi* Fern.Alonso[4]

*Non-standard abbreviations used: sed.m. *sedis mutabilis*; stat.m. *status mutabilis*.
[1] *Aragoa abietina* × *Aragoa cundinamarcensis*
[2] *Aragoa corrugatifolia* × *Aragoa cundinamarcensis*
[3] *Aragoa cleefii* × *Aragoa cundinamarcensis*
[4] *Aragoa abietina* × *Aragoa cupressina*

#### Genus 2. LITTORELLA P.J.Bergius

1(24). *Littorella americana* Fernald[5]
2(25). *Littorella australis* Griseb. ex Benth. & Hook.f.[6]
3(26). *Littorella uniflora* (L.) Asch.[7]

#### Genus 3. PLANTAGO L.

##### Subgenus ***Coronopus*** (Lam. & DC.) Rahn

###### Sectio Coronopus

1(27). *Plantago asphodeloides* Svent
2(28). *Plantago coronopus* L.[8]
3(29). *Plantago carnosa* Lam.
4(30). *Plantago crassifolia* Forssk.
5(31). *Plantago crypsoides* Boiss.[9]
6(32). *Plantago macrorhiza* Poir.
7(33). *Plantago serraria* L.[10]
8(34). *Plantago subspathulata* Pilg.

###### Sectio Maritima H.Dietr.

1(35). *Plantago alpina* L.[11]
2(36). *Plantago eocoronopus* Pilg.
3(37). *Plantago maritima* L.[12]
4(38). *Plantago rhizoxylon* Emb.
5(39). *Plantago subulata* L.[13]

##### Subgenus ***Plantago***

###### Sectio Micropsyllium Decne.

1(40). *Plantago elongata* Pursh[14]
2(41). *Plantago heterophylla* Nutt.[15]
3(42). *Plantago minor* Fr.
4(43). *Plantago polysperma* Kar. & Kir.
5(44). *Plantago pusilla* Nutt.

---

[5] *Plantago americana* (Fernald) Rahn

[6] *Plantago araucana* Rahn

[7] *Plantago uniflora* L.

[8] *Plantago aschersonii* Bolle; *Plantago cupanii* Guss.

[9] *Plantago weldenii* Rchnb. sed.m.; *Plantago commutata* Guss. sed.m.; *Plantago sabulosa* Danin & Raus. sed.m.

[10] *Plantago peloritana* Lojac

[11] *Plantago penyalarensis* Pau

[12] *Plantago atlantica* Batt.; *Plantago juncoides* Lam.; *Plantago neumannii* Opiz; *Plantago oliganthos* Roem. & Schult.; *Plantago salsa* Pall.; *Plantago schrenkii* C. Koch.; *Plantago serpentina* All.; *Plantago subpolaris* Andr.

[13] *Plantago algarbiensis* Samp., stat.m.; *Plantago almogravensis* Franco; *Plantago carinata* Schrad. ex Mert. & W.D.J.Koch; *Plantago holosteum* Scop.; *Plantago insularis* (Godr.) Nyman; *Plantago radicata* Hoffmans. & Link

[14] *Plantago bigelovii* A. Gray

[15] *Plantago hybrida* Bart.

6(45). *Plantago tenuiflora* Waldst. & Kit.

**Sectio Eremopsyllium** Pilg.

1(46). *Plantago gentianoides* Sibth. & Sm.
2(47). *Plantago reniformis* Beck

**Sectio Lamprosantha** Decne.

1(48). *Plantago arachnoidea* Schrenk ex Fisch. & C.A.Mey.
2(49). *Plantago canescens* Adams[16]
3(50). *Plantago krascheninnikovii* Ye. V. Serg.
4(51). *Plantago maxima* Juss. ex Jacq.
5(52). *Plantago media* L.[17]
6(53). *Plantago perssonii* Pilg.
7(54). *Plantago schwarzenbergiana* Schur

**Sectio Leptostachys** Decne.

1(55). *Plantago africana* Verdc.
2(56). *Plantago fischeri* Engl.
3(57). *Plantago laxiflora* Decne.
4(58). *Plantago longissima* Decne.[18]
5(59). *Plantago palmata* Hook.f.
6(60). *Plantago remota* Lam.[19]
7(61). *Plantago tanalensis* Baker

**Sectio Mesembrynia** Decne.

1(62). *Plantago alpestris* B.G.Briggs & al.
2(63). *Plantago antarctica* Decne.
3(64). *Plantago aucklandica* Hook.f.
4(65). *Plantago aundensis* P. Royen
5(66). *Plantago bellidioides* Decne.
6(67). *Plantago cladarophylla* B.G.Briggs & al.
7(68). *Plantago cunninghamii* Decne.[20]
8(69). *Plantago daltonii* Decne.
9(70). *Plantago debilis* R.Br.
10(71). *Plantago depauperata* Merr. & Perry
11(72). *Plantago drummondii* Decne.[21]
12(73). *Plantago euana* Hurlim.
13(74). *Plantago euryphylla* B.G.Briggs & al.
14(75). *Plantago exilis* Decne.
15(76). *Plantago gaudichaudii* Barnéoud[22]
16(77). *Plantago glabrata* Hook.f.

[16] *Plantago jurtzevii* Tzvel.
[17] *Plantago urvillei* Opiz, stat.m.; *Plantago brutia* Ten.
[18] *Plantago* zeyheri auct., nom.nud.
[19] *Plantago capensis* Bojer
[20] *Plantago mitchellii* Decne.
[21] *Plantago pritzelii* Pilg.
[22] *Plantago sericophylla* Decne.; *Plantago bakeri* Pilg.

17(78). *Plantago glacialis* B.G.Briggs & al.
18(79). *Plantago gunnii* Hook.f.
19(80). *Plantago hedleyi* Maiden
20(81). *Plantago hispida* R.Br.[23]
21(82). *Plantago lanigera* Hook.f.
22(83). *Plantago montisdicksonii* P. Royen
23(84). *Plantago muelleri* Pilg.
24(85). *Plantago multiscapa* B.G.Briggs
25(86). *Plantago novae-zelandiae* L.B. Moore
26(87). *Plantago obconica* Sykes
27(88). *Plantago palustris* L.R.Fraser & Vickery
28(89). *Plantago papuana* P. Royen
29(90). *Plantago paradoxa* Hook.f.
30(91). *Plantago pentasperma* Hemsl.
31(92). *Plantago picta* Colenso
32(93). *Plantago polita* Craven
33(94). *Plantago raoulii* Decne.
34(95). *Plantago robusta* Roxb., sed.m.
35(96). *Plantago spathulata* Hook.f.
36(97). *Plantago stauntonii* Reichardt
37(98). *Plantago stenophylla* Merr. & L.M.Perry
38(99). *Plantago tasmanica* Hook.f.
39(100). *Plantago triandra* Berggr.[24]
40(101). *Plantago triantha* Spreng.
41(102). *Plantago trichophora* Merr. & L.M.Perry
42(103). *Plantago turrifera* B.G.Briggs & al.
43(104). *Plantago udicola* Meudt & Garn.-Jones
44(105). *Plantago unibracteata* Rahn[25]
45(106). *Plantago varia* R.Br.[26]

**Sectio Plantago**

1(107). *Plantago cornuti* Gouan[27]
2(108). *Plantago griffithii* Decne.[28]
3(109). *Plantago major* L.[29]
4(110). *Plantago japonica* Franch. & Sav., stat.m.[30]
5(111). *Plantago tatarica* Decne.

[23] *Plantago tildeniae* Pilg.
[24] *Plantago masoniae* Cheesem.
[25] *Plantago uniflora* Hook.f., non L.
[26] *Plantago acutiloba* Pilg.; *Plantago struthionis* A.Cunn. ex Decne.
[27] *Plantago exaltata* Hornem.
[28] *Plantago aitchisonii* Pilg.
[29] *Plantago himalaica* Pilg., stat.m.; *Plantago brachyphylla* Edgew. ex Decne.; *Plantago uliginosa* F. W. Schmidt, stat.m.; *Plantago winteri* Wirtg.
[30] *Plantago macronipponica* Yamamoto

**Sectio Heptaneuron** Decne.

1(112). *Plantago cordata* Lam.

**Sectio Carpophorae** Rahn

1(113). *Plantago rigida* Kunth
2(114). *Plantago tubulosa* Decne.

**Sectio Pacifica** Hassemer

1(115). *Plantago anatolica* Tutel & R.R.Mill
2(116). *Plantago asiatica* L.[31]
3(117). *Plantago camtschatica* Link
4(118). *Plantago chihuahuensis* Shipunov sp.nov.
5(119). *Plantago depressa* Willd.
6(120). *Plantago eriopoda* Torr.[32]
7(121). *Plantago hakusanensis* Koidz.
8(122). *Plantago hasskarlii* Decne.
9(123). *Plantago hawaiensis* (A. Gray) Pilg.
10(124). *Plantago incisa* Hassk.[33]
11(125). *Plantago komarovii* Pavlov
12(126). *Plantago pachyphylla* A. Gray[34]
13(127). *Plantago princeps* Cham. & Schltdl.[35]
14(128). *Plantago rapensis* Pilg.
15(129). *Plantago rugelii* Decne.
16(130). *Plantago rupicola* Pilg.
17(131). *Plantago sparsiflora* Michx.
18(132). *Plantago tweedyi* A. Gray

**Sectio Holopsyllium** Pilg.

1(133). *Plantago macrocarpa* Cham. & Schltdl.

**Sectio Virginica** Decne. & Steinh. ex Barnéoud

1(134). *Plantago alismatifolia* Pilg.
2(135). *Plantago argentina* Pilg.
3(136). *Plantago australis* Lam.[36]
4(137). *Plantago barbata* G. Forst[37]

---

[31] *Plantago alata* Nakai; *Plantago cavaleriei* H.Lév.; *Plantago centralis* Pilg.; *Plantago coreana* H.Lév.; *Plantago erosa* Wall.; *Plantago fengdouensis* (Z.E Chao & Yong Wang) Yong Wang & Z.Yu Li, stat.m.; *Plantago formosana* Tateishi & Masam.; *Plantago hostifolia* Nakai & Kitag.; *Plantago nanchuanensis* J.Z. Liu, nom.nud.; *Plantago popovii* Tzvel.; *Plantago sawadai* Yamam.; *Plantago schneideri* Pilg.; *Plantago taquetii* H.Lév.; *Plantago yakushimensis* Masam.; *Plantago yezoensis* Pilg.; *Plantago zhongdainensis* J.Z.Liu, sed.m., nom.nud.

[32] *Plantago shastensis* Greene

[33] *Plantago densiflora* J.Z. Liu, sed.m.

[34] *Plantago muscicola* Pilg.; *Plantago hillebrandii* Pilg.; *Plantago glabrifolia* (Rock) Pilg.; *Plantago melanochrous* Pilg.; *Plantago krajinai* Pilg.; *Plantago grayana* Pilg.

[35] *Plantago longibracteata* (Mann) Tessene, nom.nud.

[36] *Plantago candollei* Raf.; *Plantago deppeana* Vatke, sed.m.; *Plantago hirtella* Kunth; *Plantago schiedeana* Decne.

[37] *Plantago monanthos* D'Urville, p.p.

5(138). *Plantago berroi* Pilg.
6(139). *Plantago bradei* Pilg.
7(140). *Plantago buchtienii* Pilg.
8(141). *Plantago catharinea* Decne.
9(142). *Plantago commersoniana* Decne. & Barnéoud
10(143). *Plantago correae* Rahn
11(144). *Plantago corvensis* Hassemer[38]
12(145). *Plantago cumingiana* Fisch. & C.A.Mey.
13(146). *Plantago cuzcoensis* Shipunov, sp.nov.
14(147). *Plantago dielsiana* Pilg.
15(148). *Plantago fernandezia* Barnéoud
16(149). *Plantago firma* Kunze ex Walp.[39]
17(150). *Plantago floccosa* Decne.
18(151). *Plantago galapagensis* Rahn
19(152). *Plantago guilleminiana* Decne.
20(153). *Plantago hatschbachiana* Hassemer
21(154). *Plantago humboldtiana* Hassemer
22(155). *Plantago jujuyensis* Rahn
23(156). *Plantago moorei* Rahn
24(157). *Plantago myosuros* Lam.
25(158). *Plantago napiformis* (Rahn) Hassemer
26(159). *Plantago orbignyana* Decne.[40]
27(160). *Plantago oreades* Decne.
28(161). *Plantago pachyneura* Steud.
29(162). *Plantago penantha* Griseb.
30(163). *Plantago pretoana* (Rahn) Hassemer
31(164). *Plantago pulvinata* Speg.
32(165). *Plantago pyrophila* Villarroel & J. R. I. Wood
33(166). *Plantago rahniana* Hassemer & R. Trevis.
34(167). *Plantago rhodosperma* Decne.
35(168). *Plantago sempervivoides* Dusen
36(169). *Plantago subnuda* Pilg., stat.m.
37(170). *Plantago tehuelcha* Speg.
38(171). *Plantago tenuipala* (Rahn) Rahn
39(172). *Plantago tomentosa* Lam.[41]
40(173). *Plantago trinitatis* Rahn
41(174). *Plantago truncata* Cham. & Schltdl.
42(175). *Plantago turficola* Rahn
43(176). *Plantago uniglumis* Wallr. ex Walp.[42]
44(177). *Plantago veadeirensis* Hassemer

[38] *Plantago aparadensis* D. Falkenberg
[39] *Plantago skottsbergii* Pilg.
[40] *Plantago hartwegii* Decne.
[41] *Plantago paralias* Decne.
[42] *Plantago monanthos* D'Urville, p.p.

45(178). *Plantago ventanensis* Pilg.
46(179). *Plantago venturii* Pilg.
47(180). *Plantago virginica* L.
48(181). *Plantago weddelliana* Decne.

**Subgenus *Bougueria*** (Decne.) Rahn

49(182). *Plantago nubicola* (Decne.) Rahn

**Subgenus *Albicans*** Rahn

**Sectio Hymenopsyllium** Pilg.

1(183). *Plantago bellardii* All.[43]
2(184). *Plantago benisnassenii* Romo & al.
3(185). *Plantago ciliata* Desf.
4(186). *Plantago cretica* L.
5(187). *Plantago cyrenaica* E.A.Durand & Barratte

**Sectio Montana** Barnéoud

1(188). *Plantago atrata* Hoppe[44]
2(189). *Plantago cafra* Decne.[45]
3(190). *Plantago monosperma* Pourr.
4(191). *Plantago nivalis* Boiss

**Sectio Lancifolia** Barnéoud

1(192). *Plantago altissima* L.
2(193). *Plantago argentea* Chaix[46]
3(194). *Plantago pilgeriana* Hassemer[47]
4(195). *Plantago lagopus* L.
5(196). *Plantago lanceolata* L.[48]
6(197). *Plantago malato-belizii* Lawalree
7(198). *Plantago loeflingii* L.[49]

**Sectio Albicans** Barnéoud

1(199). *Plantago akkensis* Coss.
2(200). *Plantago albicans* L.
3(201). *Plantago amplexicaulis* Cav.[50]
4(202). *Plantago annua* Ryding
5(203). *Plantago baltistanica* Hartmann
6(204). *Plantago boissieri* Hausskn. & Bornm.
7(205). *Plantago cylindrica* Forssk.

---

[43] *Plantago bellardi* L.
[44] *Plantago discolor* Gand.; *Plantago fuscescens* Jord.; *Plantago saxatilis* Bieb.
[45] *Plantago capillaris* E. Mey. ex Decne.
[46] *Plantago serpentinicola* Reich.
[47] *Plantago lacustris* Maire, nom.illeg.; *Plantago maireana* Hassemer
[48] *Plantago dubia* L.; *Plantago leiopetala* Lowe
[49] *Plantago scabrifolia* Thieb., sed.m.
[50] *Plantago bauphula* Edgew.

8(206). *Plantago lachnantha* Bunge[51]
9(207). *Plantago lagocephala* Bunge
10(208). *Plantago minuta* Pall.[52]
11(209). *Plantago notata* Lag.
12(210). *Plantago orzuiensis* Mohsenz. & al.
13(211). *Plantago ovata* Forssk.[53]
14(212). *Plantago psammophila* Agnew & Chal.-Kabi
15(213). *Plantago sharifii* Rech.f. & Esfand., sed.m.
16(214). *Plantago stocksii* Boiss.
17(215). *Plantago tunetana* Murb.

**Sectio Gnaphaloides** Barnéoud

1(216). *Plantago argyrea* Morris
2(217). *Plantago aristata* Michx.
3(218). *Plantago bismarckii* Niederl.
4(219). *Plantago brasiliensis* Sims
5(220). *Plantago densa* Pilg.
6(221). *Plantago erecta* Morris[54]
7(222). *Plantago grandiflora* Meyen[55]
8(223). *Plantago helleri* Small
9(224). *Plantago hispidula* Ruiz & Pav.
10(225). *Plantago hookeriana* Decne.
11(226). *Plantago johnstonii* Pilg.
12(227). *Plantago lamprophylla* Pilg.
13(228). *Plantago limensis* Pers.[56]
14(229). *Plantago linearis* Kunth
15(230). *Plantago litorea* Phil.
16(231). *Plantago lundborgii* Sparre.
17(232). *Plantago nebularis* Hassemer
18(233). *Plantago nivea* Kunth
19(234). *Plantago patagonica* Jacq.[57]
20(235). *Plantago rancaguae* Steud.
21(236). *Plantago sericea* Ruiz & Pav.[58]
22(237). *Plantago tandilensis* Pilg.
23(238). *Plantago tolucensis* Pilg.
24(239). *Plantago wrightiana* Decne.
25(240). *Plantago zoellneriana* Hassemer

[51] *Plantago evacina* Boiss.
[52] *Plantago lessingii* Fisch. & C.A. Mey.
[53] *Plantago fastigiata* Morris
[54] *Plantago speciosa* Morris
[55] *Plantago macrantha* Decne.
[56] *Plantago tacnensis* Pilg.
[57] *Plantago purshii* Roem. ex Schult.; *Plantago spinulosa* Decne.
[58] *Plantago argyrophylla* Decne.; *Plantago macbridei* Pilg., sed.m.; *Plantago nubigena* Kunth; *Plantago perreymondii* Barn.; *Plantago polyclada* Pilg.

**Subgenus *Psyllium*** (Mill.) Harms & Reiche

**Sectio Arborescens** Shipunov sect.nov.

1(241). *Plantago arborescens* Poir.[59]
2(242). *Plantago asperrima* Gand. ex Hervier[60]
3(243). *Plantago famarae* Svent
4(244). *Plantago mauritanica* Boiss. & Reut.
5(245). *Plantago sempervirens* Crantz[61]
6(246). *Plantago sinaica* Barnéoud[62]
7(247). *Plantago webbii* Barnéoud

**Sectio Psyllium** Juss.

1(248). *Plantago afra* L.[63]
2(249). *Plantago arenaria* Waldst. & Kit.[64]
3(250). *Plantago chamaepsyllium* Zohary
4(251). *Plantago euphratica* Barnéoud
5(252). *Plantago exigua* Murray[65]
6(253). *Plantago maris-mortui* Eig
7(254). *Plantago phaeostoma* Boiss. & Heldr.
8(255). *Plantago squarrosa* Murray[66]

---

[59] *Plantago costae* Menezes; *Plantago maderensis* Decne.

[60] *Plantago assoana* Senn.

[61] *Plantago suffruticosa* Lam.

[62] *Plantago arabica* Boiss.

[63] *Plantago cynops* L. 1753, non L. 1762, nom.ambig. *Plantago psyllium* L. 1762, non L. 1753, nom.ambig.; *Plantago squalida* Salisb.; *Plantago rugosa* Hochst. ex Steud.

[64] *Plantago indica* L. 1759, nom.prop.rej.; *Plantago psyllium* L. 1753, nom.ambig.; *Plantago scabra* Moench, nom.illeg.

[65] *Plantago pumila* L.f.

[66] *Plantago sarcophylla* Boiss. ex Decne.
